# How total mRNA influences cell growth

**DOI:** 10.1101/2023.03.17.533181

**Authors:** Ludovico Calabrese, Luca Ciandrini, Marco Cosentino Lagomarsino

## Abstract

Experimental observations tracing back to the 1960s imply that ribosome quantities play a prominent role in determining a cell’s growth. Nevertheless, in biologically relevant scenarios, growth can also be influenced by the levels of mRNA and RNA polymerase. Here, we construct a quantitative model of biosynthesis providing testable scenarios for these situations. The model explores a theoretically-motivated regime where RNA polymerases compete for genes and ribosomes for transcripts, and gives general expressions relating growth rate, mRNA concentrations, ribosome and RNA polymerase levels. On general grounds, the model predicts how the fraction of ribosomes in the proteome depends on total mRNA concentration, and inspects an underexplored regime in which the trade-off between transcript levels and ribosome abundances sets the cellular growth rate. In particular, we show that the model predicts and clarifies three important experimental observations, in budding yeast and *E. coli* bacteria: (i) that the growth-rate cost of unneeded protein expression can be affected by mRNA levels, (ii) that resource optimization leads to decreasing trends in mRNA levels at slow growth, and (iii) that ribosome allocation may increase, stay constant, or decrease, in response to transcription-inhibiting antibiotics. Since the data indicate that a regime of joint limitation may apply in physiological conditions and not only to perturbations, we speculate that this regime is likely self-imposed.

## Introduction

Understanding cell growth is a classic and central question in biology (1; 2; 3). A striking progress of the recent years is the development of mathematical theories that explain quantitative relationships between global parameters of biosynthesis and growth rate, initially found classically for bacteria (4; 5; 6). Such growth laws can be understood as “resource allocation” constraints, as they reflect the presence of a finite pool of resources allocated to the production of different protein classes. For instance, allocating ribosomes to production of one protein class implies withdrawing ribosomes from the production of another class. Growth laws can also be interpreted as consequences of the principles of flux balance and mass conservation. These constraints may undergo alterations in response to varying limitations (conditions), as cells align catabolic and biosynthesis fluxes to achieve steady growth (7; 8; 9; 10). Crucially, this approach has proved important to generate predictive models, for example, of the global regulation of gene expression across growth conditions (11; 12; 13), the growth response to different classes of translation-targeting antibiotics (7; 14) and the cost of producing unnecessary proteins (7). While most of the recent work in this area has been performed on *E. coli* (12; 15; 8; 11; 16; 17; 18), there are strong indications that different quantitative relationships hold across bacteria, in budding yeast and across lower eukaryotes (7; 11; 19; 20; 21; 22; 23). This suggests the presence of strong unifying principles (likely reflecting universal aspects due to resource-allocation trade-offs) in biosynthesis across kingdoms, despite the profound architectural and regulatory differences between organisms (24; 22; 25; 26; 27).

A key aspect for sustaining growth is autocatalysis from ribosome self replication, which is also a primary ingredient of growth-laws theories (9; 2; 25; 28). More broadly, growth laws originate from the presence of a finite pool of cellular resources needed for biosynthesis. As such, their specific form depends on which factors are limiting for biosynthesis, and indeed many experiments show that in most conditions ribosomes are a major limiting factor for growth. However, there are both physiological and perturbed situations where this ceases to be the case. In particular, a growing body of evidence suggests that in several circumstances transcription, RNA polymerase and mRNA levels become relevant for setting growth rate (29; 30; 31; 25; 32; 33; 34).

Therefore, a critical next question for theory is how to produce theoretical frameworks that evolve the ribosome-centered view, preserving its transparency, while also accounting for the experimental observations that call for a more comprehensive description of limitations in biosynthesis, and in particular for the general observation that gene transcription plays a major role in setting the growth rate (25; 30).

Here, our goal is to provide precisely this extended framework. In order to introduce our approach, let us revise the body of the experimental evidence that inspires it, and the previous theories the form the basis of our work. Experimentally, we started from several lines of evidence. The first is the production of unnecessary proteins, which leads to a reduction in growth rate (a “growth cost”). The established view (based on *E. coli* data and ribosome-centric models), is that the cost of unneeded protein expression comes from the mass fraction of the proteome occupied by this unneeded protein (7). In *S. cerevisiae*, a study by Kafri and collegues (32) has shown that in some conditions the growth cost of protein overexpression also comes from a transcriptional burden (but we lack a quantitative framework that captures this cost). Turning to drugs perturbing the global transcription rate in *E. coli*, several studies (35; 7; 36) have shown that the growth rate decreases smoothly under exposure to rifampicin, a well-known antibiotic that targets transcription elongation by binding to RNA polymerase. Intriguingly, Scott and coworkers (7) also found a non obvious (and yet unexplained) nutrient-dependent ribosome re-allocation under rifampicin treatment, which may suggest that cells possess specific mechanisms in place to deal with limited transcriptional capacity. From a theoretical perspective, two recent studies have included the central dogma into a theory of biosynthesis. Lin and Amir (33) have used the framework to explore the limits of full saturation, for example when ribosomes fully saturate mRNAs. Roy and coworkers (9) defined a more complex model of growth stemming from a universal ‘autocatalytic network’, which includes several layers of biosynthesis. Their model identifies several limiting regimes beyond translation-limited growth, including transcription. Importantly, both studies describe limitations by taking a minimum of the effective numbers of potentially limiting species, a classic assumption termed “Liebig’s Law of the Minimum” in an Ecology context (37; 38). However, more general descriptions with joint limitations are possible (39; 37; 40).

By contrast, theoretical considerations on ribosome recruitment and translation lead us to analyze specifically the situation of competition for transcripts during translation initiation, i.e. the process involving the formation of the ribosome-mRNA complex. Additionally, both the frameworks in (33; 9) do not consider the problem of allocation of resources optimizing growth rate. This principle was shown to underline multiple growth laws (10; 41; 42).

It should be noted that while growth rate optimality is generally a very good guiding principle for growth laws (10; 41; 42), it is also clear to us that on biological grounds, things are more complex (43). Indeed, growth rate is not the sole quantity that allocation of resources such as ribosomes and catabolic enzymes allocation should optimize, and that other quantities should be relevant such as ability to respond (or to hedge their bets) in changing environments, to filter out “false-alarms” due to noise, to resist famine conditions (44). Additionally, mechanistic regulation might be trying to achieve growth-rate optimization heuristically, and sub-optimal behavior may be determined by architectural limits of cellular regulation. Accordingly, empirical regulatory systems were also found to be sub-optimal in terms of growth rate (45).

Our framework integrates biosynthesis and the central dogma at both the transcriptional and translational level along the lines of the studies by Lin and Amir, and Roy and coworkers (33; 9), but also motivates and investigates unexplored co-limitation regimes and adds a description of growth laws with growth-rate optimization (10; 41). We analyze this model in both growth-optimized and non-optimized conditions. As we will argue in the following, this combination of ingredients results in a theory able to predict the key experimental findings described above. More widely, it describes the dependency of growth rate on both transcription and translation, through allocation of resources of the biosynthetic machinery, mainly RNA polymerase and ribosomes. Crucially, we show that mRNA levels can affect growth rate well outside of the saturation regime, in a regime where ribosomes within a finite pool compete for transcripts. In this regime the ribosome-mRNA complex formation limits biosynthesis: ribosome autocatalysis is still the engine of cell growth, but its throttle are mRNA levels. We surmise that the cell voluntarily imposes this regime, as the metabolic expenses associated with mRNA-related processes are relatively small.

## Results

### A minimal biosynthesis model including transcription

Our first step is to propose a general framework (Figure 1A) that links cellular growth physiology with two layers of the central dogma: transcription (for which we will use the suffix TX in our notation) and translation (suffix TL). The equations describing the expression of different classes of genes representing distinct coarse-grained biological functions (indexed by *i*) are similar to ref. (33). Each class is populated by *g*_*i*_ gene copies. Transcription synthesizes *m*_*i*_ mRNA copies of each gene class, which are then translated into proteins with abundances *P*_*i*_. The abundance of transcripts *m*_*i*_ and proteins *P*_*i*_ belonging to gene type *i* evolves according to the following equations,

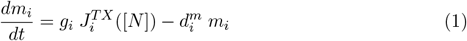

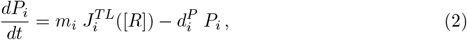

where the fluxes 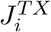 and 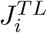 are the RNA polymerase flux per gene (corresponding to the amount of mRNA produced per unit time per gene) and the ribosome flux per mRNA (protein produced per unit time per mRNA). The previous equations consider the general case in which transcripts are degraded with rate 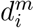 and proteins with rate 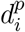. Regardless of their particular form, the transcriptional and translational fluxes 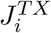 and 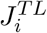 are functions of the total RNA polymerase and ribosome concentrations [*N*] and [*R*] respectively. Figure 1B graphically summarises the main elements of the model.

**Fig 1.**
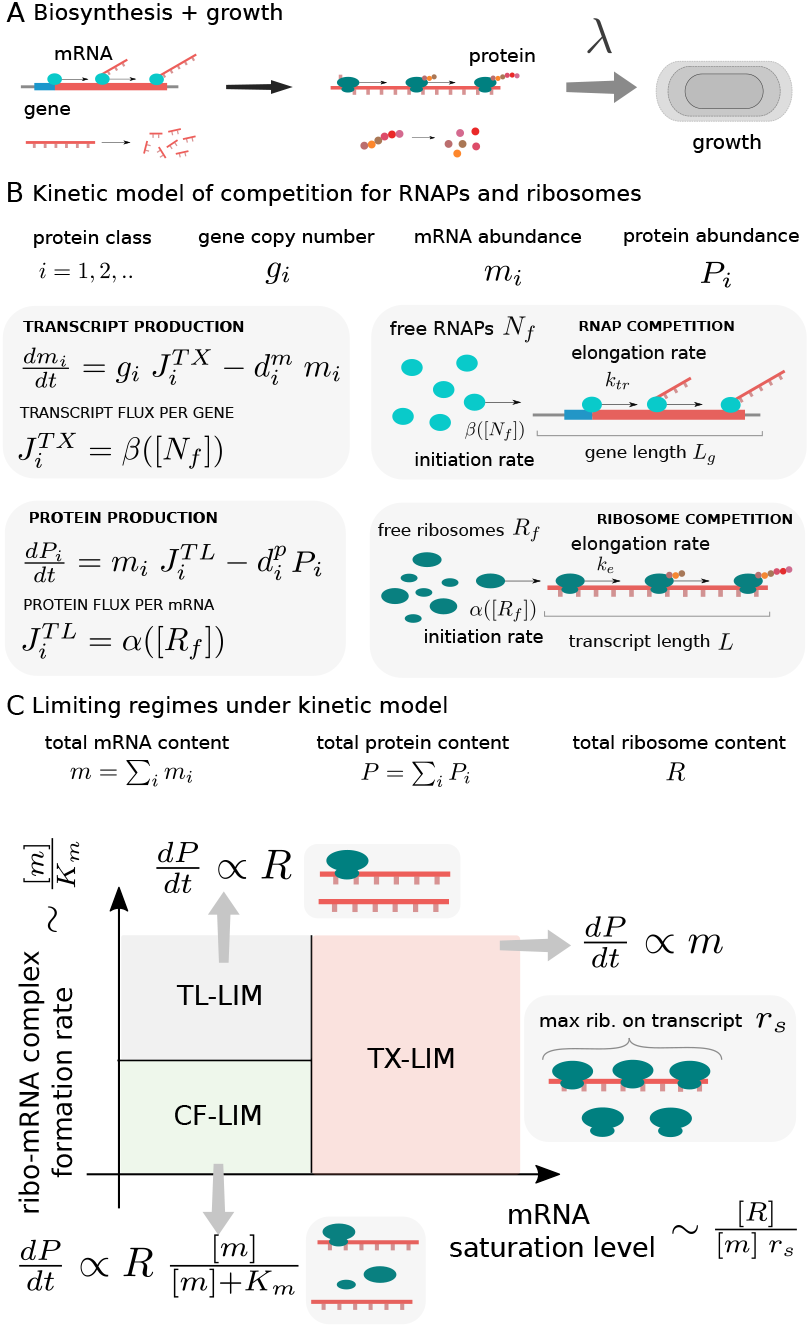
Illustration of the cell growth model including resource allocation and transcription. (A) The model predicts cellular growth rates taking into account translation and transcription. (B) In the model, transcription and translation are treated on the same footing (transcript and protein production boxes). The core of the model is encoded in the transcript flux per gene and the protein flux per gene. These fluxes are determined by the respective initiation and elongation rates, which describe the recruitment of RNA polymerases and ribosomes. In the model, different gene types compete for recruiting free RNA polymerases (“RNAP competition” box) and transcripts compete for free ribosomes (ribosome competition box). Consequently, the synthesis of all proteins is coupled through the availability of RNA polymerases and ribosomes. (C) Qualitative sketch of the different expected regimes of growth. Our model focuses on the regime in which growth is driven by both ribosome levels and mRNA concentration (green region). In this regime the formation of the ribosome-mRNA complex is the limiting step (complex-formation limiting regime, CF-LIM). Instead, if translation or transcription alone are the limiting step, protein production depends only on either the number of ribosomes (translation-limited regime TL-LIM) or the number of mRNAs (transcription-limited regime TX-LIM), according to the level of mRNA saturation by ribosomes (see grey and red regions).

The reservoirs of RNA polymerase and ribosomes couple the expression of all gene types. Such ‘global coupling’ is a crucial element of a class of biosynthesis models able to link gene expression and growth physiology (46; 33; 9). Note that the expression of proteins constituting RNA polymerase and ribosomes will also need to obey Eqs.[1] and [2] above. This gives rise to autocatalytic cycles able to sustain exponential growth. Specifically, Roy and coworkers (9) have shown how one can derive growth laws from autocatalitic cycles involving key players such as ribosomes, RNA polymerase, tRNA, etc. We show in SI Appendix how this is possible within our model.

In typical applications, e.g. for bacteria, mRNA degradation rates are considered to be high (33; 30), and protein degradation rates are neglected (11). Although these approximations should be taken with caution (32; 47), they are reasonable in regimes where the time scales of growth and protein degradation are separated (47). Thus, for the sake of simplicity, in the following we will neglect the role of protein degradation. Instead, mRNA degradation rates are fast (typical half lives of 5-20 minutes, depending on the organism), and will play a crucial role in the following. We will consider homogeneous transcript degradation rates, 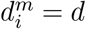.

The total amounts of mRNAs and proteins, which will be the main quantities of interest for us, are given by the sum of transcripts and proteins over all the gene types, *m* := Σ_*i*_ *m*_*i*_, and *P* := Σ_*i*_ *P*_*i*_. The model produces steady-state exponential (“balanced”) growth (see SI Appendix), where extensive quantities (defined as the quantities that increase linearly with total biomass) all grow exponentially with the same rate.

A suitable parameter to characterise cellular physiology is thus the protein-specific growth rate, defined as

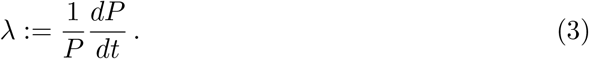

We have singled out conceptually protein production over mRNA production because measurements of total protein content are much more prevalent in the literature. At the same time, the definition above becomes the growth rate of every molecular species at the steady state, including mRNAs (7). Therefore, in balanced growth regimes (to which we will restrict ourselves here) such rate still fully characterizes the growth process.

Finally, while we are very aware that organism-specific features are often very important, we will keep our framework organism-agnostic, aiming to describe a general economy of cellular trade-offs (at the level of protein synthesis) with a reduced set of parameters.

### The model captures a regime where mRNA-ribosome complex formation limits translation

Differently from previous studies that identify limitation regimes by identifying a single molecular species in the shortest supply, we explore a scenario characterized by potential co-limitations (37; 38; 39; 40). We can use a toy-model argument based on chemical kinetics (48) to demonstrate the existence of different limiting regimes in protein production (Fig. 1C, see below and SI Appendix, sec. 3). We consider two chemical species *A* and *B* that form a complex *AB* with a binding constant *k*_*CF*_ following mass action. The complex *AB* produces *P* with a rate *k*_*p*_, and spontaneously dissociates into *A* and *B* simultaneously. We assume that all reactions are irreversible and far-from-equilibrium.

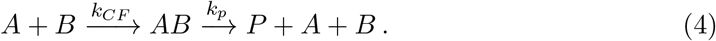

The rate equations corresponding to this process following mass action kinetic can lead to

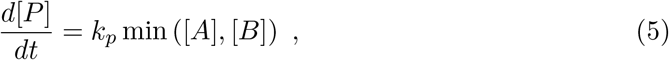

describing the regimes [*A*] ≫ *K*_*AB*_ or [*B*] ≫ *K*_*AB*_, with 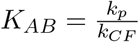, but also to

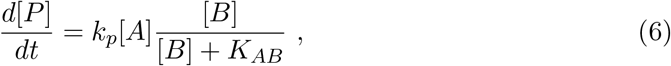

in case [*A*] ≪ *K*_*AB*_ and [*B*] ≈ *K*_*AB*_. If we now interpret [*A*], [*B*] and [*P*] in our toy model as representative, respectively, of ribosomes [*R*], mRNA [*m*] and proteins [*P*], we can map conceptually this toy model to protein production. Consequently, there can be two limiting regimes as in Eq.[5], which we name TX-LIM and TL-LIM and in which the production rate depends uniquely on the concentration of mRNAs or ribosomes, respectively (Fig. 1C). The remaining scenarios correspond to limiting regimes where the rate of product production is influenced by both [*m*] and [*R*]. In these cases, the competition between mRNA and ribosomes is crucial, and the system is constrained by the formation of the complex mRNA-ribosome (CF-LIM). It is noteworthy that, starting from the CF-LIM regime, the translation-limited regime can be approached asymptotically as the total mRNA concentration [*m*] increases.

The context of central-dogma biosynthesis includes multiple complications. In particular, the symmetry between *A* and *B* is lost due to the fact that ribosomes and mRNA are not on equal footing. However, we believe that this toy model provides the correct intuition for the presence of different limiting regimes comprising in particular two regimes limited by products (transcription, TX-LIM and translation, TL-LIM), and a complex-formation limiting regime (CF-LIM). A similar kinetic argument was used to understand the TX-LIM regime in ref. (33) (see Appendix of that study).

### mRNA-ribosome complex formation impacts growth rate

To evaluate the impact of the transcription machinery on cellular physiology we have derived a general expression that relates the exponential growth rate *λ*, the ribosome allocation, and the total mRNA concentration [*m*], and is valid in the particular regime of “limiting complex formation” (CF-LIM). Both *a priori* arguments and comparison of model predictions with data lead us to hypothesize that this regime is relevant in many experimental situations. Such an expression takes the particularly transparent form under a few further simplifying assumptions

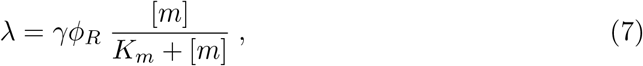

where *γ* is an effective translation rate, which can be interpreted as the inverse of the time needed to assemble the amino acids constituting a ribosome, *ϕ*_*R*_ = *P*_*R*_*/P* is the ribosomal protein fraction. We note that for simplicity of notation we use *ϕ*_*X*_ to denote proteome *number* fractions, and not mass fractions as done in previous literature (9; 30), but the two quantities can be converted into each other, using the masses of the proteins belonging to each sector.

The constant *K*_*m*_ is an effective mRNA concentration scale set by the ratio of the translation initiation and elongation rates. It is important to note that Eq. [7] is not a mere assumption of Michaelis-Menten dilute-substrate kinetics, but rather derives from the biosynthesis fluxes of transcription and translation, taking into account initiation-limited processes and a finite amount of ribosomes and RNA polymerases. In order to understand this crucial point, we discuss the steps leading to the derivation of Eq. [7] from a model of translation initiation and elongation (49), and clarify our assumptions. We first assume a translation initiation rate *α* proportional to the concentration [*R*_*f*_] of free, unbound ribosomes. Mathematically, this can be written as *α* = *α*_0_[*R*_*f*_], where *α*_0_ is a rate constant representing the affinity between ribosomes and transcripts. We combined such model of initiation with a model of protein production (14), which allows us to re-write the free ribosome concentration in terms of the total ribosome concentration [*R*], total mRNA concentration [*m*] and translation elongation rate *k*_*tl*_. Specifically, for one transcript the typical steady-state number of bound ribosomes 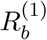 is given by the balance of the initiation flux *αP*_*in*_ with the elongation flux *k*_*tl*_*ρ*^(1)^*P*_*el*_, where 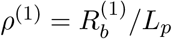 is the ribosome linear density along a single transcript, i.e. the ratio between the ribosomes bound on a transcript 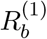 and the average mRNA length *L*_*p*_. *P*_*in*_ is the probability of a successful ribosome initiation attempt (which can fail because of ribosome interference), and *P*_*el*_ is a probability of a successful ribosome step from codon to codon (which can be impacted by ribosome interference). Under the assumption that *P*_*in*_ ≃ *P*_*el*_ (i.e. that ribosome interference is comparable along the transcript), the balance of initiation and elongation fluxes leads to the relation between the free ribosome concentration and the steady-state number of ribosomes on a single mRNA:

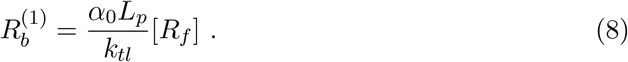

Now, since 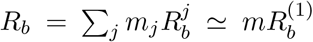 one has for the total concentration of bound ribosomes

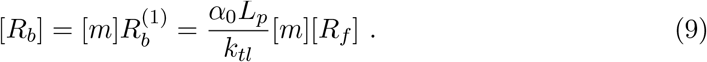

Since [*R*] = [*R*_*f*_] + [*R*_*b*_] we obtain immediately

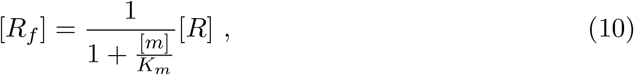

where 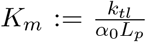 and [*m*]*/K*_*m*_ describes the ratio between the typical time needed by a ribosome to interact with a transcript, 1/(*α*_0_[*m*]), and the typical time needed to complete elongation and dissociate *L*_*p*_*/k*_*tl*_ (see also Methods and SI Appendix). Thus, in the regime where the translation flux *J*^*T L*^ is proportional to the initiation rate, i.e. assuming 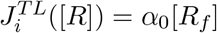, we obtain Eq. [7] from the statement that *dP/dt* = *mJ*^*T L*^. Note that the situation is different from the toy model as it locates the system in a regime where the role of *m* and *R* in the production of *P* is not symmetric. Hence, as anticipated, the Michaelis-Menten-like factor emerges naturally from a kinetic model of translation where ribosomes compete for mRNAs and initiation is limiting (see also Materials and Methods and SI Appendix).

We note that the initiation-limited regime investigate here is a particular one in one-dimensional traffic models (49; 50; 51; 52). It applies for sufficiently low density of RNA polymerases on genes, and of ribosomes on transcripts, as might be the case in physiological conditions (52; 53; 54; 30). Among the other simplifying assumptions used to derive Eq. [7], it is worth noting that we assumed that all genes have identical properties (transcript length and degradation rate, promoter and ribosome binding site strengths, etc.). While relaxing this assumption can lead to interesting predictions, such as non-linear growth laws for specific mRNAs (55), this assumption should be sufficient to capture the overall behavior of the system, and specifically the effect of total mRNA. Moreover, we have neglected gene expression regulation at the translational level (so that genes are differentially expressed only through differences at the transcriptional level (30)). Finally, we have chosen not to describe tRNA and rRNA levels, assuming that they are matched by dedicated regulatory systems (56; 23; 28; 15). This regulation should be different in eukaryotes, where a feedback control system is absent, entirely different RNA polymerase complexes are required for mRNA, tRNA and rRNA production, and control over tRNA synthesis appears to be less stringent (57) (see the SI Appendix for model variants including both rRNA and mRNA).

Similarly to previous frameworks (7; 11; 10; 9), Eq. [7] linearly relates the growth rate to the ribosomal protein fraction *ϕ*_*R*_ with a proportionality factor associated with the translation capacity *γ*. However, differently from other frameworks, Eq. [7] also includes the factor [*m*]/(*K*_*m*_ +[*m*]) reflecting the competition of transcripts for ribosomes. This term comes from the assumption of a regime where the formation of the ribosome-transcript complex is the limiting step of the process (Fig. 1C). The competition between transcripts and ribosomes, and thus the dependence of growth rate on both mRNA and ribosome levels, becomes relevant when [*m*] ≈ *K*_*m*_ (see SI Appendix).

Our hypothesis (motivated by the experimental evidence on both budding yeast and *E. coli*) is that the co-limited CF-LIM regime illustrated in Fig. 1C may be relevant in different situations. The [*m*]-dependent factor of Eq. [7] describes the non-negligible competition of mRNAs for ribosomes. If mRNAs are abundant, they strongly compete for ribosomes, which become the main limiting factor. In this situation the ratio *K*_*m*_/[*m*] becomes sufficiently small, and the growth rate *λ* depends on the ribosomal fraction *ϕ*_*R*_ only. This is the conventional translation-limited regime (TL-LIM in Fig. 1C), in which transcripts are abundant, [*m*] ≫ *K*_*m*_. At the other extreme, if transcript abundance is low and ribosomes are in excess, our framework predicts that protein production is proportional to both the amount of mRNA [*m*] and *ϕ*_*R*_ (see equation [7]). Thus, when [*m*] ≪ *K*_*m*_, *λ* ∝ *ϕ*_*R*_[*m*] and the formation of the ribosome/transcript complex still limits the protein synthesis process in this regime (CF-LIM). Contrary to the saturated transcript regime explored in previous studies (33; 9) where protein production is proportional to only transcript abundances (TX-LIM in Fig. 1C), our framework predicts a new regime of growth driven by mRNA/ribosome complex formation where growth rate depends on both ribosome and transcript concentrations.

A back-of-the-envelope estimate (see Materials and Methods) supports our hypothesis. We get *K*_*m*_ ≈ 0.05 − 0.15*μM* for *E. coli* and *K*_*m*_ ≈ 0.25 − 0.5*μM* for *S. cerevisiae*. This range of values is roughly an order of magnitude smaller than the typical mRNA concentration at fast growth (30), placing ourself in between the translation limited (TL-LIM) and the strong complex-limited (CF-LIM) competition regimes. This suggests that small perturbations of mRNA levels do not impact growth, but a ten-fold reduction of mRNA levels compared to fast growth could significantly impact the growth rate. While such perturbations could be considered quite strong, mRNA levels can indeed span an order of magnitude in *E. coli*, and they can reach [*m*] ≈ *K*_*m*_ physiologically at slow growth (30). We will see in the following that the CF-LIM regime can explain key experimental trends, further supporting our hypothesis.

Notably, the predictions of a CF-LIM scenario are significantly different from those of a transcription-limited growth (TX-LIM) scenario (SI Appendix, Sec. S4 and Fig. S1). The differences between the two regimes are primarily related to how growth rate relates to mRNA and ribosome levels. Crucially, ribosomes are not a limiting factor in the TX-LIM regime, while in the CF-LIM regime increasing ribosome levels can result in faster growth. These distinctions lead to distinct responses to various growth perturbations, which will be discussed below.

All the above arguments lead us to focus (differently from previous studies) in particular on the CF-LIM regime, where the association mRNA/ribosome becomes limiting. For simplicity (and testability of our model), we have considered a theory where ribosome-mRNA binding (competition) is the only determinant of ribosome activity. While a full description of growth limitations beyond the initiation-limited assumption of the biosynthesis fluxes is beyond the scope of this work, SI Appendix sec. 3 provides further arguments regarding the wider phase space of growth-limiting regimes and the relations of our assumptions with other studies. We are also aware that other contributions, including ribosome sequestration or hibernation, degradation, variations in translation rates at slow growth, drugs perturbing transcription and translation etc. can contribute to Eq. [7]. This is particularly expected at slow growth (58; 15; 47), where consequently Eq. [7] is less precise. The following subsection shows how the relation between fraction of ribosomes in the proteome and growth rate depends on RNA polymerase allocation through a factor that is set by the ratio of the translation-initiation rate to the translation-elongation rate.

### mRNA availability couples the translational and transcriptional machineries

Equation [7] shows that our framework relates growth and ribosome allocation taking into account transcript levels. It is possible to probe the interdependence between the translational and transcriptional machineries by decoupling the contributions of ribosomes and RNA polymerases. Indeed, the model connects mRNA concentration with RNA-polymerase proteins and with key parameters related to mRNA production and degradation.

For a typical gene of length *L*_*g*_ being transcribed at a rate *k*_*tx*_ (measured in nucleotides per unit time), we can write

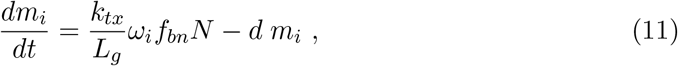

where 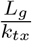 is the typical time needed to transcribe a gene, *ω*_*i*_ is the fraction of RNA polymerases transcribing genes of sector *i* and *f*_*bn*_ the fraction of RNA polymerases that are actively engaged in transcription. The expressions for *ω*_*i*_ and *f*_*bn*_ depend on the binding constants 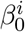 (promoter strength) and the gene copy number *g*_*i*_ (see SI Appendix). Thus, the parameter *f*_*bn*_ is the analogous of the factor [*m*]/(*K*_*m*_ + [*m*]) of Eq. [7], which describes the fraction of ribosomes bound to transcripts and actively translating. However, the expression of *f*_*bn*_ is lengthy and not particularly transparent, so it provided it in SI Appendix.

Since transcript degradation is fast compared to dilution (growth rate), we assume that transcripts are in quasi-steady state, and summing on all genes *i* we obtain

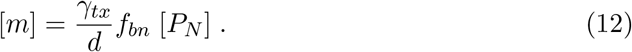

Hence in a given steady-state growth condition the total mRNA concentration is a constant, which will vary with growth rate through RNA polymerase abundance and specific binding. Assuming that *dm/dt* = *λm* leads to the different expression

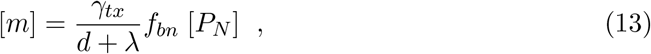

which reduces to Eq. [12] as long as *d* ≫ *λ*.

Eq. [12] encodes an intuitive picture of mRNA production: transcript concentration is proportional to the protein fraction of RNA polymerases and to a “transcription capacity” *γ*_*tx*_ (which is defined by the ratio between transcription elongation rate *k*_*tx*_ and the total length of the genes forming RNA polymerases *γ*_*tx*_ := *k*_*tx*_*/L*_*N*_), while it is inversely proportional to the transcript degradation rate *d*. Note that this expression is equivalent to the equation 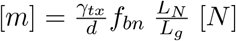 with [*N*] the RNA polymerase concentration, and the term 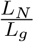 is a conversion factor allowing to pass with ease from abundances of proteins of the polymerase class *N* to the number of polymerases, assuming perfect stochiometry.

Equivalently, Eq. [12] can also be written as

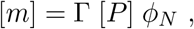

with 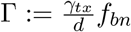 depending on the transcription efficiency *γ*_*tx*_*f*_*bn*_ and transcript degradation *d*, and [*P*] being the overall protein concentration.

Combining these considerations with Eq. [7] leads to the following expression of the growth rate in terms of protein fractions only,

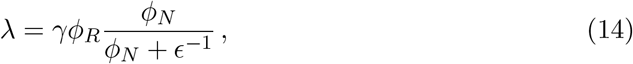

where

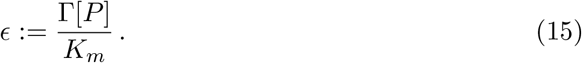

While Eq. [7] couples growth and mRNA concentration, Eq. [14] unfolds the link between growth and the fraction of RNA polymerase-associated proteins *ϕ*_*N*_. We note that *ϵ* is not a phenomenological parameter but emerges as a compound parameter from our model. Specifically, *ϵ* incorporates a purely transcriptional “supply” term Γ, which represents the mRNA concentration per unit of RNA polymerase fraction, and a purely translational “demand” term *K*_*m*_, discussed above, which corresponds to the typical amount of transcripts needed by translation. Equivalently, *ϵ* can also be seen as the increase in mRNA concentration (in units of *K*_*m*_) per units of added RNA polymerase fraction as 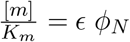. Therefore, this parameter can also be interpreted as the mRNA transcription capacity rescaled by the the amount of needed mRNA. The term *ϵ*^−1^ represents the proteome fraction of RNA polymerase-associated proteins around which the CF-LIM regime becomes relevant. If *ϕ*_*N*_ becomes significantly smaller than this quantity, then growth depends on the available fraction of RNA polymerases. At the opposite situation, if the cell allocates an RNA polymerase fraction *ϵϕ*_*N*_ ≫ 1, then ribosomes become scarce (TL-LIM).

Thus, the ratio *ϵ* can be interpreted as a “supply-demand trade-off” between transcription and translation. This parameter can be experimentally perturbed in different ways, acting on both transcription (and RNA pools) and translation (and ribosome pools). Here, we will focus on changes in transcriptional supply that vary Γ (as it happens for example for transcription-targeting drugs). Instead, in the following section we will exploit its dependence on the transcripts degradation rates to quantify transcription limitation under protein overexpression of unnecessary proteins.

### Competition for transcripts increases the cost of unneeded protein production

This section explores the consequences of transcriptional limitation on the cost of unnecessary protein expression in the competition regime *versus* translation-limited growth (Fig. 2A). Overexpression of unnecessary “burden” proteins that do not contribute to growth helps determining which factors are limiting for growth (59; 60; 11; 7; 32). Experiments can induce this burden by different methods. For example, Scott *et al*. (7) over-expressed *β*-Galactosidase from an IPTG-inducible gene on a medium-copy plasmid in *E. coli*.

**Fig 2.**
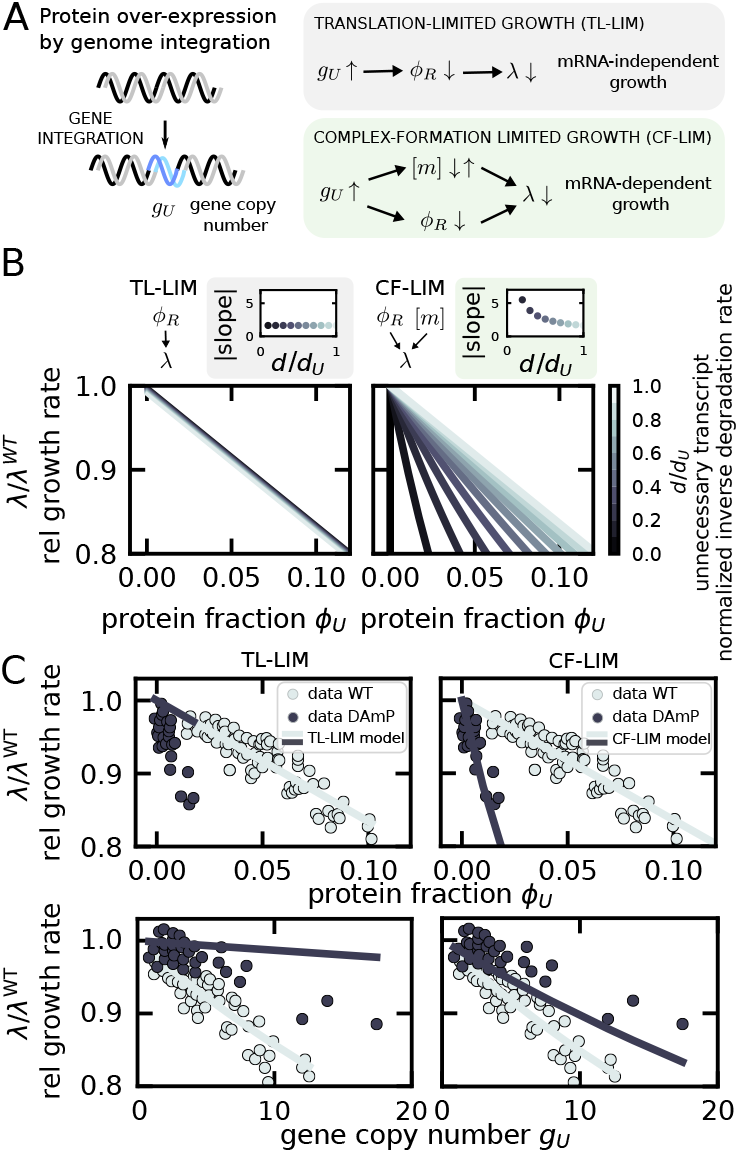
If formation of the ribosome-mRNA complex is limiting, mRNA levels contribute to the protein expression cost. (A) The expression cost of an unneeded protein was probed experimentally in refs. (7; 32) by integrating into the genome highly-expressed unnecessary genes (possibly in multiple copies), labelled by *U*. In the CF-LIM regime, the presence of such genes reduces both mRNA concentration [*m*] and ribosomal protein fraction *ϕ*_*R*_, and the decrease of total mRNA level contributes to a decrease in growth rate. Conversely, in TL-LIM only a reduction of ribosomal levels causes a reduction of the growth rate *λ*. (B) The model predicts a drop in growth rate (quantified by the relative growth rate *λ/λ*^*W T*^) as a function of protein fraction *ϕ*_*U*_ of the unnecessary proteins, and how such trend changes as the degradation rate of the unnecessary transcript *d*_*U*_ varies (right panel, where *d/d*_*U*_ is the inverse degradation rate of the unnecessary transcript normalized to the average degradation rate of the other transcripts). The left panel shows the prediction in TL-LIM. The right panel shows the prediction in the CF-LIM regime. Above the two plots, the sheded boxes show how the slopes of the curve *λ/λ*^*W T*^ (*ϕ*_*U*_) change with *d/d*_*U*_ under translation limitation and complex-formation limitation respectively. (C) *S. cerevisiae* data from ref. (32) falsify a scenario of translation-limited growth and show the trends predicted by the CF-LIM regime. The left panel shows the comparison between the data (circles) and a TL-LIM model (solid lines) plotted both as a function of *ϕ*_*U*_ (top panels) and gene copy number *g*_*U*_ (bottom panels). Light-grey circles represent data corresponding to stable transcripts (*d/d*_*U*_ ≊ 1). Dark-grey circles represent unstable trascripts (*d/d*_*U*_ ≊ 0.08). Dark- and light-grey lines are model curves for the two conditions. The light-grey is obtained by fitting the model, while the dark-grey lines is a prediction of the model. The right panels show the comparison between the data (circles) and our model of the complex-formation limiting regime (CF-LIM, solid lines), using the same color-code. The gene copy-number predictions reported here refer to presence of RNAP homeostasis, see Fig. S4.

Kafri *et al*. (32) generated a library of *S. cerevisiae* with an increasing number of mCherry reporters integrated in their genomes. This dataset quantifies the growth rate of strains expressing this unnecessary protein with different copy numbers (1-20), in wild-type (WT) and in a mutant with increased degradation rate for the mRNA of the reporter gene (DAmP - decreased abundance by mRNA perturbation), and it allows us to test our model.

To model the presence of an unnecessary “type-*U*” gene, inspired by the experimental approach of ref. (32), we considered a model variant with two classes of mRNA, the “typical” pool, and an unnecessary and unstable type-*U* mRNA. We explored this model without imposing optimization of growth rate in response to perturbations. We considered two ways of modulating protein overexpression: (i) by varying the copy number of the gene *g*_*U*_ within the cell and (ii) by changing the degradation rate *d*_*U*_ of the corresponding transcript. The former perturbation globally tunes the growth cost of protein overexpression. Increasing the copies of the genes *g*_*U*_ always increases the growth cost, regardless of which factors are limiting. Instead, tuning the degradation rate modifies the supply-demand balance between transcription and translation, impacting the relative contribution of transcription and translation burden to the overall growth cost. In the limit where unnecessary-mRNA degradation occurs infinitely fast, the presence of unnecessary genes will not affect ribosome allocation or translation burden. More generally, increasing the degradation rate of unnecessary transcripts reduces the translation burden since fewer active ribosomes are removed from the pool of available ones. However, the presence of unnecessary genes still affects overall mRNA content since they remove active RNA polymerases from the pool of all other transcribed genes (46).

In our framework, ribosomes are not necessarily the only growth-limiting component. Indeed, Eq. [7] implies that unnecessary proteins may affect growth both through their impact on ribosomes and through their effect on total mRNA content (as well as through available RNA polymerase allocation, as shown by Eq. [14]). To predict the burden of unnecessary proteins, it is therefore necessary to determine their effect on mRNA as well as on ribosomes. For the reasons explained above, the modulation of the unnecessary transcript degradation rate (32) is an ideal testing ground for our model as it allows to tune the relative contribution of translation and transcription to the growth cost. Increasing the degradation rate of unnecessary transcripts decreases the translation cost (as fewer ribosomes are allocated to unnecessary transcripts).

In order to pursue this question mathematically, let us first consider the two limit cases [*m*]*/K*_*m*_ → ∞ (complete translation limitation) and [*m*]*/K*_*m*_ → 0 (strong complex-formation limitation). By labelling as wild-type (WT) all quantities without overexpression burden, we find,

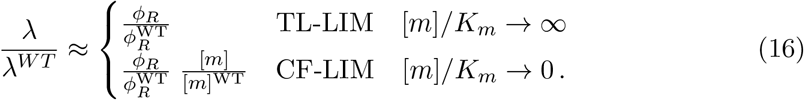

Equation [16] shows that, in these two limiting cases, the relative growth is determined by the ratios 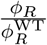 and 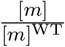. Our framework provides an expression for these ratios under the assumption that unnecessary proteins compete for resources equally with the other protein sectors.

For the ribosomal protein ratio, we recover the well-known result (7)

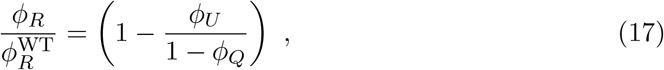

where *ϕ*_*U*_ is the fraction of the proteome taken up by the unnecessary protein, while *ϕ*_*Q*_ is the fractional size of a “*Q* sector” whose size remains constant across growth perturbations (by negative auto-regulation for instance) (7; 11).

Regarding the *Q* sector we need to distinguish the following two possible scenarios, which make a difference for the mRNA ratio: (i) RNA polymerase proteins may not be part of the *Q* sector, therefore their fraction changes across growth conditions, or (ii) RNA polymerase proteins may be part of the sector *Q*, and consequently they remain constant. Although the first hypothesis may appear more conservative, as it does not imply the existence of a circuit that keeps constant the fraction of RNA polymerases, Balakrishnan et al. (30) show experimentally that RNA-polymerase proteins are part of the *Q* sector in *E. coli*, and their effective availability is regulated by the anti-*σ*^70^ factor Rsd. As different organisms may not regulate the availability of RNA-polymerase proteins in the same way, we decided to explore both scenarios with our model.

We begin by considering scenario (i) where RNA polymerase is not a part of the *Q* sector. In this case, our model provides the following expression for the mRNA ratio

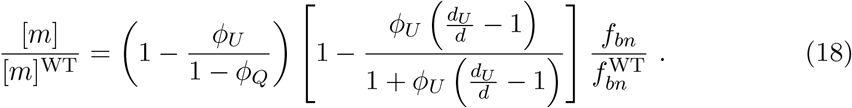

In Eq. [18], the first term 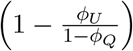 represents the decrease in the RNA polymerase fraction 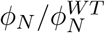 due to the burden from the unnecessary protein. In scenario (ii), RNA polymerase fraction cannot change, and consequenlty this term is simply 1. The second term 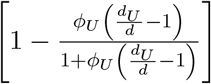 always corresponds to a decrease in mRNA content due to the reduced mRNA stability of the unnecessary gene (hence the term is 1 if *d*_*U*_ = *d*). The third term of Eq. [18], 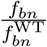represents the relative fraction of bound and active RNA polymerases. As the exact expression of the ratio 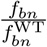 in terms of the model parameters is not particularly transparent, we provide it in SI Appendix. We proved (see SI Appendix) that such ratio can either increase or decrease as extra copies of the unnecessary genes are added. In particular, if *f*_*bn*_ ≈ 1 before the perturbation, mRNA concentration always decreases with unnecessary protein expression 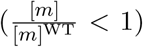. This is a consequence of two basic mechanisms: first, the reallocation of RNA polymerases to transcribe the unnecessary genes (thus competing with RNA-polymerase genes); second, the presence of fast-degrading unnecessary transcripts decreases the lifetime of the average transcript. Note that, if the fraction of RNA polymerases remains constant as in the scenario in which RNA polymerases belong to the *Q* sector, mRNA decreases only due to the second mechanism. Conversely, we find that mRNA concentration can also *increase* if *f*_*bn*_ is sufficiently smaller than 1 before the perturbation and RNA polymerase proteome fraction is maintained homeostatically upon expression of unneeded proteins (see SI Appendix and SI Fig. S10). In this scenario, the free RNA polymerase is recruited by the extra gene(s), increasing the overall transcriptional capacity (hence more RNA polymerase is specifically bound after the perturbation than it is before).

To illustrate the effect of transcript dependence (via transcript-ribosome competition) on the cost of protein overexpression, we now consider the two limits of Eq. [16] in conjunction with Eq. [17] and [18]. Fig. 2B shows the main results of the model in the scenario where the RNA polymerase fraction is constant across growth conditions. Fig. S2 shows the results for the scenario of variable RNA polymerase sector. Both scenarios can reproduce the data, but the latter can match the data only in the regime where the housekeeping Q-sector is absent, which we deem to be less realistic. We also stress that the qualitative results are independent of whether the RNA polymerase sector is constant or variable across growth conditions. The plots in Fig. 2B show the relative growth rate (the ratio 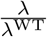) against the unnecessary gene protein fraction *ϕ*_*U*_ in the two limiting regimes. Under translation-limitation (TL-LIM), Eq. [17] predicts that the growth rate decreases linearly with the protein fraction *ϕ*_*U*_. In addition, increasing the degradation rate *d*_*U*_ of unnecessary transcripts has no effect on the slope of the curve, and consequently all the curves collapse (as observed for *E. coli* in ref. (7)). Under competition for transcripts (CF-LIM), the growth rate still decreases linearly with the protein fraction *ϕ*_*U*_. Yet, the response of the system to decreasing the unnecessary transcript lifetime is very different: the slope of the plot of relative growth rate *vs* unneeded protein fraction does not stay flat but increases with decreasing stability, indicating that higher growth cost per protein fraction stems from the decrease in total mRNA content.

The results above imply that the different behavior of overexpression experiments under increasing transcript instability provides a way to probe whether the regime of competition for transcripts described by our model is in place. In brief, the lack of curve-collapse under a change of transcript lifetime could be the result of a CF-LIM regime.

### A model exclusively reliant on translation-limited growth cannot adequately explain unnecessary protein cost data in S.cerevisiae

A study of Kafri and coworkers (32) has performed precisely such an experiment in *S. cerevisiae*. The experiment involved integrating an mCherry reporter into the genome in multiple copies and modulating the transcript’s lifetime using an antibiotic resistance cassette that inhibits termination in the mCherry gene, a technique known as DAmP (decreased abundance by mRNA perturbation). Although the reduction in the DAmP transcript’s lifetime was not directly quantified, the study showed a roughly 10-fold decrease in protein levels of the construct. We analyzed the data from this study and quantified the protein fraction of the unnecessary protein, as explained in detail in the SI Appendix and Fig. S3. We used these measurements to estimate the reduction in transcript lifetime without assuming any specific limitation regime (SI Appendix), and found a corresponding 10-fold decrease. We then incorporated this parameter into our model and observed its predictions under different limitation regimes. Fig. 2C compares the data to the predictions of our model under transcript’s lifetime modulation, which do not rely on any adjustable parameters.

It is important to highlight in which sense our model makes genuine predictions in this context, since it refers to measurements made in the past. We set up the model in a way that only certain available data, specifically the WT experiments, are needed as an input, i.e. to fix the parameters. Once the parameters are fixed, we consider some independent observations, the experiment with the DAmP construt, as predictions that different model variants may or may not capture. Data corresponding to different transcript stability do not collapse (32), suggesting that a model exclusively reliant on translation-limited growth cannot adequately explain these observations. Instead, a model describing the complex-limited regime (CF-LIM) reproduces quantitatively the experimental trends. Finally, we note that the study of Kafri and co-workers (32) also quantified the trend of the relative growth rate against the gene copy number. Our framework also makes correct predictions for this quantity (see bottom of Fig. 2C and Fig. S4). We note that a 10-fold reduction of unnecessary mRNA lifetime can also be obtained directly by fitting the data of growth cost as a function of gene copy number from ref. (32) (Fig. S8), leading to the same independent prediction of Fig. 2C. In their work, Kafri and colleagues also state that perturbing the transcript stability of the unneeded genes induces a 30-fold decrease in the unneeded mRNA in correspondence to the 10-fold decrease of protein levels used in our estimate. Instead, our model would predict the same fold change for unnecessary mRNA and protein levels. Possible explanations include a larger-than-predicted decrease in total mRNA content, reduced translation burden due to reduced misfolding and protein damage (a mechanism not taken into account by our model, but observed in similar experiment in yeast (61)) or the importance of post-transcriptional regulation of gene expression.

In Fig. S5 we also compare TL-LIM and CF-LIM predictions with data from two further nutrient perturbations explored in ref. (32), where Synthetic Complete media was depleted of either nitrogen or phosphate. As expected, in all conditions a TL-LIM model cannot describe the growth rates of the DAmP strain. Conversely, a CF-LIM model can describe all the data, with different values of *K*_*m*_ for different DAmP curves. According to this model fit, the low nitrogen condition appears to be the one with the weakest CF-LIM regime, which is consistent with our microscopic interpretation of *K*_*m*_, as this parameter is proportional to the translation elongation rate, which should decrease in low nitrogen. Surprisingly, despite expectations, the reference condition displays stronger mRNA limitation compared to low phosphate. This discrepancy suggests a potential transition to a regime where the CF-LIM scenario is not accurate and mRNA or rRNA are fully limiting.

To provide a comprehensive analysis, we also evaluated the main predictions under a transcription-limited TX-LIM regime with variable and constant RNA polymerase fraction across growth conditions (SI Fig. S6 and S7). Our findings indicate that (i) the TX-LIM regime consistently predicts a higher cost compared to the complex-formation limited CF-LIM regime under variable RNA polymerases, and (ii) with RNA polymerase homeostasis, the TX-LIM regime predicts no decrease in the growth rate without additional perturbation of transcript stability.

All the above considerations apply under the simplifying hypothesis that most RNA polymerase is specifically bound, *f*_*bn*_ ≃ 1. We also considered the case where *f*_*bn*_ is significantly smaller than 1. As mentioned above, in this situation the fraction of bound RNA polymerase also changes upon the perturbation (see SI Appendix), but our main conclusions do not change. Indeed, Fig. S12 shows that this scenario also replicates the same qualitative change in the slope of the relative growth rate under perturbation with unnecessary protein production with altered mRNA stability.

While the dataset of ref. (32) provides a natural testing ground for the mRNA dependency of the unnecessary protein cost, it is natural wonder whether an equivalent analysis might be performed also in *E. coli*. However, we could not find an equivalent data set for this model organism. Unpublished experimental data in *E. coli* (62) indicate that transcripts with differently efficient ribosome binding sites cause equivalent cost in terms of growth rate, which only depends on the fraction of unneeded proteins *ϕ*_*U*_. In *E. coli*, it is documented that ribosome stalling leads to the nonsymmetric degradation of mRNA, hence to untranslated mRNA (63; 64; 65). Roughly, this makes transcript degradation rate inversely proportional to ribosome mean coverage, which is in turn proportional to initiation rate (ribosome binding site efficiency). For example, Stoebel and coworkers (65), by looking at unstable proteins argue that the cost of unnecessary proteins primarily resides in the production process rather than the product itself. By deleting the ribosome binding site (RBS), they show that the deletion reducing translation also affects transcription, making it difficult to isolate the specific costs of transcription and translation in their experiment. An early study compared the physiological impact of lacZ expression from several ribosome binding sites with different affinities (66). They reported that mRNA half lives correlated with the strength of the ribosome binding sites, but they saw little or no variation in RNA polymerase availability on other genes or total protein synthesis rate. They concluded that the limiting factor for protein production was the availability of free ribosomal subunits. Finally, a recent study performed cloning in *E. coli* of a library of constructs expressing green fluorescent protein (GFP) fused to a 96-nucleotide sequence from a plasmid on the N-terminus (67). The authors design a library of sequences to explore sequence-intrinsic effects on translation efficiency. The promoter used in this study is unmodified and inducible by IPTG, with saturating induction. Some variation in mRNA levels abundances is observed for these transcripts, and it is mainly attributed to modifications of their degradation rates. In their work, the authors show that the growth rate across strains decreases as mRNA levels of the over-expressed protein increase, even at fixed over-expressed protein content. This result suggests that transcription also contributes to the growth-rate cost of GFP in a way that is independent of the expected cost due to the proteome fraction. This is reminiscent of our results in Fig. 2, although we stress that the experiment of Cambray et al. (67) over-expresses unneeded protein in a different manner than the one described by our model (the latter shown in Fig. 2). Our preliminary calculations also indicate that RNA polymerase sequestration from rDNA could also contribute to physiological changes caused by unnecessary protein expression at the transcriptional level (see SI Appendix).

The model formulated thus far outputs the cellular growth rate given ribosome fraction and total mRNA concentration. However, we did not put forward any hypothesis on how the cell chooses such inputs. If such quantities contribute setting growth rates, given that growth rate is a crucial component of evolutionary fitness, they are likely to be strongly regulated. Indeed, it is well-known that ribosomal proteins are tightly regulated in fast-growing organisms such as *E. coli* and *S. cerevisiae* (7; 32). Our framework can describe regulation of gene expression in a simple way. The next section describes how our model can encode different regulatory strategies, and examines the regulation strategy that maximizes the growth rate (10), formulating predictions for changes in mRNA and RNA polymerase abundance across nutrient conditions under such strategy.

### A complex-formation limiting regime explains the decrease of total mRNA in poor nutrient conditions in *E. coli*

Our model outputs the cellular growth rate given the ribosome allocation and total mRNA concentration via Eq. [7], which can be rewritten in terms of RNA polymerase allocation, as shown in Eq. [14]. However, we did not discuss the biological mechanisms responsible for fixing these allocation parameters. This section describes how our model encodes different regulatory strategies, and examines critically the role of an optimal allocation strategy maximising the growth rate (10; 41), which predicts changes in mRNA and RNA polymerase abundance across nutrient conditions. We will compare our predictions with recent data in *E. coli* reporting on an “mRNA growth law” (30), which we will argue can be captured by our model. This data set comprises many measurements, but we will mainly use the data on total mRNA, ribosome allocation, and RNA polymerase protein components availability (and sequestration) across growth conditions. The physiological features of this strain across growth conditions are very precisely characterized quantitatively by a number of studies, which allowed us to fix all the parameters of our model (see below). Incidentally, we note that we focus on the Balakrishnan data set because it offers comprehensive mRNA and proteome data, aligning with our model’s scope. Other studies (41; 9) have used diverse datasets of proteome allocation, and our model successfully replicates these data as a result of its structure inherited from previous frameworks (9; 7; 33; 47).

In order to formulate our model predictions, we focus on the proteome composition Φ = {*ϕ*_1_, …, *ϕ*_*i*_, …, *ϕ*_*S*_}, where *S* is the number of sectors. As nutrient conditions change, cells adjust their growth rate by changing their proteome composition (11), which we can summarise with the remodelling Φ → Φ′. Such changes require the presence of specific molecular pathways able to sense the nutrient conditions and relay the signal to the gene expression machinery (63; 68; 57; 69). While a detailed mechanistic understanding of such architecture is only at its dawn (43), we have already discussed how previous work has shown that often simplifying assumptions such as flux balance and growth rate optimization allow to formulate effective predictions. Such coarse-grained models bypass an explicit description of the sensing-regulation mechanisms, whose function must be at least in part to enforce growth optimality (7; 10).

We will now consider the scenario in which cells regulate resource allocation to achieve an optimal regime for growth, and look for the proteome partitioning assignment Φ^*^ that maximizes the growth rate under different perturbations, analyzing the results critically in a following step. Since at steady state proteome composition reflects proteome allocation, this is equivalent to looking for the RNA polymerase allocation assignment that maximizes the growth rate under different perturbations. In SI Appendix (sec. S7 and also Fig. S9) we discuss in more detail the correspondence, valid in balanced exponential growth, between transcript and protein fraction 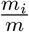 and 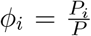 with the fraction *ω*_*i*_ of RNA polymerases transcribing a gene of type *i*.

We proceed by analyzing the predictions of our model under the assumption of growth optimality. We have already discussed how the model links growth rate to ribosome and RNA polymerase allocation via Eq. [14]. In order to optimize the growth rate, we need to find an assignment of the RNA polymerase allocation *ϕ*_*N*_ and ribosome allocation *ϕ*_*R*_ such that *λ* is maximal. Importantly, *ϕ*_*N*_ and *ϕ*_*R*_ also need to satisfy a number of constraints (Fig. 3A). A trivial constraint is normalization: as *ϕ*_*i*_ are protein fractions, their sum Σ_*i*_ *ϕ*_*i*_ must add to unity, and such sum includes *ϕ*_*N*_ and *ϕ*_*R*_. In addition, we assume a “flux balance” condition (7), according to which regulatory constraints must match the influx of protein precursors such as amino acids to the flux of protein synthesis. Specifically, if *ϕ*_*C*_ is a protein sector that synthesizes and imports precursors, the precursors influx (per unit of time) is *dA*_*in*_ = *ϕ*_*C*_ *dP*. The parameter *ν* represents the efficiency of converting nutrients to precursors, often called nutrient quality (7; 8). Biologically, this parameter depends both on the nutrient type and on the metabolic efficiency of the cell. The outflux of precursors due to conversion into proteins is 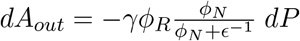. By the assumption of flux matching, we impose *dA*_*in*_ = *dA*_*out*_ and obtain

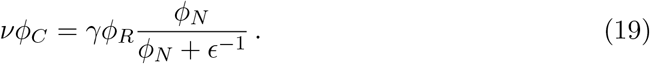

**Fig 3.**
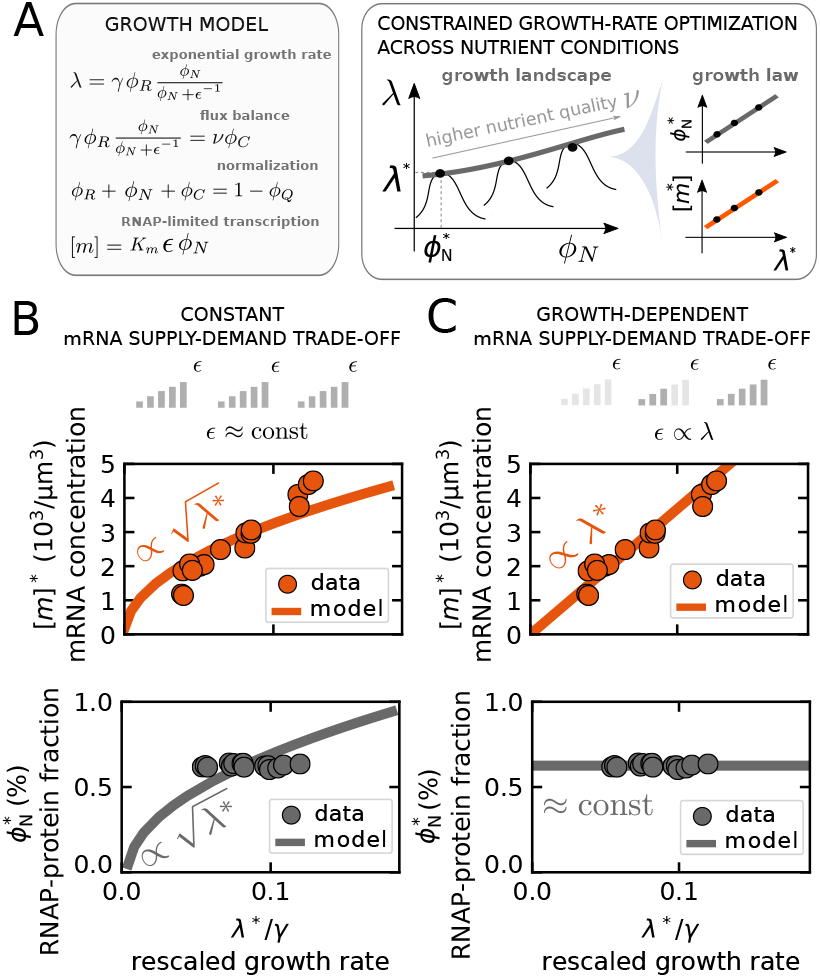
The mRNA levels that maximize growth rate vary across nutrient conditions. (A) The model predicts proteome composition and mRNA levels across nutrient conditions under the assumption that RNA polymerase (RNAP) allocation optimizes growth rate. The left panel illustrates the basic mathematical ingredients of the model, including the dependency of the growth rate on ribosomes and RNAPs, as well as the flux-balance constraint between protein-precursors metabolism and the production of housekeeping proteins. The right panel shows how the model ingredients lead to a nutrient-dependent growth “landscape” where the growth rate is a bell-shaped function of RNAP levels. By taking the maximum of the curve, the model predicts the optimal RNAP and mRNA levels. Such predictions can take the form of a growth law (10) by plotting such levels against the corresponding optimal growth rate. (B) Under growth-rate optimization and constant mRNA supply-demand trade-off *E* across nutrient conditions, the model predicts that both the total mRNA concentration (upper panel, orange solid line) and the fraction of RNAP proteins in the proteome (lower panel, grey solid line) increase as the square root of the growth rate. (C) When growth-rate optimization is combined with the constraint of a trade-off parameter *E* linearly increasing with growth rate (motivated by the data), the model predicts that the total mRNA concentration increases linearly with the growth rate (upper panel, orange solid line) while the fraction of RNAP proteins remains constant (lower panel, grey solid line). Both panel (B) and (C) show *E. coli* mRNA and RNAP data from ref. (30) across nutrient conditions (orange and grey circles), validating the scenario of increasing transcriptional activity (panel C) for this model organism.

Combining our framework with these constraints, we considered a minimal model with only four protein sectors: the RNA polymerase sector (proteome fraction *ϕ*_*N*_), the ribosomal sector (*ϕ*_*R*_), a catabolic sector (*ϕ*_*C*_) and a housekeeping sector (*ϕ*_*Q*_) that does not change with growth conditions (7; 70), and which effectively sets the maximum level *ϕ*^*max*^ = 1 − *ϕ*_*Q*_ that any other sector can reach. Therefore, the normalization condition reads

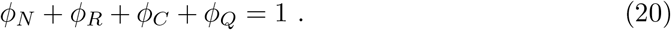

By combining Eq. [14], [19], and [20], we solved analytically the constrained optimization problem as illustrated in Fig. 3A.

A key quantity to describe the role of transcription in growth is the proteome fraction of RNA polymerases. Such quantity takes the following expression in the solution of the following optimization problem,

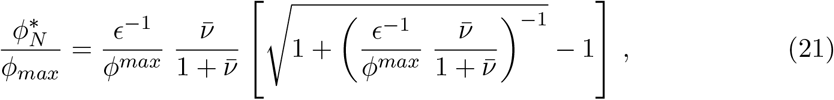

where we indicate with the superscript ^*^ the optimized variables, and 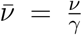 is a dimensionless nutrient quality. Equation [21] shows that 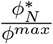 solely depends on a single composite parameter 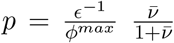, which varies changing either the dimensionless nutrient quality 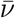 or the mRNA trade-off parameter *ϵ*. The Materials and Methods and SI Appendix provide details of these calculations.

We note that if the supply-demand trade-off parameter *ϵ* is a function of the nutrient quality *ν*, the expression above still holds. Motivated by the findings of ref. (30), we considered two particular cases, which we can be compared with data. In the first scenario, *ϵ* is a constant that does not depend on the nutrient quality. Conversely, in the second scenario *ϵ* takes the form 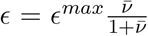, which is equivalent to stating that *ϵ* is linearly proportional to the growth rate (see SI Appendix and Fig. S13).

Fig. 3B shows the model predictions for overall mRNA concentration in the first scenario. As nutrient conditions improve (increasing 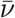), the RNA-polymerase protein fraction *ϕ*_*N*_ increases under the optimal solution. As total mRNA levels are proportional to *ϕ*_*N*_, they also increase with increasing nutrient quality. Interestingly, the model predicts that such quantities have a square-root dependency on the growth rate as opposed to the typical linear laws (see also SI Appendix for an extended discussion). Fig. 3B also shows *E. coli* data from ref. (30). These data agree qualitatively, but not quantitatively, with the mRNA trend in the data and fail to capture the measured constant RNA-polymerase protein fraction. Finally, Fig. 3C shows the result of the optimization under the assumption that the transcription-translation trade-off is proportional to growth rate *ϵ* ∝ *λ*^*^. In this case, the model predicts that mRNA increases linearly with the growth rate, while the the fraction of RNA-polymerase proteins stays constant. As it happens in *E. coli*, the model also crucially predicts that there is no need to tune the proteome fraction of RNA polymerase in this scenario, as shown by Fig. 3C. Specifically, Balakrishnan and coworkers show that *ϕ*_*N*_ stays constant across growth conditions (gray circles), and that the expression of *σ*-70 sequestration factor Rsd decreases with growth rate, modulating mRNA synthesis fluxes at constant fraction of RNA polymerase (30). In our framework, the RNA polymerase sequestration at slow growth is achieved by the linear reduction of the transcription-translation trade-off parameter *ϵ*, which can be interpreted as a lowering of the fraction of actively transcribing RNA polymerases *f*_*bn*_ Fig. S13) in order to maintain the linear density of ribosomes on transcripts constant across conditions (30), essentially by tuning the ratio between transcription and translation elongation rates. This can be interpreted as a signature that growth rate is not the sole quantity being optimized in this context, but maintaining an overall ribosome flux, related to ribosome linear density along transcripts (54; 52) may also be a target of the system (realized in our model as a constraint on *ϵ*, and mechanistically as increasing RNA polymerase sequestration at decreasing growth rates). Indeed, our analysis also shows that such density increases if *ϵ* stays fixed, but remains constant under growth-dependent *ϵ* (Fig. S14 and S15).

An important (more general) aspect is that whatever the architecture causing this trend, and whatever the scenario, in both data and model mRNA concentration decreases with decreasing growth rate, suggesting that at slow growth competition for transcripts is likely relevant as the mRNA concentration becomes closer to *K*_*m*_, the concentration scale that determines to what extent mRNA affect the growth rate.

We note again that this slow-growth regime is also the one where the simplified model considered here for illustration becomes approximate. Specifically, it neglects documented processes in *E. coli* leading to a reduction of the ribosomes that are effectively available for translation, due to active sequestration, and protein degradation (15; 58; 47); only free ribosomes due to mRNA competition are included in our model.

### Under transcription-targeting drugs, RNA polymerase levels must increase to achieve optimal growth, but ribosome levels may not

The expressions obtained in the previous section can be used to investigate the effect of transcription-targeting drugs on growth (7; 35; 9). Previous studies have considered growth-rate changes under these perturbations (35; 9). We will focus here on ribosome allocation, and compare with *E. coli* data on *ϕ*_*R*_ changes under transcriptional inhibition (rifampicin) from ref. (7), once again from the strain where all the parameters have been precisely characterized and they can be easily fixed in our model.

Specifically, we modeled a transcription-targeting drug that affects the global mRNA concentration [*m*], and we analyzed the optimal and non-optimal re-arrangement of coarse-grained proteome composition in response to such challenges, starting from a growth-rate optimal condition. Our model shows in a straightforward way (Eq. [7]) that in the regime where growth is dependent on mRNA concentration, transcription-targeting drugs also directly affect growth. We can ask the further question of how the proteome responds to such perturbations.

Such drugs might target different steps of the transcription process to reduce overall mRNA concentration. From the perspective of our model, we focus on transcription-targeting drugs that modify the trade-off parameter *ϵ* (Eq. [15]).

We recall that [*m*]*/K*_*m*_ = *ϵϕ*_*N*_, which implies that in principle transcription-inhibitory drugs could also decrease mRNA levels by reducing RNA polymerase expression. Note that *ϵ* is a composite parameter shaped by several distinct biological processes. According to our model, drugs attacking distinct aspects of transcription can cause quantitatively similar results. For instance, drugs targeting transcriptional elongation decrease *ϵ* by lowering the elongation rate *k*_*tx*_; drugs targeting transcriptional initiation decrease *ϵ* by reducing the fraction of active RNA polymerases *f*_*bn*_; drugs targeting transcript stability decrease *ϵ* by curbing the transcript degradation rate *d*. Hence, by focusing generically on how *ϵ* changes, our model makes generic predictions about multiple situations.

To derive the effect of transcription-inhibitory drugs in the model, we follow the same formal steps of the previous section. The growth rate is a function of the coarse-grained proteome composition made of ribosomal, RNA polymerase and catabolic proteins. In addition, it also depends on the mRNA supply-demand trade-off parameter *ϵ* and the nutrient quality *ν*. Mathematically, the growth rate *λ* is a function *f* (*ϕ*_*R*_, *ϕ*_*N*_, *ϕ*_*C*_, *ϵ, ν*) given by equations [14] and [19]. We find the composition {*ϕ*_*R*_, *ϕ*_*N*_, *ϕ*_*C*_} that maximizes the growth rate at given supply-demand trade-off *ϵ*. Consequently, the composition that maximises growth rate is a function of the trade-off parameter *ϵ*. If transcription-targeting drugs effectively vary *ϵ*, the growth-optimal proteome composition as a function of *ϵ* corresponds to the cellular gene-regulatory control that maximises growth rate in response to transcription-targeting drugs - see also the sketch in Fig. 4A.

**Fig 4.**
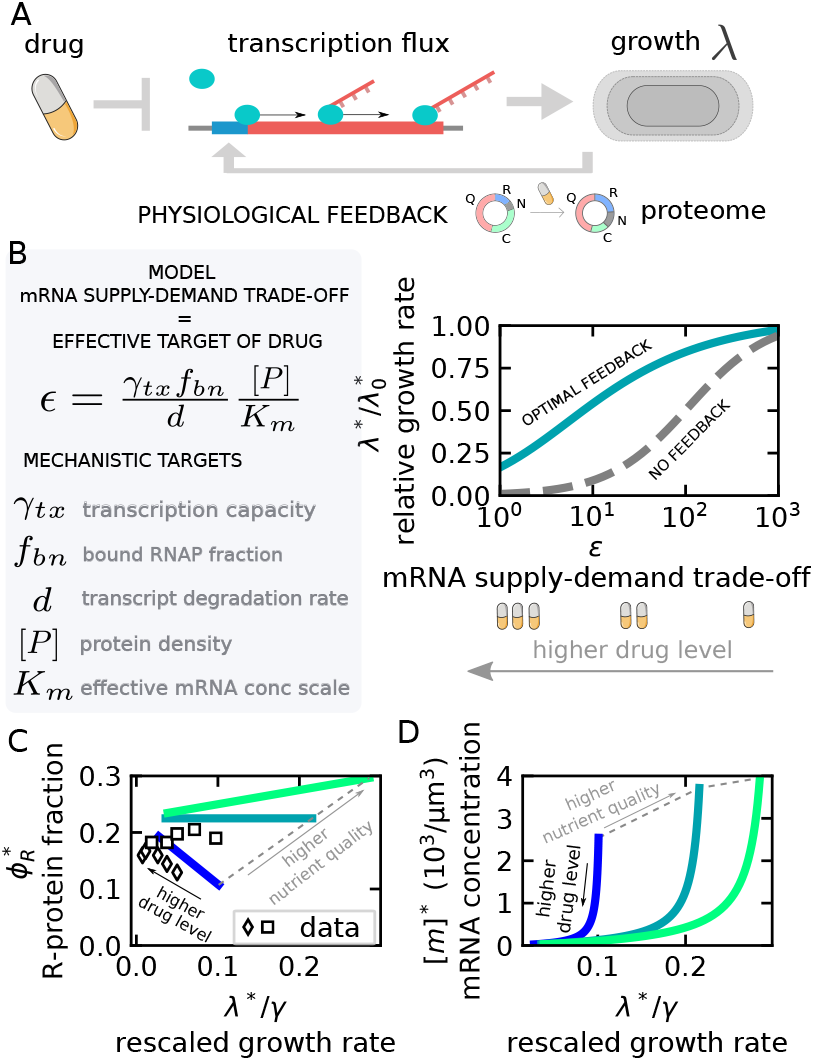
The predicted response to transcription-targeting drugs entails up-regulation of the RNA polymerase sector, but not necessarily of ribosomes. All the predictions refer to the CF-LIM regime. (A) Transcription-targeting drugs affect growth directly, but the overall effect also depends on the physiological response of the cell to reduced transcriptional capacity. Our model predicts ribosome- and RNA polymerase-(RNAP) allocation rearrangements under transcription inhibitors. (B) The grey box illustrates the model expression for the transcription-translation trade-off *ϵ*, i.e., the ability of RNAP to produce mRNA (see Fig. 3A). This effective parameter combines several quantities, including the transcription elongation rate of RNAP on genes, the fraction of gene-bound RNAPs and the mean lifetime of transcripts. The model predicts that drugs attacking any of these parameters modify the way mRNA level affect growth rate only through changes in *ϵ*. The plot shows the growth rate reduction *vs ϵ* under growth-optimized (continuous blue line) and non-optimized (grey dashed line) conditions. Note that 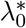 is the growth rate in the unpertubed condition. Proteome re-arrangement optimizing growth rate “rescues” the growth rate decrease under transcription inhibition (reduction of *ϵ*). (C) The predicted ribosomal proteome fraction *ϕ*_*R*_ under growth optimization changes as transcription is inhibited (*ϵ* reduced). Solid lines of different colors indicate different nutrient conditions with darker colors representing poorer media. Importantly, the qualitative trend depends on the nutrient quality. Symbols are experimental data points from ref. (7) (D) The model prediction for mRNA concentration [*m*] changes as *ϵ* decreases under optimal allocation. Different solid lines indicate different nutrient conditions, with darker colors representing poorer media.

Biologically, there can be situations where cells will or will not implement a growth-optimal response strategy, hence it makes sense to compare a growth-optimized scenario with a non-optimized one. Fig. 4B shows how the growth rate slows down under transcription inhibitors when mRNA trade-off *ϵ* decreases with and and without the enforcement of growth-rate optimization. The dashed grey line shows the relative growth rate against the trade-off parameter *ϵ* without any response in proteome rearrangement (Eq. [14] plotted as a function of *ϵ*). The solid blue line shows the relative growth rate under optimization, following the procedure described above. The plot shows that optimal feedback always determines a faster growth rate. We discuss here mainly the assumption that the RNA polymerase fraction can vary under treatment of transcription-targeting drugs. SI Appendix, considers more in detail the response maximizing growth rate under constant level of RNA polymerase fraction.

Besides growth rate changes, our model also produces testable predictions on how the proteome re-arranges under transcription-targeting drugs. The RNA polymerase fraction of the proteome always increases, to compensate the lower transcriptional efficiency with more RNA polymerases (see equation [21]). Despite such increase, overall mRNA concentration still decreases (Fig. 4D), although it decreases less than it would without growth-optimized response. Clearly, RNA polymerase upregulation must come at the expense of other protein sectors. How do the other coarse-grained protein sectors behave? Intriguingly, the ribosomal proteome fraction changes qualitatively differently under treatment of transcription-targeting drugs in nutrient-poor *vs* nutrient-rich media. Fig. 4C shows that the model predicts that, under optimized growth rate, when transcription is inhibited with respect to untreated conditions the ribosomal protein fraction *ϕ*_*R*_ should increase in poorer media, and decrease in richer media. For a “critical” value of the nutrient quality, the ribosomal change does not change at all upon inhibition of transcription. These predictions are in agreement with *E. coli* experimental data under treatment with rifampicin, an inhibitor of bacterial RNA polymerase, hence of global transcription (Fig. 4C, data from ref. (7)).

In other words, data and model appear to agree on the very interesting conclusion that the reaction to transcription inhibition in terms of ribosome allocation changes qualitatively across growth conditions. In the model, the reasons for such a trend change are conceptually subtle, although easy to prove computationally and analytically. In brief, as transcription capacity decreases, the model predicts that the cell will reduce the size of the catabolic sector *ϕ*_*C*_ (see SI Fig. S16) because fewer precursors are being used due lower overall biosynthetic capacity. Hence, we can think that the fraction of the proteome Δ*ϕ*_*C*_ freed by this contraction becomes available to the other sectors. Quantitatively, the proteome fraction change Δ*ϕ*_*C*_ depends on the nutrient quality of the medium. In rich media 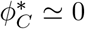, while in poor media 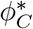 can become considerable. This is because in rich media nutrients are easily imported and catabolized, so 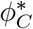 can be small. Part of the freed Δ*ϕ*_*C*_ will increase the fraction of the proteome occupied by ribosomes, offsetting its decrease due to the upregulation of RNA polymerase. If 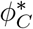 is sufficiently large (which may happen in poor media), the ribosomal fraction *ϕ*_*R*_ can also *increase* as transcriptional capacity decreases.

## Discussion

We presented an organism-agnostic framework describing biosynthesis, accounting explicitly for the two key steps of the central dogma, mRNA and protein production, which can describe the growth laws determined by RNA polymerase and ribosome allocation. The fact that our model is able to formulate correct predictions for both *E. coli* and *S. cereviesiae* supports the hypothesis that unifying principles, due to simple trade-offs (allocation, flux balance, etc.) may apply across organisms and kingdoms, and therefore adds up to the thread of evidence supporting the existence of universal aspects of growth physiology (24; 22; 25; 26; 47). Clearly, further work could investigate the role of specific aspects of different transcription-translation architectures. For example, in *E. coli* transcription and translation co-occur in the nucleoid, while in budding yeast the formation of ribosome-mRNA complexes needs nucleocytoplasmic transport of mRNA, which relies on multiple proteins and organelles. Such differences may affect the parameters leading to regime of competition for complexes, as well as the biological perturbations that affect this regime (34; 71). While the effect of proteome allocation on translation is clear from previous work, general feedbacks between translation and transcription capacity remain relatively unexplored. Our results show clearly that there are interesting physiological and perturbed situations where competition for transcripts sets growth rate. Interestingly, the RNA polymerase proteome sector, while being very small, plays the crucial role of controlling mRNA levels, therefore it can be determinant to decide to which extent mRNAs set growth rate. A clear limitation of our model, particularly in relation to eukaryotic systems, is the assumption of regulated rRNA (and tRNA) levels, which were left implicit. SI Appendix explores model variants in this direction: our preliminary conclusions are that in *E. coli* RNA polymerase sequestration from rDNA should determine minor changes in the total mRNA transcript pool compared to those observed in ref.(30) across growth conditions. In eukaryotes, a further extension of our framework could investigate the role of different dedicated RNA polymerase pools. Previous work has assumed that a perfect coordination of RNA polymerase I and III optimizes growth (22), but no work exists in this context addressing the role of RNA polymerase II. Here, we have decided, in the spirit of minimizing the parameters and because of lack of explicit data, to assume a single RNA polymerase sector also for yeast, but clearly this hypothesis may be turn out to be simplistic in some situations, such as in the presence of RNA polymerase type-specific challenges to transcription.

Recent results in budding yeast support the idea that RNA polymerase II is the main limiting factor for mRNA levels (72). The authors find that as cells grow, both RNA polymerase II recruitment and changes in mRNA degradation rates contribute to maintain mRNA concentration. Additionally, this study appears to support a co-limited regime for transcription, where both gene copy number and RNA polymerases are relevant for total mRNA production. The increase in transcription capacity with cell size has been attributed to a transcription-limited regime, but this may be at odds with the exponential growth of single cells (33). While the current version of our model is not adapted to deal with cell-size, another interesting extension of our work could test a possible role of a complex-formation limiting regime of growth in this process.

As we have mentioned in the Introduction, two recent studies (33; 9) have considered the impact of the transcriptional layer of protein production on growth, but our work differs in several important ways. We built our transcription-translation framework following Lin and Amir (33), who focus on the transcription-limited growth regime (TX-LIM) where *only* mRNA determines growth (and RNA polymerase autocatalysis is essential for exponential growth). Conversely, our model focuses on the regime where both mRNA and ribosomes are relevant for setting growth rate, as a consequence of the competition for ribosomes by mRNA. We note that cells grow exponentially in this competition regime, through ribosome autocatalysis. The study by Roy and coworkers (9) also did not consider this regime. In addition, our study also compares growth-rate maximizing and non-optimal RNA-polymerase/ribosomal allocation across nutrient conditions, while both previous studies focused only on non-optimized relationships. Finally, our framework is very similar to the data-driven model proposed by Balakrishan and coworkers (30), with the advantage of being able to provide a mechanistic interpretation for the factor relating growth rate and total mRNA concentration, and to compare growth-optimized situations with non-optimized ones as done in ref. (10).

In our model, which is focused on the CF-LIM regime, the relationship between mRNA and growth rate is described by Eq. [7] which accounts for the competition for transcripts to the pool of free ribosomes. It is important to note that in the scenario we propose total mRNA concentration affects the growth rate only through the availability of free ribosomes. This is fully consistent with the phenomenological model proposed by Balakrishnan and coworkers (30). However, our contribution is to propose the complex formation limiting (CF-LIM) regime as a mechanistic explanation for the availability of free ribosomes, leading to an explicit expression for Eq. [7]. It is worth noting that the term “mRNA cost” can have a broader meaning in some contexts, which is not explicitly considered in our model (61; 73). In our model, changes in global transcription are mainly associated with the availability of free ribosomes. The process of protein biosynthesis remains ribosome-centric, based on ribosome autocatalysis.

Since the size of the RNA polymerase sector is very small, and there is no associated CF-LIM regime is unlikely to be a consequence of a substantial metabolic burden associated with RNA polymerases or mRNA production in terms of the proteome. Instead, it appears to be a deliberate physiological state, potentially influenced by factors like ribosome traffic, as suggested by Balakrishnan and colleagues, or the objective of orchestrating growth control through various aspects of the central dogma concurrently.

Growth limitations can be important across different perturbed and physiological contexts, therefore we expect that our approach can form the basis for further investigations. A first perturbation where transcriptional limitation is obviously important is treatment of cells with transcription inhibitors. The net effect of protein synthesis inhibitors on growth is mediated by the physiological feedback of the cell in response to drug treatment (7). A well-known example of this is the growth reponse of E. coli to ribosome-targeting antibiotics. Such drugs also induce up-regulation of ribosomal content which partially rescues growth rate decrease. More in detail, the response depends on the growth condition as well as the affinity of the drug to its target (14). The structure of this feedback has been used to predict the shape of dose-response curves for different translation-targeting antibiotics (74). This kind of analysis remains largely open for the case of transcription inhibitors. The relationship of transcription inhibitors with growth rate (in non-optimized conditions) was considered in ref. (9). Our model adds the further step of being able to compare growth-optimized with non-optimized resonse scenarios, and to make definite predictions for ribosomal and RNA polymerase sector response to transcription inhibitors. However, additional elements such as drug affinity and feedback mechanisms (74; 14) may be important to fully understand the physiological response to transcription inhibitors. Future studies could extend our framework in these directions.

A second important perturbation where transcription becomes important for growth is the expression of unnecessary proteins (7; 60; 59; 25). Unnecessary protein expression imposes a cost on growth by affecting the abundance of growth-limiting components. In the standard ribo-centric growth model, the growth rate is proportional to ribosome content (25). Because of finite resources, expression of unnecessary protein decreases the expression of ribosomal proteins, which reduces the number of ribosomes and slows down growth. In our framework, ribosomes are not necessarily the only growth-limiting components. We found that the growth cost depends on both mRNA and ribosomalcontent and provided a quantitative predictive model of the corresponding growth cost. The scenario of predominantly bound and actively translating RNA polymerase predicts a decrease in mRNA concentration with unnecessary protein expression. However, a more recent study by Kafri et al. (73) reported that mRNA concentrations can remain constant or even slightly increase under the burden condition. Our model shows that this could be the case if there is sufficient free RNA polymerase in the unperturbed case (as some studies suggest (72; 75)) and RNA polymerase is maintained homeostatically upon the perturbation (SI Appendix and SI Fig. S10). The increase of mRNA concentration under expression of unnecessary proteins could also be due to physiological feedback mechanisms, such as cell volume regulation (76) or chaperones (61) that are unaccounted by our model. We acknowledge the need for further experimental investigation to address these points.

Assuming our model, the transcription rate of the unnecessary gene always increases the growth cost of overexpression, while the decrease of unnecessary transcript stability decreases the growth cost, because fewer unnecessary transcripts take up ribosomes from the unnecessary mRNAs. Our quantitative framework describes both perturbations jointly. We speculate that this quantitative understanding could be useful to predict the fitness landscape of situations with perturbed gene dosage, such as large-scale gene duplications in absence of dosage-compensation mechanisms (77). In addition, our model provides a method to rigorously test the [m]-dependent growth via transcript-ribosome competition by considering how the cost of overexpression varies as the mRNA stability of the unnecessary protein changes. Hence, the problem of the growth cost of unneeded proteins may be important as an experimental testing tool for the presence of transcriptional limitations in a specific reference regime. Other situations where transcription may become limiting indirectly, as a consequence of orthogonal perturbations are inhibition of translation (as already shown in *E. coli* (64)) and DNA dilution (as shown in yeast under G1/S arrest (31)).

In physiological conditions, our model predicts that under the assumption of transcript competition, optimization of growth rate leads to the observed decrease of total mRNA levels in poor nutrient conditions (in absence of any external perturbations). This prediction is in agreement with the recent experimental observation that total mRNA concentration decreases with decreasing growth rate in *E. coli* (30). However, if no parameters are fixed and growth rate is optimized, the resulting square root growth law in the CF-LIM regime does not match the experimental data quantitatively. Conversely, to achieve a quantitative agreement with the data from Balakrishnan and coworkers, our model needs to add the constraint of a linearly increasing mRNA supply-demand trade-off *ϵ* with growth rate, highlighting the fact that growth-rate optimization is not the sole principle guiding the growht laws. As the authors of ref. (30) have noted, this constraint tunes the effective initiation rate in order to maintain a fairly constant linear density of ribosomes along transcripts across a wide range of growth conditions. We interpret this trend as a signature that ribosome flux (which is related to ribosome density (52)) may be an additional important parameter that cells attempt to regulate. Mechanistically, as the authors show, the increasing trend for *ϵ* is realized in *E. coli* by increasing RNA polymerase sequestration from anti-sigma factor Rsd, indicating that growth rate is modulated by a change of the fraction of active RNA polymerases, instead of the total RNA polymerase levels. We further speculate that this sequestration mechanism could help the system to react faster in nutrient upshifts, as desequestration of RNA polymerase components could be much faster than transcriptional reprogramming (30). We also note that these findings by Balakrishnan and coworkers challenge the dogma that mRNA-specific initiation rates determine protein concentrations: translational characteristics are similar across most mRNAs but RNA polymerase activity is fine-tuned to match translational output, therefore (in *E. coli*) specific protein concentration appears to be primarily set at the promoter level. Transcriptional limitation in physiological conditions was also reported to be relevant for phosphate limitations, in both yeast and *ϵ*.*coli* (29; 73; 25), and may be relevant for regulating cell-cycle dependent expression in budding yeast (78). On more general grounds, an understanding of the link between mRNA concentration and ribosome / RNA polymerase allocation needs a theoretical framework able to link growth rate to mRNA levels (33; 9; 30).

Finally, we note that in transcription-limited situation, as well as under competition for transcripts, mRNA scarcity may contribute to inactive ribosomes (15; 58; 47). Since mRNA concentration can be quite low for slow-growing *E. coli* (30), we speculate that this factor may play a role in setting the fraction of idle ribosomes.

## Methods and Materials

This Materials and Methods section discusses the model ingredients, assumptions, and derives the main mathematical results presented in the text, in particular the expression linking the growth rate to the ribosome fraction and mRNA concentrations, Eq. [7], and the formulas for the cost of protein overexpression Eqs. [16], [17], [18]. See also the SI Appendix for more detailed information on the model.

### Initiation limited fluxes

We consider a situation in which the transcription and translation fluxes 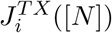 and 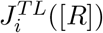 in Eqs. [1] and [2] are initiation-limited:

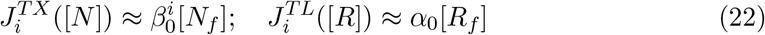

where 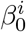 is the rate constant of transcription initiation of the genes belonging to class *i*, and *α*_0_ is the rate constant for translation initiation, assumed to be identical for all transcripts. [*N*_*f*_] and [*R*_*f*_] are the concentrations of free RNA polymerase (RNAP) and of ribosomes, respectively (see SI Appendix). Eqs. [22] relate the biosynthetic fluxes with the process of recruitment and complex formation (RNAP-gene and ribosome-transcript). We approximate for simplicity the current with the initiation rate, neglecting RNAP and ribosome traffic

Each term in this equation can be interpreted in a simple way: (i) 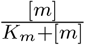 is the fraction of bound ribosomes, (ii) 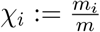 is the fraction of bound ribosome translating a transcript of type *i* and (iii) 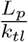 is the typical time to translate a protein. Associating a symbol to each of these quantities, we write schematically

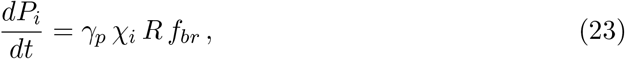

where *γ*_*p*_ is the inverse time to translate the typical protein,, *χ*_*i*_ is the fraction of bound ribosomes translating transcripts of type *i* and *f*_*br*_ is the overall fraction of ribosomes bound to transcripts. This way to represent Eq. [2] is particularly interpretable, and our model has a parallel equation for transcript production (see below).

Assuming perfect stochiometry (i.e. all ribosomal subunits are involved in a functional ribosome), we write the number of total ribosomes *R* in terms of the number of ribosomal proteins *P*_*R*_ as 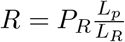, where *L*_*R*_ stands for the number of amino-acids present in a ribosome. Summing Eq. [23] for all classes and dividing by the total protein abundance *P* gives Eq. [7] after defining *γ* := *k*_*tl*_*/L*_*R*_.

### Connecting mRNA concentration with RNA polymerase abundance

Mirroring Eq. [23] for the transcript dynamics (see SI Appendix for details), we write

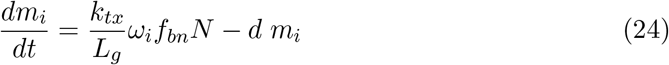

with 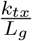, *ω*_*i*_ and *f*_*bn*_ being respectively the inverse typical time needed to transcribe a gene (the ratio of a typical transcription elongation rate and a typical gene length), the fraction of bound RNA polymerases transcribing genes of type *i* and *f*_*bn*_ the overall fraction of RNA polymerases bound to genes. SI Appendix derives the exact expressions for *ω*_*i*_ and *f*_*bn*_. Crucially, they depend only on the binding constants 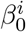 (promoter strength) and the gene copy number *g*_*i*_. Compared to the case of translation, these expressions are more complex due to gene- or sector-specific promoter strengths, although qualitatively they behave very similarly to the corresponding translation quantities *χ*_*i*_ and *f*_*br*_ provided we substitute the transcripts *m*_*i*_ for the genes *g*_*i*_.

Since transcript degradation is fast compared to dilution (growth rate), we assume that transcripts are steady state, and we sum all the amount *m*_*i*_ for all classes *i*, from Eq. [11] we obtain Eq. [12]. We convert the amount of RNA polymerases in number of proteins composing them: 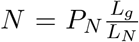, where *L*_*N*_ is the total number of amino acids needed to form a RNA polymerase complex and *L*_*g*_ the typical gene length (perfect stochiometry assumption). This allows us to write the concentration [*N*] in terms of fraction of polymerases *ϕ*_*N*_ and total protein concentration [*P*] as 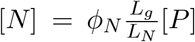. Finally, to derive Eq. [14] we plug the obtained relation 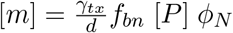 into Eq. [7].

### Regime of limiting complex formation (CF-LIM)

Eq. [7] gives the mRNA dependence of the growth rate via a factor representing the fraction of free ribosomes, which becomes relevant when 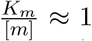. This quantity has a simple interpretation as the ratio between the ribosome-mRNA association and dissociation times

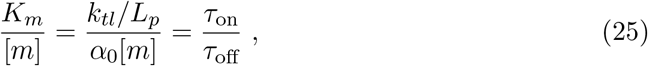

since (*k*_*tl*_*/L*_*p*_) can be seen as the inverse of a ribosome characteristic “dissociation” time, i.e., the time necessary to elongate the typical protein, while a free ribosome takes a time roughly inversely proportional to *α*_0_[*m*] to find a transcript and form a complex, i.e., “associate” with it. The growth rate depends on total mRNA concentration when the time scale of ribosome-mRNA complex formation (*τ*_on_) is comparable to the timescale of full protein elongation (*τ*_off_). The slower complex formation is with respect to full protein elongation, the more dependent the growth rate becomes on mRNA. Eq. [7] provides a general relation between growth rate, ribosome fraction and transcript concentration, without assuming that ribosomes or mRNAs are limiting. Note that this trade-off depends on a “supply” of mRNA and a “demand” from translation: if *τ*_on_ *< τ*_off_ ribosomes accumulate on mRNAs, because supply is scarce, and viceversa if *τ*_on_ *> τ*_off_ the demand for transcripts is scarce with respect to the supply.

In the competition regime described above, it is simple to show that the growth rate 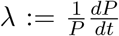 is proportional to a (Michelis-Menten-like) factor 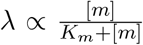 (see SI Appendix). The effective concentration *K*_*m*_ is

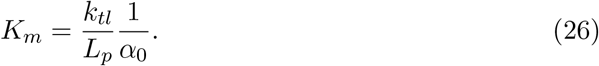

Let us recall that we defined the translation initiation rate as *α* = *α*_0_[*R*_*f*_], where [*R*_*f*_] is the concentration of free ribosomes. Therefore, *α*_0_ is a binding constant that can be estimated with the ratio 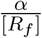. To estimate *K*_*m*_ from literature data we expressed it as

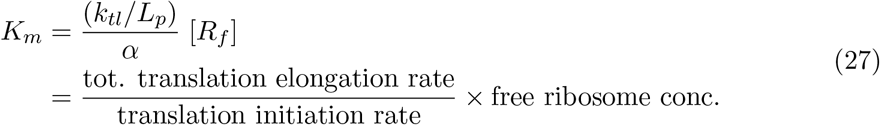

#### Estimate of *K*_*m*_ for *E. coli*

To estimate *K*_*m*_ for *E. coli* we started from the quantities linked to translation elongation. As *k*_*tl*_ ≈ 10 − 20 *aa/s*

For translation initiation, we took *α* ≈ 0.2 − 0.3 *s*^−1^ =≈ 12 − 18 min^−1^ = 700 − 1000 *h*^−1^

#### S. cerevisiae estimate of *K*_*m*_

In yeast, we used *k*_*tl*_ ≈ 10 *aa/s*

### Minimal derivation of the growth cost of protein overexpression

To derive the growth of cost of protein overexpression, we used an expression for the ribosomal-protein fraction fold-change 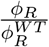 and the total mRNA fold change 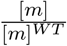 in the presence of the unneeded protein type *U*. Subsequently, we used Eq. [7] to obtain the growth cost (see SI Appendix).

#### Fold change of the ribosomal protein fraction

The fold change of the ribosomal-protein fraction 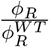 was obtained directly from the normalization condition Σ_*i*_ *ϕ* = 1 by assuming that each protein fraction changes in the same way. With this assumption 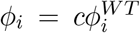for all *i* ≠ *U, Q* (including *R*-proteins). Since 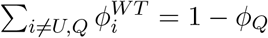, we can find from the normalization condition that *c*(1 − *ϕ*_*Q*_) + *ϕ*_*U*_ + *ϕ*_*Q*_ = 1, which means 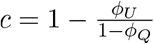. Therefore 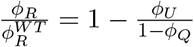, as stated in the main text in Eq. [17].

#### Fold change of the total mRNA content

To derive the change of the total mRNA, we used Eq. [1], obtaining the following expression for the mRNA transcript for sector *i* and *U* (see SI Appendix)

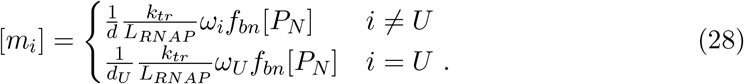

Note that *ω*_*i*_ above is the fraction of RNAPs transcribing transcript type *i*. the total mRNA was obtained by summing [*m*_*i*_] on *i*,

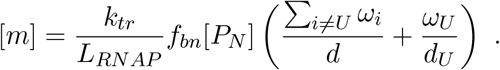

To further simplify this expression, we expressed the concentration in terms of the protein fraction, i.e. [*P*_*N*_] = [*P*]*ϕ*_*N*_ with [*P*] the protein concentration, and used the the fact that Σ_*i*≠*U*_ *ω*_*i*_ = 1 − *ω*_*U*_. Finally, we also used the link between *ω*_*U*_ and *ϕ*_*U*_, i.e., 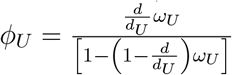 (see SI Appendix and Fig. S9 for the full derivation). Following this procedure, we obtained Eq. [18] in the main text.

## Acknowledgments

We are grateful to Ariel Amir, Terence Hwa, Eyal Metzl Raz, Matteo Mori, and Rami Pugatch for useful feedback on our work. This work was supported by Associazione Italiana per la Ricerca sul Cancro AIRC IG grant no. 23258 (MCL and LC), and by the French National Research Agency (REF: ANR-21-CE45-0009) (LCi). LC was supported by an AIRC Fellowship.

