## Supplementary Material for "How total mRNA influences cell growth"

#### This PDF file includes:

Supporting text
Figs. S1 to S16
Table S1
SI References

### Supporting Information Text

This SI Appendix contains further analytical arguments and derivations regarding our mathematical models, as well as further details on the data analysis procedures and comparisons with data.

### S1. Model of gene expression and cellular growth

The gene expression model introduced in the main text describes the evolution of the abundances of mRNAs and proteins ($m_i$ and $P_i$, respectively) belonging to the class $i$. For the sake of clarity we rewrite here Eqs. [1] and [2] of the main text, which define the model

$$\frac{dm_i}{dt} = g_i \; J_i^{TX}([N]) - d_i^m \; m_i \qquad [S1]$$

$$\frac{dP_i}{dt} = m_i \; J_i^{TL}([R]) - d_i^P \; P_i \,. \qquad [S2]$$

As other studies have pointed out (see e.g. (1)), expressing the equations in terms of absolute numbers of molecules instead of concentrations has the advantage of showing clearly (i) whether the system is growing or not (ii) whether the system exhibits balanced exponential growth. Note that other studies use total mass instead of total numbers (2).

As mentioned in the main text, we simplify the previous equations by considering transcript degradation rates that are identical for each class, and by setting to zero all protein degradation rates (meaning that the model should apply for nutrient conditions with moderate-to-fast growth rates (3)),

$$d_i^m = d \quad \forall i \quad ; \quad d_i^P = 0 \quad \forall i \,.$$

Next, we assume that the biosynthesis fluxes $J_i^{TX}([N])$ and $J_i^{TL}([R])$ are initiation-limited and therefore we describe them by a function of transcription and translation initiation respectively, which we call $\beta_i$ and $\alpha_i$. This is not a trivial assumption as many nonequilibrium models of transcription and translation show the existence of regimes controlled by different limiting steps of the process (initiation, elongation, termination) (4–7). The initiation-limited regime that we consider here applies to regimes of low density of RNA polymerases on genes, and of ribosomes on transcripts, i.e., genes and transcripts are assumed to be far from being saturated with RNA polymerases and ribosomes. Roughly, a low-density regime should generally apply in standard physiological conditions, at least in mRNA translation, which provides some support for the assumption of initiation-limited regimes (7–10).

In order to fully specify the model one needs to provide the dependence of the initiation rates on RNA polymerase and ribosome abundances. In this study, we assume transcription and initiation rates to be proportional to the free concentrations $[N_f]$ of RNA polymerases and $[R_f]$ of ribosomes. In addition, we assume that most regulation happens at the transcriptional level. We restrict our analysis here to identical translation initiation rates $\alpha$ for all genes, while transcription rates $\beta_i$ can be class-dependent. This is a relatively strong assumption, but (i) it is not essential to derive the existence of the growth limiting regime used in the main text and (ii) it is widely employed in some organisms such as *E. coli* (10).

The assumptions in the present section are described mathematically by the following equations:

$$J_i^{TX}([N]) = \beta_0^i [N_f]; \quad J_i^{TL}([R]) = \alpha_0^i [R_f] \,, \qquad [S3]$$

where $\alpha_0^i$ and $\beta_0^i$ are rate constants. These rate constants represent the affinity of ribosomes for mRNAs and RNA polymerases for genes, respectively.

### S2. Derivation of the expression for the mRNA-dependent growth rate

This section derives Eq. [7] of the main text, which relates growth rate to mRNA-dependent translation initiation. It starts with the translation current $J_i^{TL}([R])$ from Eq. [S3], which links the initiation rate $\alpha_i = \alpha_0^i [R_f]$ to the free ribosome concentration. The derivation assumes that all transcripts are identical at the last step and uses the index $i$ for the gene class to provide a general derivation.

It is possible to relate the free ribosomal pool and the the total amount of ribosomes by counting the number of bound ribosomes on each mRNA. For instance, we can evaluate the ribosome density on each transcript by knowing its translation initiation and elongation rates. To this aim, we can exploit nonequilibrium traffic models (4) in a regime of negligible ribosome traffic to estimate the ribosome density (number of ribosomes per codons) on an mRNA belonging to the class $i$ as the ratio between the ribosome initiation and elongation rate $\alpha_i/k_{tl}$ -see (3). Thus,

$$R_{b,i} = \frac{\alpha_0^i}{k_{tl}} L_i [R_f]. \qquad [S4]$$

The total number of bound ribosomes is obtained by summing $R_{b,j}$ over all classes $j$, considering the number of transcript $m_j$ in each class, i.e., $R_b = \sum_j m_j R_{b,j}$. By using Eq. [S4] in this definition, we find

$$[R_b] = \sum_j \frac{\alpha_0^j}{k_{tl}} L_j [m_j][R_f] \,. \qquad [S5]$$

Finally as $[R] = [R_f] + [R_b]$, we can express the free- and bound-ribosome concentrations in terms of the total ribosome concentration as follows,

$$[R_f] = \frac{1}{1 + \sum_j L_j \frac{\alpha_0^j}{k_{tl}}[m_j]}[R] \tag{S6}$$

$$[R_b] = \frac{\sum_j L_j \frac{\alpha_0^j}{k_{tl}}[m_j]}{1 + \sum_j L_j \frac{\alpha_0^j}{k_{tl}}[m_j]}[R]\,. \tag{S7}$$

It is worth noticing that the ratio

$$f_{br} := \frac{\sum_j L_j \frac{\alpha_0^j}{k_{tl}}[m_j]}{1 + \sum_j L_j \frac{\alpha_0^j}{k_{tl}}[m_j]} \tag{S8}$$

represents the fraction of bound ribosomes engaged on the $m$ transcripts. Therefore, we obtain the following expression for the translational current, describing the protein synthesis of a transcript of type $i$,

$$J_i^{TL}([R]) = \alpha_0^i \frac{1}{1 + \sum_j L_j \frac{\alpha_0^j}{k_{tl}}[m_j]}[R] \tag{S9}$$

To simplify our predictions, we assume now that all transcripts have the same length and initiation rate constant. This assumption can be seen as a "mean-field" approximation, where all transcripts follow the average behavior.

$$\alpha_0^i = \alpha_0 \text{ and } L_i = L_p \quad \forall i\ , \tag{S10}$$

where $L_p$ is the typical protein length (in units of codons). Consequently, we can drop the subscript and express the overall translation flux as

$$J^{TL}([R],[m]) = \frac{k_{tl}}{L_p}\frac{[R]}{K_m + [m]} \qquad K_m := \frac{k_{tl}}{L_p \alpha_0}\,. \tag{S11}$$

In this expression, crucial to our central predictions, the protein synthesis rate is a function of both the total amount of ribosomes and of the total mRNA concentration -the latter determining the amount of bound ribosomes.

Finally, we can convert the ribosome concentration into the ribosomal protein concentration. If there are $n_R$ ribosomal protein in each ribosome, then $[R] = \frac{1}{n_R}[P_R]$, leading to $J^{TL}([R],[m]) = \frac{k_{tl}}{L_R}\frac{[P_R]}{K_m+[m]}$, where we defined $L_R := L_p n_R$ that is the total protein length in a ribosome. As discussed in the main text, the ratio $\gamma := \frac{k_{tl}}{L_R}$ represents the inverse of the time needed to translate all the ribosomal proteins making a ribosomes (see also ref. (1)). Thus Eq. [S2] for the protein production of each class $i$ becomes

$$\frac{dP_i}{dt} = \gamma \frac{m_i}{m} P_R \frac{[m]}{K_m + [m]}\,. \tag{S12}$$

This work assumes that, under the approximations made, the number of ribosomal proteins $P_R$ is coupled to the production of all proteins, including their own production. This autocatalysis cycle is responsible for the steady exponential growth and the relationship between the exponential growth rate and the fraction of ribosomal proteins (1, 2). If all mRNAs have the same initiation rates $\alpha$, same elongation rates $k_{tl}$, and length, the ratio $m_i/m$ corresponds to the fraction of ribosomes translating the transcripts of class $i$, $m_i R_{b,i} / \sum m_j R_{b,j} = m_i/m$, which we define as $\chi_i$, the ribosome allocation parameter.

Summing Eq. [S12] over all gene classes gives the total protein production rate

$$\frac{dP}{dt} = \gamma P_R \frac{[m]}{K_m + [m]}\,. \tag{S13}$$

We divide this expression by the total protein number $P$ and obtain Eq. [6] of the main text,

$$\lambda = \gamma \phi_R \frac{[m]}{K_m + [m]}\,, \tag{S14}$$

providing the prediction of this model for the relation between growth rate, ribosome allocation and mRNA concentrations for moderate-to-fast growth rates.

We note that steady-state balanced growth, all protein classes grow at the same rate $\lambda$. Eq. [S12] and Eq. [S14] imply that $\chi_i = \phi_i$, meaning that ribosome allocation corresponds to proteome allocation (2). We use the symbol $\phi_i$ to indicate the number fractions $P_i/P$, although in the literature, this symbol is often used to indicate mass fractions. However, it is immediate within our mean-field approach to convert number fractions into mass fractions using the typical masses of proteins in each sector.

### S3. Argument locating the complex-limited growth regime

This section provides further arguments identifying distinct limiting regimes for growth, and relating the assumption of complex-limited growth made in our model to other studies. It is crucial to understand that our model assumptions are designed to apply for a specific regime of a more general "phase space" of growth limitations (which yet needs to be fully characterized in the literature), and that other studies have focused on other regimes in partial overlap with ours. In the following paragraphs, we will first characterize the limiting regimes accessible by our model, then we will provide a simple argument to support how they are related to a more general phase space, and finally we will discuss how this relates to previous works.

We start by listing the growth regimes accessible to our model. Following Roy *et al.* (1), we characterize a growth limiting regime by a list of all the molecular species that set the total protein production rate $\frac{dP}{dt}$. However, our model consider co-limitation regimes, in which more molecular species can concurrently impose constraints on growth. The equations derived in the previous section show that our model exhibits two limiting regimes:

1. when $[m] \gg K_m$, the total protein production rate becomes a function of the ribosomal proteins $P_R$ only. This is also equivalent to stating that the growth rate $\lambda$ depends only on the ribosome protein function $\phi_R$, or equivalently on the ribosome concentration $[R]$ only (since total protein density is assumed constant and $\phi_R \propto [R]/[P]$). We call this regime the translation limiting regime (TL-LIM) because only quantities related to translation determine total growth;

2. when $[m] \ll K_m$, (more generally when $\frac{[m]}{[m]+K_m}$ is not close to 1), the total protein production rate is a function of both the mRNA concentration $[m]$ and ribosomal protein $P_R$. This is equivalent to stating that the growth rate depends on both mRNA concentration $[m]$ the ribosome protein fraction $\phi_R$, or equivalently on the ribosome concentration $[R]$. We call this regime the complex-formation limiting regime (CF-LIM), referring to mRNA-ribosome complex.

We proceed by placing our model in a wider phase space of growth limitations. The illustration in Fig. 1C of the main text shows the existence of a third growth limiting regime where the total production rate depends solely on the mRNA $m$. We call this regime the transcription limiting regime (TX-LIM). To understand it, we use a toy-model argument based on simple chemical kinetics between two reacting species, and we show how all three regimes emerge in this toy model. Despite its simplicity, we believe this argument provides the essential intuition behind the existence of all three regimes. Let us consider two chemical species $A$ and $B$ in a solution of fixed volume. $A$ and $B$ meet and form a complex $AB$ with a binding constant $k_{CF}$ following mass action. The complex $AB$ produces $P$ with a rate $k_p$, and spontaneously dissociates into $A$ and $B$ simultaneously. The process is represented by the following chain of reactions,

$$A + B \xrightarrow{k_{CF}} AB \xrightarrow{k_p} P + A + B\,. \qquad \text{[S15]}$$

Note that all such reactions are assumed to be *irreversible*, far-from-equilibrium.

Writing down the rate equations corresponding to this process is straightforward following mass action kinetic, provided that one accounts for both the free and the total quantities. For instance, the production rate of the complex $AB$ would be $k_{CF}[A_f][B_f]$ with the subscript $_f$ indicating the free quantities. Defining $K_{AB} = \frac{k_p}{k_{CF}}$, the equations for the dynamics of the product concentration in terms of total quantities read

$$\frac{d[P]}{dt} = \begin{cases} k_p \min\left([A],[B]\right) & [A] \gg K_{AB} \quad OR \quad [B] \gg K_{AB} \\ k_p[A]\frac{[B]}{[B]+K_{AB}} & [A] \ll K_{AB} \quad [B] \approx K_{AB} \\ k_p[B]\frac{[A]}{[A]+K_{AB}} & [A] \approx K_{AB} \quad [B] \ll K_{AB} \\ k_{CF}[A][B] & [A] \ll K_{AB} \quad AND \quad [B] \ll K_{AB}\,. \end{cases} \qquad \text{[S16]}$$

The first case in Eq. [S16] represents the regime of limiting *product* formation. In this case, the product production rate depends either on $[A]$ or $[B]$, depending on which concentration is lower. Consequently, this case splits into two limiting regimes (for $A$ or $B$). All the other cases represent limiting regimes where the product production rate depends on both $[A]$ and $[B]$. This is the case of limiting *complex* formation. The consequences of these different limiting regimes are particularly striking in the last case, where the product production rate depends on $[A][B]$.

If we now interpret $[A]$, $[B]$ and $[P]$ in our toy model as representative, respectively, of mRNA $[m]$, ribosomes $[R]$ and proteins $[P]$, we can map conceptually these exact results to the biologically relevant case of protein production. We are of course aware that the context of realistic biosynthesis includes multiple complications, one of which being the fact that mRNAs can form a complex with multiple ribosomes, which may also interfere destructively with one another. Such complications would require a more complex model. Yet, we believe that our toy model provides the correct intuition for the presence of three main limiting regimes: two regimes limited by products (transcription TX and translation TL), and a complex-formation (CF) limiting regime.

Finally, these simple arguments also help us to compare our study with previous work (1, 11, 12). Crucially, previous studies mainly focus on product-limited regimes. Lin and Amir (12) explore a one-dimensional phase space where protein-to-DNA ratio is the control variable. This allows them to go from ribosome-centric exponential growth to a regime where mRNAs are saturated and exponential growth is supported by the RNA polymerase autocatalytic cycle, as well as an additional regime where genes are saturated, and exponential growth is not possible. Specifically, Lin and Amir (12) state that a transcription-limited regime emerges when mRNA are spatially saturated by ribosomes - therefore, increasing the number of ribosomes does not increase translation fluxes. Within our model, the regime can arise by relaxing the assumption of initiation-limited translation flux $J_i^{TL}([R])$. Dropping this assumption makes the model access regimes of high ribosome density on mRNA, which corresponds to the condition of mRNA saturation explored in (12). Roy and coworkers (1) also explore several saturation regimes, but do not address complex formation. A full description of growth limitation beyond the initiation-limited assumption of the translation flux is beyond the scope of this work. Our main contribution is to focus on the regime limited by complex formation and to show that it can explain experimental trends that are not explained by any product-limited regime (Figs. 2-4 of the main text).

### S4. Different predictions between a CF-LIM and a TX-LIM regime

This section compares the main predictions of our model in a transcription-limited (TX-LIM) regime with the complex-formation limited (CF-LIM) regime discussed in the main text and other sections of this SI Appendix. The comparison is conducted without the optimization procedure explained in the main text (which is inessential for comparing the two regimes).

**Mathematical definition of the transcription limited (TX-LIM) regime.** We define the transcription-limited regime is defined as the case where the protein production rate depends solely on mRNA. Note that this is a specific case of the more general Eq. [S2]. Using the same notation, the transcription-limited regime can be expressed as

$$\frac{dP_i}{dt} = m_i J^{TL} \,. \tag{S17}$$

As stated earlier, $J^{TL}$ represents the ribosome flux or protein synthesis rate per mRNA. However, in the transcription-limited (TX-LIM) regime, it becomes a constant value that is independent of the total ribosome concentration. In this regime, initiation is no longer the limiting step, as assumed in Eqs. [S3]. The reason for a constant $J^{TL}$ is not relevant for the subsequent analysis. This situation may occur when the protein synthesis rate on a single mRNA is no longer affected by the availability of free ribosomes. This scenario can arise when all transcripts are saturated, indicating that translation is constrained by ribosome interference instead of initiation. Moreover, we assume that $J^{TL}$ is identical for all proteins $P_i$ to simplify the analysis.

To complete the model, we need an equation for the dynamics of mRNA $m_i$. In the following analysis, we consider two choices for the dynamics of mRNA: (i) RNA polymerase-limited mRNA transcription, which was presented both in the main text and previously in this SI (initiation-limited transcription), and (ii) DNA-limited mRNA transcription, where the transcription flux $J^{TX}$ in Eq. [S1] becomes a constant that is independent of RNA polymerases, using the same reasoning that led us to consider $J^{TL}$ as independent of ribosomes. We also note that these cases have also been investigated in (12). We will highlight the difference with the CF-LIM regime (explored in the main text) below.

**Transcription limited (TX-LIM) model of biosynthesis with RNA polymerase-limited transcription.** We write the dynamical equations of the transcript $m_i$ and protein $P_i$ for each gene class $i$:

$$\frac{dm_i}{dt} = \frac{k_{tx}}{L_N}\, f_{bn}\, \omega_i\, P_N - d\, m_i \,. \tag{S18}$$

$$\frac{dP_i}{dt} = m_i J^{TL} \tag{S19}$$

In the first equation, we have assumed RNA polymerase-limited transcription, which is similar to Eq. [S1] used in the previous sections (also converting RNA polymerases complexes to the number of protein within. $\frac{k_{tx}}{L_N}$, $\omega_i$ and $f_{bn}$ are respectively the inverse typical time needed to transcribe RNAP genes (the ratio of a typical transcription elongation rate and total RNAP gene length), the fraction of bound RNA polymerases translating genes of type $i$ and $f_{bn}$ the overall fraction of RNA polymerases bound to genes. The second equation follows from the previous paragraph, where we recall that $J^{TL}$ is a constant parameter that is independent of ribosome or mRNA concentration, unlike equation [S2].

Next, we derive an expression for the growth rate in this regime by assuming quasi-steady-state in $m_i$, which yields

$$m_i = \frac{k_{tx}}{dL_N}\, f_{bn}\, \omega_i\, P_N \,. \tag{S20}$$

By plugging this expression into the equation for the protein dynamics, we find that

$$\frac{dP_i}{dt} = \gamma_N\, f_{bn}\, \omega_i\, P_N \,, \qquad \gamma_N = \frac{k_{tx} J^{TL}}{d\, L_N} \,. \tag{S21}$$

Therefore, the production of all proteins is coupled to RNA polymerase abundances in this regime. In particular, RNA polymerase proteins are involved in an autocatalytic cycle and exponential growth. Summing over all protein classes in equation [S21] gives the total protein production rate

$$\frac{dP}{dt} = \gamma_N\, f_{bn}\, P_N \,. \tag{S22}$$

We divide this expression by the total protein number to obtain the exponential growth rate

$$\lambda = \gamma_N\, f_{bn}\, \phi_N \,. \tag{S23}$$

Finally, we also sum over all transcript classes in equation [S20] and obtain the total mRNA concentration

$$[m] = \frac{k_{tx}}{d\, L_N}\, f_{bn}\, \phi_N [P]\,. \tag{S24}$$

By putting together the last two equations, we obtain

$$\lambda = [m]\, \frac{J^{TL}}{[P]}\,. \tag{S25}$$

Figure S1 compares the transcription-limited regime with RNA polymerase-limited transcription and the complex-formation limited (CF-LIM) regime using Eq. [S25] for the TX-LIM regime and the growth rate equation for the CF-LIM regime, which can be found in Eq. [8] of the main text and in Eq. [S44] of this SI Appendix.

It is worth noting that we compare the CF-LIM and TX-LIM regimes for fixed parameter values without the growth rate optimization discussed in the main text.

Fig. S1B indicates that the growth rate increases with total mRNA concentration in both regimes, although it increases linearly only in the TX-LIM regime. Figure S1C shows that increasing ribosome levels has no effect in the TX-LIM regime but matters in the CF-LIM regime.

**Transcription limited (TX-LIM) model of biosynthesis with DNA-limited transcription.** We write the equations of the transcript and protein dynamics for each gene class-$i$:

$$\frac{dm_i}{dt} = g_i J^{TX} - d\, m_i\,, \tag{S26}$$

$$\frac{dP_i}{dt} = m_i J^{TL}\,. \tag{S27}$$

Both $J^{TX}$ and $J^{TL}$ are constant parameters independent of the dynamical variables $m_i$ and $P_i$. In this case, the dynamics is highly dependent on the dynamical behavior of $g_i$, i.e., DNA replication. A full treatment of this case that must including a detailed model of DNA replication is beyond the scope of this work. Nonetheless, we mention a number of key points.

If $g_i$ is constant in time, $m_i$ also reaches a constant value given by $m_i = g_i \frac{J^{TX}}{d}$, assuming quasi-steady-state. Therefore, protein are produced linearly, not exponentially, as $\frac{dP_i}{dt}$ is a constant in the equation [S27]. The linear growth rate of total protein amount takes the expression:

$$\lambda_l = g\, \frac{J^{TL} J^{TX}}{d}\,. \tag{S28}$$

If $g_i$ changes in a time, we may still assume quasi-steady-state in the transcript dynamics, that is, $m_i(t) = g_i(t)\, \frac{J^{TX}}{d}$. We obtain the following equation for the total protein content:

$$\frac{dP}{dt} = g(t)\, \frac{J^{TL} J^{TX}}{d}\,. \tag{S29}$$

Therefore, a single cell does not follow a simple growth law in this case, as for example linear or exponential. At the bulk population level, we may associate a growth rate to this model by considering the time it takes to duplicate the proteome and assume that this is equal to the division time. Defining $\tau_d$ as the division time, we use the equation above to find the time it takes to duplicate the total protein content:

$$2P_0 - P_0 = \int_0^{\tau_d} \frac{dP}{dt} dt = \int_0^{\tau_d} g(t)\, \frac{J^{TL} J^{TX}}{d} dt = \tau_d\, \frac{\bar{g}}{P_0}\, \frac{J^{TL} J^{TX}}{d}\,. \tag{S30}$$

where $\bar{g}$ is the time average total DNA content along the cell cycle. Defining the bulk growth rate as $\lambda = \frac{\ln 2}{\tau}$, we find

$$\lambda = \frac{\bar{g}}{P_0}\, \frac{J^{TL} J^{TX}}{d}\,. \tag{S31}$$

The term $\frac{\bar{g}}{P_0}$ is close to the average DNA concentration along the cell cycle, as can be seen by multiplying and dividing by the volume $V_0$. The same equation can be re-written in terms of the time-averaged mRNA content,

$$\lambda = \frac{\bar{m}}{P_0}\, J^{TL}\,. \tag{S32}$$

Once again, $\frac{\bar{m}}{P_0}$ is approximately equal to the average mRNA concentration throughout the cell cycle. We multiply and divide this expression by the average volume and total protein amount throughout the cell cycle to obtain

$$\lambda = r\, \frac{\overline{[m]}}{\overline{[P]}}\, J^{TL}\,. \tag{S33}$$

Here, $[\bar{m}]$ and $[\bar{P}]$ represent the average mRNA and protein concentration throughout the cell cycle, respectively, while $r = \frac{\bar{P}}{P_0}$ is the ratio of the average total protein amount to the initial protein amount. The exact value of this ratio depends on the specific cell cycle model, but it should be a term of order 1. Therefore, with a suitable reinterpretation of the parameters, the TX-LIM model with DNA-limited transcription can exhibit the same dependence on mRNA concentration as the TX-LIM model with RNA polymerase-limited transcription.

### S5. The origin of the linear relation between mRNA and RNA polymerase concentrations in the CF-LIM regime

Back to the CF-LIM regime, this section concerns the derivation of Eq. [12] of the main text. We start by expressing the transcript production rate per gene $J_i^{TX}$ as a function of the total concentration $[N]$ of RNA polymerases. Following the same steps as in the previous section for the translation current, we find an expression for the concentrations $[N_f]$ and $[N_b]$ of free and of bound RNA polymerase respectively, in terms of the total RNA polymerase concentration $[N]$, the transcription initiation rate $\beta_i = \beta_0^i [N_f]$, the transcription elongation rate $k_{tx}$, and the gene copy number concentration $[g_i]$ of the gene $i$. We obtain the following equations,

$$[N_f] = \frac{1}{1 + \sum_j L_{g_j} \frac{\beta_0^j}{k_{tx}} [g_j]} [N] \qquad \text{[S34]}$$

$$[N_b] = \frac{\sum_j L_{g_j} \frac{\beta_0^j}{k_{tx}} [g_j]}{1 + \sum_j L_{g_j} \frac{\beta_0^j}{k_{tx}} [g_j]} [N] \,, \qquad \text{[S35]}$$

where, in analogy with the case of translation, we can define $f_{br}$ as the fraction of bound RNA polymerase, *i.e.*, those that are actively transcribing,

$$f_{bn} := \frac{\sum_j L_{g_j} \frac{\beta_0^j}{k_{tx}} [g_j]}{1 + \sum_j L_{g_j} \frac{\beta_0^j}{k_{tx}} [g_j]} \,. \qquad \text{[S36]}$$

Contrary to the case of translation, we consider a general scenario where transcription initiation rates vary from gene to gene, to enable the model to describe the expression of different protein categories at varying levels. Therefore, the expression of the transcription current $J_i^{TX}$ is not simplified in this case. However, we express it in a way that makes it easier to understand its physical meaning,

$$J_i^{TX}([N], [g_i]) = \beta_0^i \frac{1}{1 + \sum_j L_{g_j} \frac{\beta_0^j}{k_{tx}} [g_j]} [N] = \frac{k_{tx}}{L_{g_i}} \, \frac{\omega_i}{[g_i]} \, f_{bn} \, [N] \,. \qquad \text{[S37]}$$

Let us discuss the interpretation of the term $\omega_i$ isolated in the previous equations. We have defined

$$\omega_i := \frac{L_{g_i} \beta_0^i [g_i]}{\sum_j L_{g_j} \beta_0^j [g_j]} \,, \qquad \text{[S38]}$$

which can be interpreted as the fraction of active RNA polymerases that is transcribing gene of type $i$. In other words, $\omega_i$ represents the allocation parameter of RNA polymerase transcribing gene $i$, which depends on the promoter strength $\beta_0^i$.

Having defined $\omega_i$, Eq. [S1] takes the form

$$\frac{dm_i}{dt} = \frac{k_{tx}}{L_{g_i}} \, \omega_i \, f_{bn} \, N - d \, m_i \,. \qquad \text{[S39]}$$

As mRNA degradation rates are fast compared to changes of the production rate, transcripts are often assumed to be in steady state (1, 12). We also follow this approach and write,

$$m_i = \frac{k_{tx}}{d \, L_{g_i}} \, \omega_i \, f_{bn} \, N \,, \qquad \text{[S40]}$$

which links the amount of RNA polymerase and mRNAs. After simplifying the equation by considering that all genes have the same length $L_g$, and by summing the previous equation on all sectors one obtains

$$m = \frac{k_{tx}}{d \, L_g} \, f_{bn} \, N \,. \qquad \text{[S41]}$$

Finally, we convert the amount of RNA polymerases $N$ into the total number of proteins that make up the RNA polymerases, which is given by $N = P_N \frac{L_g}{L_N}$, where $L_N$ represents the total number of amino acids in an RNA polymerase complex (we assume that gene regulation implements perfect stoichiometry). Thus, we obtain Eq. [12] of the main text:

$$[m] = \frac{\gamma_{tx}}{d} \, f_{bn} \, [P_N], \qquad \gamma_{tx} := k_{tx}/L_N \,. \qquad \text{[S42]}$$

### S6. The relation between growth rate and RNA polymerase proteome allocation

We can use Eq. [S42] in conjunction with Eq. [S14] to determine the relationship between the growth rate and the RNA polymerase fraction $\phi_N$, as shown in Eq. [14] of the main text. To accomplish this, we first need to express the concentration $[P_N]$ in terms of the fraction of RNA polymerases in the proteome and the total protein concentration: $[P_N] = \phi_N[P]$, where $[P]$ is the total protein concentration. We can rewrite equation [S42] as

$$[m] = \frac{k_{tx}}{d\, L_N}\, f_{bn}\, \phi_N [P]\,. \tag{S43}$$

Substituting the expression for $[m]$ from Eq. [S43] into Eq. [S14] yields the expression given in Eq. [14] of the main text,

$$\lambda = \gamma\phi_R \frac{\phi_N}{\phi_N + \epsilon^{-1}}\,. \tag{S44}$$

where

$$\epsilon := \frac{\Gamma[P]}{K_m}\,. \qquad \Gamma = \frac{k_{tx}}{d\, L_N}\, f_{bn}\,, \tag{S45}$$

The above considerations imply that both $\epsilon$ and $\Gamma$, which are related to the RNA polymerase pool, can be interpreted as a transcriptional efficiency. Specifically, $\Gamma$ represents the amount of mRNA concentration added per unit of RNA polymerase concentration, while $\epsilon$ is slightly more complex to interpret due to the presence of the parameter $K_m$. As described in the main text, $\epsilon$ combines a purely transcriptional "supply" term $\Gamma$ with a purely translational "demand" term $K_m$, which represents the typical amount of transcripts required by translation. Therefore, this parameter may be interpreted as a supply-demand trade-off or the mRNA transcription efficiency/capacity "normalized" by the amount of mRNA needed.

### S7. Equivalence between ribosome, RNA polymerase and proteome allocations

By dividing Eq. [S40] by Eq. [S41] we obtain that $\chi_i = \omega_i$, i.e. that RNA polymerase and ribosome allocations are identical in the quasi-state-state approximation. As explained previously, in steady-state balance growth, ribosome allocation $\chi_i$ is also equal to proteome allocation $\phi_i$, which allows us to write,

$$\omega_i = \chi_i = \phi_i\,. \tag{S46}$$

We should keep in mind that $\omega_i$ is an external parameter that represents the fraction of RNA polymerases transcribing gene $i$, i.e., the allocation of RNA polymerases. Biologically, $\omega_i$ depends on both the gene copy number $g_i$ and the promoter strength of a gene of type $i$ for RNA polymerases. It is worth noting that this equation is equivalent to stating that gene-weighted relative promoter activities are equal to mRNA fractions, which are equal to protein fractions - a central result in the work of Balakrishnan and coworkers (10). Although we provide a mechanistic and quantitative link between these three quantities, we believe it is often conceptually helpful to consider $\omega_i$ as a primary causal parameter of the framework. The cell may choose to allocate RNA polymerases using different mechanisms of transcriptional regulation.

In the event of a change in RNA polymerase allocation due to a nutrient shift or another environmental change, the transformation $\Omega \to \Omega^{'}$ can be used with our framework to calculate the relative changes in growth rate, transcript composition, and proteome composition. For simplicity (see the main text for a discussion) we often assumed here that cells regulate resource allocation to achieve an optimal growth regime and search for the RNA polymerase allocation assignment $\Omega^*$ that maximizes the growth rate under different perturbations. Since such allocation reflects proteome composition at steady-state, this is the same as looking for the proteome composition $\Phi = \{\phi_1, \ldots, \phi_i, \ldots, \phi_S\}$ that maximizes the growth rate.

It is also worth noting that Eq. [S46] may not be valid if we drop certain assumptions, such as a homogeneous transcript degradation rate or identical translation initiation rate for all protein classes. Specifically, we reconsider the relationship between protein fraction and RNA polymerase allocation fraction when predicting the growth rate reduction due to overexpression of unnecessary protein under different transcript degradation rates (see section S10 and SI Fig. S9).

### S8. The fraction of bound RNA polymerases is a function of a single dimensionless parameter

This section presents a simplified expression for the fraction of bound RNA polymerase, which depends on a single compound parameter. This expression is significant for two reasons: (i) in deriving the primary outcomes of our work, we suppose that the fraction of bound RNA polymerases is close to 1, and this expression clarifies this regime, and (ii) the expression demonstrates how to surpass this assumption and to what extent it affects the model's predictions.

We start by rewriting Eq. [S36] for the fraction of bound RNA polymerases. We assume that each gene has a length of $L_g$. Since gene length heterogeneity can be easily absorbed by redefining the pseudo-promoter strength $\beta_0^j$ (without loss of generality), we obtain

$$f_{bn} = \frac{\sum_j \beta_0^j [g_j]}{\frac{k_{tx}}{L_g} + \sum_j \beta_0^j [g_j]}\,. \tag{S47}$$

Throughout this study, we kept the values for the gene copy number $g_i$ fixed. We find it convenient to explicitly write out the volume rather than the concentration,

$$f_{bn} = \frac{\sum_j \beta_0^j \, g_j}{\left(\frac{k_{tx}}{L_g}\right) \, V + \sum_j \beta_0^j \, g_j} \, . \quad \text{[S48]}$$

In the above equation, the quantity $\sum_j \beta_0^j \, g_j$ represents the overall promoter activity, with higher values corresponding to higher fractions of RNA polymerases specifically bound to promoter regions of the genome. Conversely, the term $(k_{tx}/L_g) \, V$ accounts for the factors that decrease the binding of RNA polymerases to DNA, including overall DNA dilution (volume $V$) and the rescaled elongation rate $(k_{tx}/L_g)$. Faster transcript elongation leads to quicker unbinding of RNA polymerases. Therefore, the ratio $(V k_{tx}/L_g)/(\sum_j \beta_0^j \, g_j)$ is a relevant parameter for setting the fraction of bound RNA polymerases.

It is even more convenient to isolate the promoter activity for all genes that belong to the protein class $Q$, which has a proteome fraction that remains constant across growth conditions. By following this approach, we can isolate possible changes in $f_{bn}$ only through the activity of genes belonging to other protein classes. By isolating the protein class $Q$, we obtain the following expression:

$$f_{bn} = \frac{1}{1 + (1 - \phi_Q) \, Z_g} \, , \qquad Z_g = \frac{(k_{tx}/L_g) \, V}{\sum_{j \neq Q} \beta_0^j g_j} \, . \quad \text{[S49]}$$

The parameter $Z_g$ encodes the global activity of the promoters in terms of mechanistic parameters. In particular, if $Z_g = 0$, then $f_{bn} = 1$. Therefore, assuming that $f_{bn} \approx 1$ is equivalent to stating $Z_g \approx 0$. We derived the equation above using the fact that $\phi_Q = \omega_Q = \frac{\sum_{j \in Q} \beta_0^j g_j}{\sum_j \beta_0^j g_j}$, as discussed in previous sections of this SI Appendix.

Eq. [S49] highlights a challenge for the existence of a sustainable balanced growth steady-state. If we assume that $V$ is proportional to $P$ and that the gene copy numbers $g_j$ remain constant, then $f_{bn}$ must decrease as the cell grows. This, in turn, affects the overall mRNA concentration (as per Eq. [S41]) and eventually protein production (as per Eq. [S12]). To avoid this issue, one should explicitly consider DNA replication and the fact that gene copy numbers $g_j$ increase over time. However, adding an explicit model of DNA replication and the cell cycle is beyond the scope of this work, because it would require endowing the framework with an explicit cell-cycle model. Nevertheless, we argue that in many practical situations, $f_{bn}$ would exhibit little variation, and assuming $f_{bn} \approx 1$ (or equivalently, $Z_g \approx 0$) should be a reasonable approximation. Specifically,

1. if $Z_g \approx 0$, changes in the volume have little effect on $f_{bn}$, which remains close to 1; indeed, some studies (13, 14) have argued that the fraction of bound RNA polymerase is always close to 1;

2. if all $g_j$ were to increase continuously with time, their trend would at least party cancel the continuous increase in volume, leaving $f_{bn}$ fairly constant;

3. if all $g_j$ increase in a step-wise manner, $f_{bn}$ would change, but in an oscillatory fashion.Therefore, one could interpret Eq. [S49] as a time average (fixing $f_{bn}$ to a constant value). We imply a time average when comparing our model to bulk data.

Other studies, both in budding yeast (15) and in *E. coli* (16) have argued in favor of a fraction of bound RNA polymerase (our $f_{bn}$) close to 50%. In this case $Z_g \approx 1$ and there may be some consequences. We explored this regime in the case of the cost of unneeded proteins, as detailed below.

### S9. The RNA polymerase autocatalytic cycle in the CF-lim regime

Roy and coworkers (1) have shown how one can derive growth laws from the mass and time-scale budget of the autocatalitic cycles involving key players such as ribosomes, RNA polymerase, tRNA, rRNA. Our model mainly describes ribosomes, mRNA and RNA polymerase, and the latter are nearly equivalent if one assumes that rapid mRNA degradation leads mRNA to a quasi-steady state (see below for model variants including rRNA). It is simple to show that in the TL-LIM regime the ribosome autocatalytic cycle is equivalent to that of the model by Roy and coworkers. More generally, mRNA and ribosomes are co-limiting in our model, and following the resulting autocatalytic cycle leads to Eq. [S14].

It is more interesting to follow the RNA polymerase autocatalytic cycle, which in our model involves RNA polymerase (translated by ribosomes), mRNA, and ribosomes (transcribed by RNA polymerases). Cosidering total mRNA first, we can write

$$\frac{dm}{dt} = \frac{k_{tx}}{L_g} f_{bn} N - dm \, . \quad \text{[S50]}$$

Since for steady exponential growth $\frac{dm}{dt} = \lambda m$, we obtain

$$[m] = \Gamma [N] = \Gamma \, [P] \, \phi_N \, , \quad \text{[S51]}$$

setting a relationship between mRNA and RNA polymerase concentration, with

$$\Gamma = \frac{k_{tx}}{L_g (d + \lambda)} f_{bn} \, , \quad \text{[S52]}$$

which depends only weakly from $\lambda$ since $\lambda \ll d$. We assumed constant total protein concentration $[P] = P/V$, so that $\phi_x = \frac{[X]}{[P]}$. Turning now to the linked cycles of ribosomes and RNA polymerase, we get

$$\frac{dR}{dt} = \gamma\omega_R R \frac{[N]}{\frac{K_m}{\Gamma} + [N]} \ , \tag{S53}$$

and

$$\frac{dN}{dt} = \gamma\omega_N R \frac{[N]}{\frac{K_m}{\Gamma} + [N]} \ , \tag{S54}$$

where $\omega_i$ represents RNA polymerase allocation towards class $i$. For steady-exponential growth we can impose that $\frac{dN}{dt} = \lambda N$, $\frac{dR}{dt} = \lambda R$, and $\omega_i = \phi_i$. Using these conditions, we obtain

$$\lambda = \gamma\phi_R \frac{\phi_N}{\epsilon^{-1} + \phi_N} \tag{S55}$$

and

$$\lambda = \gamma\phi_N \frac{R}{N} \frac{\phi_N}{\epsilon^{-1} + \phi_N} \ , \tag{S56}$$

with $\epsilon := \frac{\Gamma[P]}{K_m}$. Hence, the RNA polymerase autocatalytic cycle leads to the definition of the supply-demand parameter $\epsilon$, and both the equations for RNA polymerase and ribosomes lead to a single growth law, equivalent to Eq. [S14], but describing the link between growth rate and the fraction of RNA polymerase-associated proteins $\phi_N$. Thanks to this link, our model predicts how the ribosomal proteome fraction (under growth optimization) changes as transcription is inhibited (Fig. 4 of the main text), which is not the case in the model by Roy and coworkers(1).

### S10. Unnecessary proteins

This section discusses the calculations leading to the predictions concerning the physiological role of unnecessary proteins within the model.

**Growth rate reduction with unnecessary proteins in the TL-LIM and CF-LIM regime.** This section discusses the impact of unnecessary proteins on growth rate reduction in the TL-LIM and CF-LIM regime. Specifically, let $U$ be the protein class that corresponds to unneeded proteins, and assume there are $g_U$ genes in this class. The fraction of RNA polymerases transcribing this class is then $\omega_U(g_U)$. However, this sector competes for resources with other sectors, which can negatively affect the growth rate $\lambda$ compared to the situation where the unnecessary genes are absent (referred to as "WT").

To obtain the relative growth rate $\lambda/\lambda^{WT}$ in presence of the unnecessary proteins, we use Eq. [S14], obtaining

$$\frac{\lambda}{\lambda^{WT}} = \frac{\phi_R}{\phi_R^{WT}} \frac{[m]}{[m]^{WT}} \frac{1 + \frac{K_m}{[m]^{WT}}}{\frac{[m]}{[m]^{WT}} + \frac{K_m}{[m]^{WT}}} \ . \tag{S57}$$

The limit $\frac{K_m}{[m]^{WT}} \to 0$ corresponds to the translation limiting regime (TL-LIM), while the limit $\frac{K_m}{[m]^{WT}} \to \infty$ corresponds to the complex-formation limiting regime (CF-LIM), as discussed above. We thus begin by computing the ratios $\frac{\phi_R^{WT}}{\phi_R^{WT}}$ and $\frac{[m]}{[m]^{WT}}$ in the CF-LIM and TL-LIM cases studied in the main text.

***Changes in the ribosomal protein fraction ratio*** $\frac{\phi_R}{\phi_R^{WT}}$**.** We name $\phi_i^{WT}$ the protein fractions in the absence of the unneeded sector, and we assume that each protein fraction $\phi_i$ is affected in the same way by the presence of the unneeded protein sector $\phi_U$. The proteome allocations $\phi$ in presence of unnecessary protein expression can be written as $\phi_i = c\phi_i^{WT}$ for all $i \neq U, Q$. The normalization condition is $\sum_i \phi_i = 1$, and we re-write this condition by explicitly considering the sector $\phi_U$ and the housekeeping sector $\phi_Q$. We obtain $c\sum_{i\neq U,Q} \phi_i^{WT} + \phi_U + \phi_Q = 1$. Then $c = 1 - \frac{\phi_U}{1-\phi_Q}$ as $\sum_{i\neq U,Q} \phi_i^{WT} = 1 - \phi_Q$. Therefore:

$$\frac{\phi_i}{\phi_i^{WT}} = \left(1 - \frac{\phi_U}{1 - \phi_Q}\right) \ . \tag{S58}$$

In particular, the ribosome protein fraction $\frac{\phi_R}{\phi_R^{WT}}$ is also affected in the same way, leading to Eq. [11] of the main text.

***Changes in the mRNA concentration ratio*** $\frac{[m]}{[m]^{WT}}$**.** Next, we derive the changes in mRNA content $[m]/[m]^{WT}$. Combining Eq. [S40] with the considerations made above gives an expression for the transcripts of sectors $i$ and $U$,

$$m_i = \begin{cases} \frac{1}{d}\frac{k_{tx}}{L_g}\omega_i f_{bn} N & i \neq U \\ \frac{1}{d_U}\frac{k_{tx}}{L_g}\omega_U f_{bn} N & i = U \ . \end{cases} \tag{S59}$$

Summing on all $i$ gives the total mRNA $m = \frac{k_{tx}}{L_g} f_{bn} N \left( \frac{1}{d} \sum_{i \neq U} \omega_i + \frac{\omega_U}{d_U} \right)$. To further simplify this expression, we re-write the concentration in terms of the protein fraction and we use the fact that $\sum_{i \neq U} \omega_i = 1 - \omega_U$. The total mRNA content then reads

$$[m] = \frac{\gamma_{tx}}{d} [P] \left[ 1 - \left( 1 - \frac{d}{d_U} \right) \omega_U \right] f_{bn} \phi_N \, , \qquad \text{[S60]}$$

where we recall that $\gamma_{tx} = k_{tx}/L_N$ and $[N] = \phi_N \frac{L_g}{L_N} [P]$. Note that both $f_{bn}$ and $\phi_N$ can also depend on quantities linked to sector $U$. In particular, if $\phi_N$ behaves as any other sector, it will obey Eq. [S58].

Eq. [S60] for the total mRNA content is explicitly dependent on $\omega_U$. However, this is not very practical for experimental comparison as the fraction of RNA polymerases transcribing sector $U$ is not easily measurable. Fortunately, the model permits us to express it in terms of the protein fraction $\phi_U$ instead. At steady state, we have $\phi_U = \frac{m_U}{m}$, which yields the following expression

$$\phi_U = \frac{\frac{d}{d_U} \omega_U}{\left[ 1 - \left( 1 - \frac{d}{d_U} \right) \omega_U \right]} \, . \qquad \text{[S61]}$$

Note that if $d_U = d$ then $\omega_U = \phi_U$, while if $d_U \neq d$, the protein fraction $\phi_U$ is no longer equal to the RNA polymerase fraction $\omega_U$. We can invert this relationship to get the conversion between these two quantities

$$\omega_U = \frac{\phi_U}{\frac{d}{d_U} + \phi_U \left( 1 - \frac{d}{d_U} \right)} . \qquad \text{[S62]}$$

Finally, from Eqs. [S58], [S60] and [S62] we obtain the following expression for the relative ratio of mRNA content between the unperturbed situation and the presence of unnecessary proteins,

$$\frac{[m]}{[m]^{\mathrm{WT}}} = \left[ 1 - \frac{\phi_U \left( \frac{d_U}{d} - 1 \right)}{1 + \phi_U \left( \frac{d_U}{d} - 1 \right)} \right] \frac{f_{bn}}{f_{bn}^{\mathrm{WT}}} \frac{\phi_N}{\phi_N^{WT}} \, . \qquad \text{[S63]}$$

The behavior of the ratio $\frac{\phi_N}{\phi_N^{WT}}$ depends on whether the RNA polymerase protein fraction is part of the housekeeping protein sector $Q$ or not. To recapitulate both cases, we write

$$\frac{[m]}{[m]^{\mathrm{WT}}} = \begin{cases} \left[ 1 - \frac{\phi_U \left( \frac{d_U}{d} - 1 \right)}{1 + \phi_U \left( \frac{d_U}{d} - 1 \right)} \right] \left( 1 - \frac{\phi_U}{1 - \phi_Q} \right) \frac{f_{bn}}{f_{bn}^{WT}} \, , & \phi_N \notin Q \\ \left[ 1 - \frac{\phi_U \left( \frac{d_U}{d} - 1 \right)}{1 + \phi_U \left( \frac{d_U}{d} - 1 \right)} \right] \frac{f_{bn}}{f_{bn}^{WT}} \, , & \phi_N \in Q \, . \end{cases} \qquad \text{[S64]}$$

The ratio $\frac{f_{bn}}{f_{bn}^{WT}}$, is given by the expression

$$\frac{f_{bn}}{f_{bn}^{WT}} = \frac{1 + (1 - \phi_Q) \, Z_g^{WT}}{1 + \left( 1 - \frac{\omega_U}{\omega_U^{max}} \right) (1 - \phi_Q) \, Z_g} \, , \qquad \text{[S65]}$$

where $\omega_U^{max}$ reflects the fact that the size of protein class $Q$ stays constant, setting a bound to the fraction of RNA polymerases transcribing sector $U$ genes. We will prove in the next section that $\omega_U^{max} = (1 - \phi_Q)/[1 - \phi_Q(1 - d/d_U)]$. Note that the presence of the unneeded gene increases the overall fraction of bound RNA polymerases (that is, if $\omega_U > 0$ the ratio $\frac{f_{bn}}{f_{bn}^{WT}}$ becomes greater than 1). Finally, we can re-write Eq. [S65] in terms of $\phi_U$ by using Eq. [S62] (see the next subsection).

We note that the process by which RNA polymerase is recruited to the genome in presence of an unneeded protein is not relevant for the global transcriptional capacity if $f_{bn}^{WT} \approx 1$ ($Z_g \approx 0$), i.e. if most RNA polymerases were already bound. In this scenario we expect that total mRNA concentration generally decreases in the presence of the unneeded protein. However, in the alternative scenario $f_{bn}^{WT} \approx 0.5$ ($Z_g \approx 1$), additional recruitment of RNA polymerases can become important and the total mRNA concentration can also *increase* in presence of unneeded proteins.

Specifically, we can rewrite Eq. [S63] as

$$\frac{[m]}{[m]^{\mathrm{WT}}} = \left[ 1 - \left( 1 - \frac{d}{d_U} \right) \omega_U \right] \frac{f_{bn}}{f_{bn}^{\mathrm{WT}}} \frac{\phi_N}{\phi_N^{WT}} \, , \qquad \text{[S66]}$$

where $\omega_U$, the fraction of RNA polymerase allocated to the unneeded genes, is a function of $g_U$ and $\beta_0^U$. Hence, $g_U$ and $\beta_0^U$ affect the change in total mRNA only through $\omega_U$.

The ratio $\frac{\phi_N}{\phi_N^{WT}}$ depends on whether $\phi_N$ belongs to the Q-sector. In general, we assume that it either stays constant or it decreases as $\frac{\phi_N}{\phi_N^{WT}} = 1 - \frac{\phi_U}{1 - \phi_Q}$. The ratio $\frac{f_{bn}}{f_{bn}^{WT}}$, is given by Eq. [S65], where $\omega_U^{max}$ reflects the fact that the proteome fraction relative to the $Q$ sector stays constant, setting a bound to the fraction of RNA polymerases transcribing sector-$U$ genes. The presence of the unneeded gene increases the overall fraction of bound RNA polymerase because if $\omega_U > 0$ the ratio $\frac{f_{bn}}{f_{bn}^{WT}}$ becomes greater than 1. Therefore, the increase in $\frac{f_{bn}}{f_{bn}^{WT}}$ might cause an increase in the total mRNA concentration.

Fig. S10 shows the change in mRNA concentration as a function of the fraction of unneeded proteins for different vaules of $Z_g$, also considering whether or not RNA polymerase is maintained homeostatically upon the perturbation (i.e., $\phi_N$ belongs to the Q sector, see below). We find that $[m]$ may increase if $\phi_N$ belongs to the Q sector and a sufficiently high fraction of RNA polymerase is not specifically bound to promoters or transcribing, and can therefore be recruited by the newly introduced unnecessary gene(s). We further investigate quantitatively this phenomenon in the final subsection of this section.

The detailed procedure to derive Eq. [S65] is presented in a following subsection ("The dependency of the growth rate ratio $\frac{\lambda}{\lambda^{WT}}$ on the gene copy number $g_U$").

***Limit regimes of $\frac{\lambda}{\lambda^{WT}}$.*** We will now derive the expressions for $\frac{\lambda}{\lambda^{WT}}$ in the TL-LIM and CF-LIM cases used in the main text. We will consider two scenarios: one where RNA polymerase proteins belong to the $Q$ sector ($\phi_N \in Q$, which stays constant), and another where they are not part of it ($\phi_N \notin Q$, which can vary). In the regime where translation limits growth, we take the limit $\frac{K_m}{[m]^{WT}} \to 0$ in Eq. [S57] and obtain

$$\frac{\lambda}{\lambda^{WT}} = \begin{cases} \left(1 - \frac{\phi_U}{1-\phi_Q}\right) , & \phi_N \notin Q \\ \left(1 - \frac{\phi_U}{1-\phi_Q}\right) , & \phi_N \in Q \, . \end{cases} \qquad \text{[S67]}$$

Conversely, in the regime where complex formation limits growth, we take the limit $\frac{K_m}{[m]^{WT}} \to \infty$ in Eq. [S57] and obtain the expression

$$\frac{\lambda}{\lambda^{WT}} = \begin{cases} \left(1 - \frac{\phi_U}{1-\phi_Q}\right) \left[1 - \frac{\phi_U\left(\frac{d_U}{d}-1\right)}{1+\phi_U\left(\frac{d_U}{d}-1\right)}\right] \left(1 - \frac{\phi_U}{1-\phi_Q}\right) \frac{f_{bn}}{f_{bn}^{WT}} , & \phi_N \notin Q \\ \left(1 - \frac{\phi_U}{1-\phi_Q}\right) \left[1 - \frac{\phi_U\left(\frac{d_U}{d}-1\right)}{1+\phi_U\left(\frac{d_U}{d}-1\right)}\right] \frac{f_{bn}}{f_{bn}^{WT}} , & \phi_N \in Q \, . \end{cases} \qquad \text{[S68]}$$

This equation can be compared with Eq. [S65] for the ratio of bound RNA polymerases.

***The dependency of the growth rate ratio $\frac{\lambda}{\lambda^{WT}}$ on the gene copy number $g_U$.*** So far, we have obtained expressions for the growth rate ratio $\frac{\lambda}{\lambda^{WT}}$ in the presence of unnecessary proteins in terms of the protein fraction $\phi_U$. In the experimental study by Kafri et al. (11), the authors also report measurements of the relative growth rate as a function of the gene copy number $g_U$, which they use to modulate unnecessary protein expression. This section shows how to derive such relationship from our model.

We will begin by providing an intuitive derivation that outlines the basic steps for obtaining the correct expression of $\frac{\lambda}{\lambda^{WT}}$ versus $g_U$, although it is not quantitatively accurate. From Eq. [S38], we can write

$$\omega_U = \frac{\beta_0^U g_U}{\sum_{j \neq U} \beta_0^j g_j + \beta_0^U g_U} = \frac{g_U}{g_U + \Pi} \, , \qquad \text{[S69]}$$

where $\Pi = \sum_{j \neq U} \beta_0^j g_j / \beta_0^U$ represents an effective relative (inverse) promoter strength of unnecessary genes $U$. Since all equations in the previous sections are expressed in terms of $\phi_U$, which can be written in terms of $\omega_U$, we use this expression to estimate the dependency of the relative growth rate on the gene copy number $g_U$. All the expressions derived above will then feature an extra factor containing $\Pi$ in place of $\omega_U$.

Although this approximate argument contains many correct ideas, it fails to account for the presence of the constant-size protein class $Q$. Indeed, if $Q$ is under homeostatic control, the activity of its promoters must also increase to compensate for the increased competition for shared resources such as RNA polymerases and ribosomes allocated to the $U$-sector. Fortunately, we can account for this in a simple way. The full relationship between $\omega_U$ and $g_U$, taking into account homeostasis of protein class $Q$ is the following:

$$\omega_U = \omega_U^{max}(\phi_Q) \frac{g_U}{\Omega(\phi_Q) + g_U} \, , \qquad \omega_U^{max}(\phi_Q) = \frac{1 - \phi_Q}{1 - \phi_Q(1 - \frac{d}{d_U})} \, , \qquad \Omega(\phi_Q) = \Pi \frac{1}{1 - \phi_Q(1 - \frac{d}{d_U})} \, . \qquad \text{[S70]}$$

In the remaining of this section we will derive the formulas above.

Note that accounting for the protein class $Q$ does not require any new parameter, apart from $\Pi$ - which now takes a slightly different form:

$$\Pi = \frac{\sum_{j \neq U,Q} \beta_0^j g_j}{\beta_0^U} \, .$$

This factor is, as in the simplified case introduced above, an effective (inverse) promoter strength relative to the promoter activity of all genes belonging to protein classes different from $Q$ and $U$. To compute the relative growth rate as a function of $g_U$, we can use the expression $\frac{\lambda}{\lambda^{WT}}$ vs $\phi_U$ [S67] and [S68] in conjunction with the expression of $\phi_U$ in terms of $\omega_U$ [S61]. We would then plug in $\omega_U(g_U)$ as in Eq. [S70] and obtain the final $\frac{\lambda}{\lambda^{WT}}$ vs $g_U$.

However, in order to obtain Eq. [S70], we first use the link between the protein fraction and the RNA polymerase allocation. Since the protein sector $Q$ is assumed to stay constant across conditions, we write

$$\phi_Q = \frac{\omega_Q/d}{\sum_{i\neq U,Q}\omega_i/d + \omega_Q/d + \omega_U/d_U} = \phi_Q^{WT} = \omega_Q^{WT} \ . \qquad \text{[S71]}$$

In Eq. [S71], the first equality comes from rewriting the relationship between protein composition and RNA polymerase allocation in the presence of different transcript degradation (using Eq. [S59] for the mRNA abundance $m_i$, computing the mRNA fraction $m_i/m$ and considering the protein fraction $\phi_i = m_i/m$). Next, we express the RNA polymerase allocation in terms of promoter strengths $\omega_i = \frac{\beta_0^i g_i}{\sum_j \beta_j g_j}$. By combining these two expressions, we find how the overall promoter strength of protein class $Q$ changes to keep $\phi_Q$ constant.

To simplify the notation, we introduce the variables $s_Q = \beta_0^Q g_Q$, $s_q = \beta_0^U g_U$, and $s = \sum_{i\neq U,Q}\beta_0^i g_i$, which summarize the total promoter strength (as they also depend on the total number of genes). Note that $s$ does not change under the expression of the unneeded protein under our assumptions. Using this notation, we can rewrite Eq. [S71] as $\phi_Q = \frac{s_Q}{s+s_Q+s_U\frac{d}{d_U}} = \frac{s_Q^{WT}}{s+s_Q^{WT}}$. Therefore, $s_Q = s_Q^{WT}\left(1+\frac{s_U}{s}\frac{d}{d_U}\right)$, which quantifies how much the overall promoter strength of protein class $Q$ should change to keep $\phi_Q$ constant. Furthermore, this shows that $\frac{s_Q^{WT}}{s} = \frac{\phi_Q}{1-\phi_Q}$. After plugging these results into $\omega_U = \frac{s_U}{s+s_U+s_Q}$, we obtain $\omega_U = \frac{s_U/s}{1+\frac{\phi_Q}{1-\phi_Q}+s_U/s\left(1+\frac{\phi_Q}{1-\phi_Q}\frac{d}{d_U}\right)}$. Since $s_Q = \beta_0^U g_U$, this expression provides the link between $\omega_U$ and $g_U$. Finally, rearranging this equation yields Eq. [S70].

We can finally describe the steps required to derive Eq. [S65]. In absence of the unneeded protein sector $U$, the fraction of bound RNA polymerases takes the form outlined in Section S8. In presence of the unneeded protein sector $U$, we isolate the terms corresponding to both protein class $Q$ and $U$, as follows

$$f_{bn} = \frac{\sum_{j\neq U,Q}\beta_0^j\, g_j + \beta_0^Q\, g_Q + \beta_0^U\, g_U}{\left(\frac{k_{tx}}{L_g}\right)\ V + \sum_{j\neq Q,U}\beta_0^j\, g_j + \beta_0^Q\, g_Q + \beta_0^U\, g_U} \ . \qquad \text{[S72]}$$

We can use the expressions derived in the previous subsection to rewrite the expression for $f_{bn}$ in terms of $\phi_Q$, $\omega_U$, and $Z_g$. Using the notation $s$, $s_Q$, and $s_U$, we can simplify the expression and obtain $f_{bn} = \frac{1+\frac{s_U}{s}+\frac{s_Q}{s}}{1+\frac{s_U}{s}+\frac{s_Q}{s}+Z_g}$, where $Z_g = \left(\frac{k_{tx}}{L_g}\right)\frac{V}{s}$ (consistently with the previous notation).

In the absence of the unneeded protein ($s_U = 0$), we have $f_{bn}^{WT} = \frac{1+\frac{s_Q^{WT}}{s}}{1+\frac{s_Q^{WT}}{s}+Z_g^{WT}}$. By using $\frac{s_Q^{wt}}{s} = \frac{\phi_Q}{1-\phi_Q}$, we recover $f_{bn}^{WT} = \frac{1}{1+(1-\phi_Q)Z_g^{WT}}$. In the presence of the unneeded protein, we use the link between $s_U/s$ and $\omega_U$ found in the previous subsection. After some algebra, we obtain the expression $\frac{f_{bn}}{f_{bn}^{WT}} = \frac{1+(1-\phi_Q)Z_g}{1+\left(1-\frac{\omega_U}{\omega_U^{max}}\right)(1-\phi_Q)\ Z_g^{WT}}$.

**Growth rate reduction with with unnecessary proteins in the TX-LIM regime with RNA polymerase-limited transcription.** This subsection focuses on obtaining the relative growth rate $\lambda/\lambda^{WT}$ in the TX-LIM regime with RNA polymerase-limited transcription. We start with Eq. [S25] and obtain

$$\frac{\lambda}{\lambda^{WT}} = \frac{[m]}{[m]^{WT}} \ . \qquad \text{[S73]}$$

It is important to note that this equation assumes that the parameters $J^{TL}$ and $[P]$ do not change under overexpression of unneeded proteins. We also note that on the transcription side, nothing changes in the TX-LIM regime compared to the CF-LIM regime. Therefore, equation [S63] for the ratio $\frac{[m]}{[m]^{WT}}$ remains unchanged. This leads to the following expression for the relative growth rate,

$$\frac{\lambda}{\lambda^{\mathrm{WT}}} = \begin{cases} \left[1 - \frac{\phi_U\left(\frac{d_U}{d}-1\right)}{1+\phi_U\left(\frac{d_U}{d}-1\right)}\right]\left(1-\frac{\phi_U}{1-\phi_Q}\right)\frac{f_{bn}}{f_{bn}^{WT}}\ , & \phi_N \notin Q \\ \left[1 - \frac{\phi_U\left(\frac{d_U}{d}-1\right)}{1+\phi_U\left(\frac{d_U}{d}-1\right)}\right]\frac{f_{bn}}{f_{bn}^{WT}}\ , & \phi_N \in Q\ . \end{cases} \qquad \text{[S74]}$$

All the relationship between $\phi_U$ and the gene copy number $g_U$ and between the $f_{bn}$ and the mechanistic parameters also hold as before. By employing these equations and relationships, Fig. S7 and Fig. S6 compare the drop in growth rate due to protein overexpression under the TX-LIM regime and the CF-LIM regime in the case of constant and variable RNA polymerase respectively.

**Changes in the fraction of actively transcribing RNA polymerases and their effect on the changes in the mRNA concentration and growth rate ratio.** So far we have mainly focused for simplicity on the situation where most RNA polymerase is actively transcribing, $f_{bn} \simeq 1$. This section explores in more detail how the fraction of actively transcribing RNA polymerase changes with over-expression of unneeded proteins, and how this change may affect the total mRNA concentration and the growth rate. In particular, the amount of change in the ratio $\frac{f_{bn}}{f_{bn}^{WT}}$ must depend on the original fraction of active RNA polymerase $f_{bn}^{WT}$. This is most evident when $f_{bn}^{WT} = 1$. In this case protein over-expression by adding unneeded genes cannot further change the fraction of active RNA polymerase, hence $\frac{f_{bn}}{f_{bn}^{WT}} = 1$. If $f_{bn}^{WT} < 1$, this is generally no longer the case. Indeed, adding unneeded genes may increase the overall recruitment of RNA polymerase, thereby changing $f_{bn}$. It is intuitive to think that the amount of change will in turn depend on the availability of RNA polymerase before the perturbation, quantified by $1 - f_{bn}^{WT}$. In the following, we show that this is indeed the case in our model and we provide a detailed quantitative expression of this change.

We have previously derived Eq. [S65] for the relative change of the fraction of DNA-bound RNA polymerase,

$$\frac{f_{bn}}{f_{bn}^{WT}} = \frac{1 + (1 - \phi_Q)\, Z_g^{WT}}{1 + \left(1 - \frac{\omega_U}{\omega_U^{max}}\right)(1 - \phi_Q)\, Z_g} \,. \quad \text{[S75]}$$

In order to understand how this ratio depends on the availability of RNA polymeraese in absence of unnecessary protein expression, we assume $Z_g \approx Z_g^{WT}$, i.e. that the intrinsic ability of the genome to recruit RNA polymerase does not change much under the perturbation. Using $f_{bn}^{WT} = \frac{1}{1+(1-\phi_Q)\, Z_g^{WT}}$, we can express $Z_g^{WT}$ and $Z_g$ in terms of $f_{bn}^{WT}$. With some algebra, we obtain that

$$\frac{f_{bn}}{f_{bn}^{WT}} = \frac{1}{1 - \frac{\omega_U}{\omega_U^{max}}\,(1 - f_{bn}^{WT})} \,. \quad \text{[S76]}$$

This expression shows that $f_{bn}$ changes the most relative to the original $f_{bn}^{WT}$ when nearly all RNA polymerase is initially inactive, that is $f_{bn}^{WT} = 0$, and changes the least when all RNA polymerase is already active before applying the perturbation ($f_{bn}^{WT} = 1$).

Next, we can express this quantity in terms of the unneeded protein fraction $\phi_U$, which is easier to measure experimentally. We use the link between $\omega_U$ and $\phi_U$ provided by Eq. [S62] to obtain

$$\frac{f_{bn}}{f_{bn}^{WT}} = \frac{1}{1 - \frac{\phi_U}{1-\phi_Q}\,(1 - f_{bn}^{WT})\, \frac{[1-\phi_Q(1-\frac{d}{d_U})]}{\frac{d}{d_U}+\phi_U(1-\frac{d}{d_U})}} \,. \quad \text{[S77]}$$

This expression is somewhat difficult to understand. In order to capture the main trends and to highlight the primary biological aspects, we initially approximate it for $\phi_U \simeq 0$, which is particularly pertinent for our data analysis. This yields

$$\frac{f_{bn}}{f_{bn}^{WT}} \approx 1 + \phi_U\, \frac{1 - f_{bn}^{WT}}{1 - \phi_Q} \left[\frac{d_U}{d}\,(1 - \phi_Q) + \phi_Q\right] \,. \quad \text{[S78]}$$

As the slope of this linear expansion is always positive, the expression shows that the fraction of active RNA polymerase always increases under sufficiently small protein over-expression, except if $f_{bn}^{WT} = 1$ as all RNA polymerase is already bound. Note also that the slope is a function of the system parameters $f_{bn}^{WT}$, $\phi_Q$ and $\frac{d_U}{d}$.

Next, we show how the perturbation affects the total mRNA concentration, through the relative ratio $\frac{[m]}{[m]_{WT}}$. Since $\frac{[m]}{[m]_{WT}} \propto \frac{f_{bn}}{f_{bn}^{WT}}$, an increase in the fraction of active RNA polymerase always increases the total mRNA concentration. However, this tendency is counterbalanced by the other term in the ratio $\frac{[m]}{[m]_{WT}}$ (see Eq. [S66]), which tends to decrease the total mRNA concentration. Consequently, the largest term determines whether $\frac{[m]}{[m]_{WT}}$ decreases or increases. Here, we investigate the exact conditions for this scenario in the limit of $\phi_U$ close to zero. In the following, we sketch the required steps and provide the general expression. We use Eq. [S64] for the mRNA ratio in conjunction with the expression for $\frac{f_{bn}}{f_{bn}^{WT}}$ found in equations [S77]. Susequently, we perform a first-order Taylor expansion around $\phi_U = 0$. The final result is the following:

$$\frac{[m]}{[m]_{WT}} \approx \begin{cases} 1 + \phi_U \left\{-\left(\frac{d_U}{d} - 1\right) + \frac{1-f_{bn}^{WT}}{1-\phi_Q}\left[\frac{d_U}{d}\,(1-\phi_Q) + \phi_Q\right]\right\} & \text{if } N \in Q \\ 1 + \phi_U \left\{-\left(\frac{d_U}{d} - 1\right) + \frac{1-f_{bn}^{WT}}{1-\phi_Q}\left[\frac{d_U}{d}\,(1-\phi_Q) + \phi_Q\right] - \frac{1}{1-\phi_Q}\right\} & \text{if } N \notin Q \,. \end{cases} \quad \text{[S79]}$$

The slope of this expression can take both positive and negative values, depending on the value of the parameters. This means that total mRNA concentration can both increase and decrease under protein over-expression. The model provides precise conditions under which this switch occurs. We can formulate the following general remarks regarding these conditions:

1. If all RNA polymerase is actively transcribing before the perturbation ($f_{bn}^{WT} = 1$), then the slope is always negative and the total mRNA concentration always decreases.

2. If $f_{bn}^{WT} < 1$ and the RNA polymerase sector is part of the Q sector, the sign of the slope changes from positive to negative for a precise value of $\frac{d_U}{d}$. Thefore, total mRNA concentration may increase at $d_U = d$ but decreases as $d_U$ increases (as unneded mRNA becomes more unstable).

3. If $f_{bn}^{WT} < 1$ and the RNA polymerase sector is not part of the Q sector, then the slope is always negative and total mRNA concentration always decreases. We sketch the proof of this fact. Let us call $s$ the slope. First, we can prove that $\frac{\partial s}{\partial \frac{d_U}{d}} < 0$. Consequently, if we prove that $s(\frac{d_U}{d} = 1)$ is negative, we automatically prove that the slope $s$ is always negative for any $d_U > d$. By plugging in $\frac{d_U}{d} = 1$ in the expression above, we can compute $s = -\frac{f_{bn}^{WT}}{1-\phi_Q}$, which is indeed negative.

Note that for sufficiently high unnecessary protein expression, the increasing trend of total mRNA must break down, as the fraction of active RNA polymerases becomes close to 1 and after this point the mRNA concentration will decrease again; indeed, using the exact expression rather than the first-order expansion shows this behavior.

Finally, we may use the link between the growth rate and the mRNA concentration to study how the changes in the fraction of actively transcribing RNA polymerase affect the growth rate. Conceptually, this is straightforward as one simply needs to substitute all the expressions found in this section into Eq. [S57], re-written here for the sake of readability,

$$\frac{\lambda}{\lambda^{WT}} = \frac{\phi_R}{\phi_R^{WT}} \frac{[m]}{[m]^{WT}} \frac{1 + \frac{K_m}{[m]^{WT}}}{\frac{[m]}{[m]^{WT}} + \frac{K_m}{[m]^{WT}}} \,. \qquad \text{[S80]}$$

Fig. S12 shows the growth rate changes in the particular case where $f_{bn}^{WT} = 0.5$, which lies in the regime with significant change in the fraction of active RNA polymerase under unneeded protein over-expression. We used model parameters that reproduce the data of Kafri and coworkers (17).

### S11. Derivation of growth-optimal proteome composition

This section describes in further detail the derivation of the coarse-grained proteome composition that maximizes the growth rate in our model, and describes how the the optimal-growth assumption leads to the prediction of specific growth laws involving mRNAs and proteins. We begin by writing the three relevant equations for optimization, illustrated by Fig. 3 of the main text,

$$\lambda = \gamma \phi_R \frac{\phi_N}{\epsilon^{-1} + \phi_N} \qquad \text{[S81]}$$

$$\gamma \phi_R \frac{\phi_N}{\epsilon^{-1} + \phi_N} = \nu \phi_C \qquad \text{[S82]}$$

$$\phi_R + \phi_N + \phi_C = 1 - \phi_Q \,. \qquad \text{[S83]}$$

We indicate with a star $^*$ the optimized variables (*e.g.*, $\lambda^*$, $\phi_R^*$, $\phi_N^*$ etc.). We now derive an expression of the optimized variables in terms of the parameters $\gamma$, $\nu$, $\epsilon$ and $\phi_Q$ only. In order to do that, it is convenient to define the two parameters $\bar{\lambda} := \frac{\lambda}{\gamma}$ and $\bar{\nu} := \frac{\nu}{\gamma}$, which are a dimensionless growth rate a dimensionless nutrient quality. Note that $\bar{\lambda}$ is the rescaled growth rate that we plot on the $x$ axis of all the figures related to Fig. 3 in the main text. We also define $\phi_{max} := 1 - \phi_Q$, as it simplifies the notation. $\phi_{max}$ is the maximum size that can be taken by any protein class assuming that the protein sector $Q$ takes a constant fraction $\phi_Q$ of the proteome, regardless of the growth condition or perturbation.

Using Eq. [S82] into Eq. [S83] leads to an expression of the ribosomal protein fraction in terms of the RNA polymerase protein fraction

$$\phi_R = \frac{\phi_{max} - \phi_N}{1 + \frac{1}{\bar{\nu}} \frac{\phi_N}{\epsilon^{-1} + \phi_N}} \,. \qquad \text{[S84]}$$

Substituting this equation into the expression of the growth rate gives

$$\bar{\lambda} = \frac{\bar{\nu}}{1 + \bar{\nu}} \frac{\phi_N \left(\phi_{max} - \phi_N\right)}{\frac{\bar{\nu}}{1+\bar{\nu}}\, \epsilon^{-1} + \phi_N} \,. \qquad \text{[S85]}$$

This expression gives the growth rate as a function of the RNA polymerase protein fraction only, thanks to the two constraints from Eqs. [S82] and [S83]. The function takes the simple form $f(x) \approx \frac{x(a-x)}{b+x}$, which always exhibits a maximum for $x < a$ and $b > 0$. Next, we take the first derivative and set it to zero, to compute the RNA polymerase fraction such that the growth rate is maximum. A straightforward calculation leads to the result found in the main text:

$$\frac{\phi_N^*}{\phi_{max}} = \frac{\epsilon^{-1}}{\phi^{max}} \frac{\bar{\nu}}{1 + \bar{\nu}} \left[ \sqrt{1 + \left( \frac{\epsilon^{-1}}{\phi^{max}} \frac{\bar{\nu}}{1 + \bar{\nu}} \right)^{-1}} - 1 \right] \,. \qquad \text{[S86]}$$

All the other growth-optimized variables are obtained from this expression using the relationships derived above. For completeness, we summarize them here

$$\phi_R^* = \frac{\phi_{max} - \phi_N^*}{1 + \frac{1}{\bar{\nu}} \frac{\phi_N^*}{\epsilon^{-1} + \phi_N}} \,, \qquad \phi_C^* = 1 - \phi_Q - \phi_R^* - \phi_N^* \,, \qquad \bar{\lambda}^* = \frac{\bar{\nu}}{1 + \bar{\nu}} \frac{\phi_N^* \left(\phi_{max} - \phi_N^*\right)}{\frac{\bar{\nu}}{1+\bar{\nu}}\, \epsilon^{-1} + \phi_N^*} \,. \qquad \text{[S87]}$$

Note that all the growth optimized variables (excluding $\phi_{max}$, which by assumption is a constant) are a function of two parameters, $\bar{\nu}$ and $\epsilon$. To obtain their trends across nutrient conditions, we vary the parameter $\bar{\nu}$, as in Fig. 3 of the main text. To obtain trends across transcription-targeting drugs, we vary the parameter $\epsilon$, as in Fig. 4 of the main text.

**Growth laws under growth-rate optimization.** Next, we can visualize the relationship between the growth-optimized variables by plotting them against one another. In particular, we can plot the optimal fraction of a protein class against the optimal growth rate. For instance, we consider $\phi_N^*(\bar{\nu}, \epsilon)$ and $\bar{\lambda}^*(\bar{\nu}, \epsilon)$ while varying $\bar{\nu}$. The resulting relationships are so-called growth laws (18). We note that in the literature people often introduce growth laws without referencing to an optimization principle, essentially as constraints between the growth rate and the catalysts of biosynthesis at balanced growth (1) (as our Eq. [S81]). In this work, we mainly use this term to mean the relationship between the growth rate and proteome composition after optimization, following reference (18).

Finally, we briefly describe how to derive an mRNA growth law (10) within our framework. The equation for total mRNA concentration is $[m] = K_m\, \epsilon\, \phi_N$. At optimality, the resulting mRNA concentration is $[m]^*(\bar{\nu}, \epsilon) = K_m\, \epsilon\, \phi_N^*(\bar{\nu}, \epsilon)$. Consequently, we can obtain an mRNA growth law by plotting $[m]^*(\bar{\nu}, \epsilon)$ against $\bar{\lambda}^*(\bar{\nu}, \epsilon)$ under different assumptions, leading to the plots shown in Fig 3 of the main text.

**Square-root growth laws.** In the main text, we stated that the CF-LIM regime, under the assumption of growth-rate optimization and not further constraint (and specifically, if the supply-demand trade-off parameter $\epsilon$ is constant), predicts square-root growth laws in RNA polymerase and mRNA content, instead of the typical linear law (2, 19). In this section, we show how our model can generate such square-root growth laws in this scenario of constant $\epsilon$.

Let us recall the equation of the growth rate,

$$\lambda = \gamma \phi_R \frac{\phi_N}{\epsilon^{-1} + \phi_N} \; . \qquad \text{[S88]}$$

We start by presenting an intuitive argument in a specific limit. First, we consider the case where the nutrient quality is very high, threfore the catabolic sector $\phi_C$ is negligible. In such a case, the normalization condition is $\phi_R + \phi_N = 1 - \phi_Q$. Second, we consider the limit where $\epsilon \to 0$ and $\lambda \to \gamma\epsilon\, \phi_R \phi_N$. Maximizing this expression under the normalization constraint gives $\phi_R^* = \phi_N^*$. Therefore, $\lambda^* \propto \phi_N^*\, \phi_N^*$, which implies that a square-growth law $\phi_N^* \propto \sqrt{\lambda^*}$ emerges naturally if the growth rate is proportional to the product of two sectors.

To provide a more rigorous derivation, we simplify our notation as follows: we define $\alpha = \frac{\epsilon^{-1}}{\phi^{max}} \frac{\bar{\nu}}{1+\bar{\nu}}$, $\lambda' = \left(\frac{\epsilon^{-1}}{\phi^{max}}\lambda\right)$, and $\phi_N' = \frac{\phi_N}{\phi^{max}}$. Using this notation, we can rewrite the equation from the previous section as follows

$$\phi_N'^* = \alpha \left[\sqrt{1+\alpha^{-1}} - 1\right] \, , \qquad \text{[S89]}$$

$$\lambda'^* = \phi^{max} \alpha \frac{\phi_N'^* \left(1 - \phi_N'^*\right)}{\alpha + \phi_N'^*} \, . \qquad \text{[S90]}$$

The two above equations implictly contain the growth law that relates $\phi_N'^*$ and $\lambda'^*$. To derive this law, we need to re-write $\alpha$ in terms of $\lambda'^*$ and substitute it into the first equation. However, due to the presence of a square root, this task is not simple and requires some approximations. We can consider two special cases:

1. The first case is the limit $\phi_N'^* \ll 1$, and consequently $\alpha\left[\sqrt{1+\alpha^{-1}} - 1\right] \ll 1$, which implies $\alpha \ll 1$. In this limit, $\phi_N'^* \approx \sqrt{\alpha}$. Using this expression into $\lambda'^*$, and since $\left(1 - \phi_N'^*\right) \approx 1$ and $\frac{\sqrt{\alpha}}{\sqrt{\alpha}+\alpha} \approx 1 - \sqrt{\alpha}$, in this limit $\alpha \ll 1$, and we find $\lambda'^* \approx \phi^{max}\alpha\,(1 - \sqrt{\alpha})$. Since $\phi_N'^* \approx \sqrt{\alpha}$ and $\alpha$ is small, this means that approximately $\lambda'^* \propto \phi_N'^{*2}$, which gives a square-root growth law when inverted.

2. The second special case is the limit $\phi_N'^* \approx \frac{1}{2}$ (however not so realistic as $\phi_N \simeq$0.01 in *E. coli*). Within this limit, $\phi_N'^*\left(1 - \phi_N'^*\right) \approx \phi_N'^{*2}$. Using the equations above, $\phi_N'^* \approx \frac{1}{2}$ meaning that $\alpha\left[\sqrt{1+\alpha^{-1}} - 1\right] \approx \frac{1}{2}$, which is possible only when $\alpha^{-1} \approx 0$ or $\alpha \gg 1 > \phi_N'^*$. Therefore, $\lambda'^* \propto \phi_N'^{*2}$; inverting this expression gives once again a square-root growth law.

### S12. Response to transcription-targeting drugs under constant level of RNA polymerase fraction and optimized growth rate

If the RNA polymerase protein fraction is maintained constant across perturbations by a physiological response, the derivation of sec. S11 does not apply. This section addresses protein composition under treatment of trascriptional targeting drugs, in the scenario the RNA polymerase protein fraction is kept constant and that growth is optimized. The relevant equations for optimization remain the same,

$$\lambda = \gamma \phi_R \frac{\phi_N}{\epsilon^{-1} + \phi_N} \qquad \text{[S91]}$$

$$\gamma \phi_R \frac{\phi_N}{\epsilon^{-1} + \phi_N} = \nu \phi_C \qquad \text{[S92]}$$

$$\phi_R + \phi_N + \phi_C = 1 - \phi_Q \, . \qquad \text{[S93]}$$

The RNA polymerase protein fraction $\phi_N$ is now part of the protein fraction $\phi_Q$. Therefore, it is helpful to make the following change of variables $\phi_Q \to \bar{\phi_Q} = \phi_Q + \phi_N$ and $\gamma \to \bar{\gamma} \frac{\phi_N}{\epsilon^{-1}+\phi_N}$. The three equations above become

$$\lambda = \bar{\gamma}\phi_R \qquad \text{[S94]}$$

$$\gamma\phi_R\bar{\gamma} = \nu\phi_C \qquad \text{[S95]}$$

$$\phi_R + \phi_C = 1 - \bar{\phi_Q}\,. \qquad \text{[S96]}$$

These equations are formally identical to the model described in ref. (2). The only difference is that the pseudo-translation-rate $\bar{\gamma}$ can now also be modulated by transcription-targeting drugs as it contains the parameter $\epsilon$. The solutions for these three equations are

$$\lambda = (1 - \bar{\phi_Q})\,\frac{\nu\,\bar{\gamma}}{\bar{\gamma}+\nu}\,; \qquad \phi_R = (1 - \bar{\phi_Q})\,\frac{\nu}{\bar{\gamma}+\nu}\,; \qquad \phi_C = (1 - \bar{\phi_Q})\,\frac{\bar{\gamma}}{\bar{\gamma}+\nu}\,. \qquad \text{[S97]}$$

In particular, when the nutrient quality $\nu$ is kept constant, while $\bar{\gamma}$ changes, we can obtain the growth laws

$$\phi_R = (1 - \bar{\phi_Q}) - \frac{\lambda}{\nu},; \qquad \phi_C = \frac{\lambda}{\nu}\,. \qquad \text{[S98]}$$

Hence, in this scenario, decreasing $\bar{\gamma}$ with transcription-targeting drugs decreases the growth rate $\lambda$.

### S13. The effect of rRNA transcription

The model variants considered so far do not explicitly incorporate rRNA transcription. This section describes how to extend our framework to include rRNA transcription and discusses the prediction of these extended models for budding yeast and *E. coli*.

At the translation level, we retain the same model. The production of a protein $P_i$ is proportional to the number of transcripts $m_i$ times the protein flux per transcript $J^{TL}$, which takes the usual form $J^{TL}([R],[m]) = \frac{k_{tl}}{L_p}\frac{[R]}{K_m+[m]}$, where $R$ and $m$ are the total numbers of ribosomes and mRNAs, respectively (sec. S2 of this SI Appendix). In particular, $\frac{dP_i}{dt}$ follows Eq. [S12)].

Throughout this section, we will still work under the simplifying assumption of perfect stochiometry between ribosomes, rRNA and ribosomal proteins (20–23). This is equivalent to assuming a fast assembly time for the ribosome compared to the production time of its components, with perfect coordination between rRNA and ribosomal proteins, due to regulatory systems that we do not describe. Consequently, the number of ribosomes $R$ is always proportional to the number of ribosomal proteins $P_R$ and the number of rRNAs $r$, with a proportionality constant that depends only on the stochiometry of the ribosome, which in turn is independent of the particular growth conditions. Under this assumption, Eq. [S14] for the growth rate used throughout this study still applies

$$\lambda = \gamma\phi_R\frac{[m]}{K_m+[m]}\,. \qquad \text{[S99]}$$

According to this equation, rRNA transcription may affect the growth rate only through its influence on ribosomal protein fraction $\phi_R$ or on the total mRNA concentration $[m]$. However, under the assumption of perfect coordination between the production of rRNA and ribosomal proteins, the first case is radically simplified by the assumption of a linear relationship between the number of rRNAs and the number of ribosomes. While we note that this assumption may break down under particular perturbations, we deal in the following with the second case, which is more complex even under the assumption of perfect stoichiometry.

We distinguish two ways of extending our present framework with rRNA transcription: (i) rRNA transcription is carried by a specific type of RNA polymerase that only transcribes rRNA and (ii) rRNA transcription is carried by the same RNA polymerase that also transcribes mRNA. The first case obviously applies to eukaryotic transcription, including *S. cerevisiae*. The second case applies to prokaryotes, including *E. coli*. We begin by describing the first case, which is considerably simpler.

**Model extension with rRNA and two classes of RNA polymerases.** Let us call $N_m$ and $N_r$ the polymerases transcribing mRNA and rRNA respectively. These are distinct macro-molecular complexes and there is no common pool that transcribes both mRNA and rRNA. We assume that transcription is limited by RNA polymerase availability and obtain the following equations for the dynamics of mRNA ($m_i$) and rRNA ($r$) amounts (see sec. S1),

$$\frac{dm_i}{dt} = \frac{k_{tx}}{L_{g_m}}\,\omega_i\,f_{bn}\,N_m - d\,m_i \qquad \text{[S100]}$$

$$\frac{dr}{dt} = \frac{k_{tx}}{L_{g_r}}\,f_{bnr}\,N_r\ , \qquad \text{[S101]}$$

where we assumed that all RNAs are elongated with the same rate $k_{tx}$, mRNAs are degraded with the same rate $d$, rRNAs are not degraded as they are stably incorporated inside ribosomes, and mRNAs and rRNAs have length $L_{g_m}$ and $L_{g_r}$, respectively. Finally, we call $f_{bn}$ and $f_{bnr}$ the fraction of polymerases that are bound and active for the two RNA types.

Under the assumption of perfect stoichiometry, we can convert the count of polymerase protein subunits into the count of polymerases for the two categories, $N_m = P_{N_m}/n_m$ and $N_r = P_{N_r}/n_r$, where $n_m$ and $n_r$ represent the number of proteins forming RNA polymerases transcribing mRNAs and rRNAs, respectively. With this assumption, we can define the compound rate parameters $\gamma_{tx}^m := \frac{k_{tx}}{L_{gm} n_m}$ and $\gamma_{tx}^r := \frac{k_{tx}}{L_{gr} n_r}$. Imposing quasi-steady-state on the mRNA and re-writing the rRNA equation, we get

$$m_i = \frac{\gamma_{tx}^m}{d} \, \omega_i \, f_{bn} \, P_{N_m} \ , \qquad \text{[S102]}$$

and

$$\frac{dr}{dt} = \gamma_{tx}^r \, f_{bnr} \, P_{N_r}. \qquad \text{[S103]}$$

Let us now find the total mRNA and rRNA concentration. To find the former, we perform the sum $\sum_i m_i$ and divide by the volume. Crucially, $\sum_i \omega_i = 1$, since the RNA polymerases transcribing mRNAs do not transcribe rRNA. To find the steady-state rRNA concentration, we perform the chain rule on $r/V$, keeping in mind that volume is proportional to total protein content $P$ and total protein content grows exponentially with growth rate $\lambda$, which is given by Eq. [S44] as usual (rewritten above as Eq. [ S99]). This gives

$$[m] = \frac{\gamma_{tx}^m}{d} \, f_{bn} \, [P] \, \phi_{N_m}, \qquad \text{[S104]}$$

$$[r] = \frac{\gamma_{tx}^r}{\lambda} \, f_{bnr} \, [P] \, \phi_{N_r}. \qquad \text{[S105]}$$

These equations immediately show that rRNA production does not affect directly mRNA production. Increasing rRNA transcription may come at the expense of mRNA only through a global constraint on the proteome fraction, which includes $\phi_{N_m}$ and $\phi_{N_r}$, but such potential trade-off is already contained in the model without rRNA, as rRNA and ribosomal proteins are proportional ($[r] \sim \phi_R$). Additionally, coordination between rRNA and ribosomal proteins implies that $[r] \sim [P]\phi_R$ (i.e. ribosomal proteins and rRNAs are present in a fixed ratio in a ribosome). In turn, this fixes the amount of protein resources $\phi_{N_r}$ allocated to rRNA production as

$$\phi_{N_r} \sim \frac{\phi_R \, \lambda}{\gamma_{tx}^r \, f_{bnr}} = \phi_R^2 \frac{\gamma}{\gamma_{tx}^r \, f_{bnr}} \frac{[m]}{K_m + [m]} \ . \qquad \text{[S106]}$$

A version of this equation was found before for the case $K_m = 0$ (pure translation-limitation) (20, 21, 23). In this case, the term with the mRNA concentration disappears and $\phi_{N_r} \sim \phi_R^2 \sim \lambda^2$, which is a non-linear growth law (although other terms in Eq. [S106] may depend on $\lambda$ and affect the quadratic scaling). In general, in this extension of the model, rRNA is merely an extra-variable that is enslaved to all the variables present in the original model. In particular, this means that all our analysis related to *S. cerevisiae* should be largely independent of explicitly considering rRNA production in our model.

**Model extension with rRNA and one RNA polymerase.** In this model variant, there is a common pool that transcribes both mRNA and rRNA. Following sec. S1, we get

$$\frac{dm_i}{dt} = \frac{k_{tx}}{L_{gm}} \, \omega_i \, f_{bn} \, N - d \, m_i \ , \qquad \text{[S107]}$$

and

$$\frac{dr}{dt} = \frac{k_{tx}}{L_{gr}} \, \omega_r \, f_{bn} \, N \qquad \text{[S108]}$$

for the mRNA and rRNA dynamics resepctively, where $\omega_i$ is the fraction of RNA polymerases transcribing type-$i$ mRNA, while $\omega_r$ is the fraction of RNA polymerases transcribing rRNA, and we assumed once again that all mRNAs are degraded with the same rate $d$ and elongated with the same rate $k_{tx}$, and rRNAs are not degraded. Imposing quasi-steady-state for mRNAs as above, we get

$$m_i = \frac{\gamma_{tx}^m}{d} \, f_{bn} \, \omega_i \, P_N, \qquad \text{[S109]}$$

$$\frac{dr}{dt} = \gamma_{tx}^r \, f_{bn} \, \omega_r \, P_N, \qquad \text{[S110]}$$

leading to the following equations for total mRNA and rRNA concentrations,

$$[m] = \frac{\gamma_{tx}^m}{d} \, f_{bn} \, [P] \, \omega_m \, \phi_N, \qquad \text{[S111]}$$

$$[r] = \frac{\gamma_{tx}^r}{\lambda} \, f_{bn} \, [P] \, \omega_r \, \phi_N \ , \qquad \text{[S112]}$$

where $\omega_m = (1 - \omega_r)$ reflects allocation of RNA polymerase to mRNA transcription.

We note that in this model variant, total mRNA and rRNA concentration are directly related since allocating RNA polymerases to one of the RNA species immediately decreases the RNA polymerase availability to the other species. The expression for $\omega_r$ in terms of the other model derives from the assumption of perfect coordination between rRNA and ribosomal proteins, that is $[r] \sim [P]\phi_R$. Hence, considering Eq. [S99],

$$\omega_r \ \sim \frac{\phi_R \, \lambda}{\gamma_{tx}^r \, f_{bn} \, \phi_N} = \phi_R^2 \, \frac{\gamma}{\gamma_{tx}^r \, f_{bn} \, \phi_N} \frac{[m]}{K_m + [m]} \ . \qquad \text{[S113]}$$

In addition, in this scenario proteome composition $\phi_i$ $(P_i/P)$, ribosome allocation $\chi_i$ $(m_i/m)$ and RNA polymerase allocation $\omega_i$ follow a different relation than the one found in sec. S7 of this SI Appendix,

$$\phi_i = \chi_i = \frac{\omega_i}{\omega_m} \qquad \forall\ i\ . \tag{S114}$$

**Change in RNA polymerase allocation to rDNA as a result of changes in ribosome proteome fraction $\phi_R$.** Eqs. [S113] and [S114] have several interesting consequences. First, increasing the ribosome fraction $\phi_R$ while keeping fixed all other quantities, including total RNA polymerase concentration leads to a decrease in the mRNA concentration $[m]$ (due to coordination between rRNA and r-proteins, greater $\phi_R$ also implies more RNA polymerases devoted to rRNA over mRNA). Note that the total mRNA concentration in order to increase with increasing growth, as observed in *E. coli* has to contrast the increase of $\phi_R$, diverting RNA polymerases to rRNA.

We proceed to quantify how much $[m]$ should change if one hypothetically changes $\phi_R$. From Eqs.[S111], [S112] and Eq. [S99] we can write the following equation for the relative change of $[m]$ as the ribosomal fraction changes $\phi_R \to \phi_R^{'}$

$$\frac{[m](\phi_R^{'})}{[m](\phi_R)} = \frac{1-\omega_r(\phi_R^{'})}{1-\omega_r(\phi_R)}\ . \tag{S115}$$

Let us first discuss the nucleotide and protein composition of *E. coli* ribosomes (24). Ribosomes are made of large and a small sub-unit, respectively called the 50S and the 30S unit. All sub-units are made of both proteins and rRNA. Focusing on the RNA components, 5S and 23S rRNA are the RNA components of the 50S subunit and 16S rRNA is the RNA component of the 30S subunit. In *E. coli*, 5S, 23S and 30S are respectively 120, 2906 and 1542 nucleotides long. Consequently, a ribosome contains 4568 rRNA nucleotides in total. We will count these as “one” rRNA. Considering the protein content, there are 21 proteins in 30S and 31 proteins in 50S. So for each rRNA (counted as a 4568-nucleotide block), we have 52 proteins, assuming perfect stochiometry. This implies $r = P_R/52$, and in terms of concentrations $[r] = [P]\ \phi_R/52$

By using Eq. [S112], we obtain the following expression for $\omega_r$,

$$\omega_r = \frac{1}{52}\frac{\phi_R\ \lambda}{\gamma_{tx}^r\ f_{bn}\ \phi_N} = \frac{\phi_R^2}{52}\ \frac{\gamma}{\gamma_{tx}^r\ f_{bn}\ \phi_N}\frac{[m]}{K_M+[m]}\ . \tag{S116}$$

Note the $\phi_R^2$ dependency, due to the fact that both growth rate and rRNA concentration are proportional to $\phi_R$ (21, 25).

If the transcription elongation rate $k_{tx}$ is 40 nt per second and the number of nucleotides in the rRNA in a ribosome is 4568 nt, then

$$\gamma_{tx}^r = k_{tx}/(L_{g_r} n_r) = 0.0088 \cdot \frac{1}{5}\ s^{-1}\ , \tag{S117}$$

which gives, $\gamma_{tx}^r \simeq 6.3\ h^{-1}$. Further, we can use $\gamma = 10\ h^{-1}$ for fast growth (26) and $\phi_N \approx 0.01$ (10) and $f_{bn} \approx 0.5$ (16). We can say that $\frac{[m]}{K_M+[m]} \approx 0.1$ in a CF-LIM regime and 1 in a TL-LIM regime. Using Eq. [S116], we get $\omega_r \approx 6.1\ \phi_R^2$ for TL-LIM and $\omega_r \approx 0.5\ \phi_R^2$ for CF-LIM.

In these two situations, we consider the change in mRNA concentration that follows a change in ribosomal protein fraction from 0.1 to 0.2 keeping everything else constant using Eq. [S115]. In the TL-LIM regime ($\frac{[m]}{K_M+[m]} \approx 1$), we get $\frac{[m](\phi_R=0.2)}{[m](\phi_R=0.1)} \approx 0.8$, whereas in a the CF-LIM regime with $\frac{[m]}{K_M+[m]} \approx 0.1$), we get $\frac{[m](\phi_R=0.2)}{[m](\phi_R=0.1)} \approx 0.98$. Both changes are tiny compared to the at least 300% change observed in the mRNA concentration across conditions with the same ribosomal protein fractions (10).

To conclude this part we observe that the new term in Eq. [S111] affects growth rate optimization (Fig. 3 of the main text). To solve the optimization problem, one needs to consider the growth rate expression $\lambda = \gamma\phi_R \frac{[m]}{K_m+[m]}$, use Eq. [S111] for $[m]$ and find the optimal protein composition $(\phi_R, \phi_N)$ that maximizes the growth rate.

**Effect of rRNA transcription on the cost of unneeded proteins.** Additionally, this model variant predicts that the effect of over-expression of proteins on growth rate should be complicated by the fact that it biases transcription of mRNA with respect to rRNA, which increases total mRNA concentration. Referring to cells without unneeded proteins as $WT$ (as above), under perfect ribosome stochiometry the relative change in growth rate is still determined by equation [S57]. Therefore, the ratios $\frac{\phi_R}{\phi_R^{WT}}$ and $\frac{[m]}{[m]^{WT}}$ determine the relative growth rate. The ribosomal fraction ratio does not change when taking into account rRNA transcription, if we still assume (Eq. [S58]) that each protein fraction is affected in the same way by the presence of the unneeded protein sector. Conversely, the mRNA ratio changes significantly compared to the previous model. If we perform the calculation in the case without any modulation of the unneeded mRNA stability (that is, $d_U = d$ in our notation), we obtain

$$\frac{[m]}{[m]^{WT}} = \left[1 + \frac{\phi_U}{(1-\phi_Q)-\phi_U}\ \omega_r^{WT}\right]\ \frac{f_{bn}}{f_{bn}^{WT}}\ \frac{\phi_N}{\phi_N^{WT}} \tag{S118}$$

In this expression, the term $\left[1 + \frac{\phi_U}{(1-\phi_Q)-\phi_U}\ \omega_r^{WT}\right]$ increases with unneeded protein expression, a phenomenon not necessarily leading to a total mRNA increase, as it depends also on $\frac{f_{bn}}{f_{bn}^{WT}}$ and $\frac{\phi_N}{\phi_N^{WT}}$. However, this mechanism, absent in the previous model without rRNA transcription, indicates that total mRNA is more likely to increase in the presence of unneeded proteins due to the reallocation of RNA polymerases from rRNAs to mRNAs.

In summary, while a full exploration of this model extension is beyond the scope of this work, our preliminary analysis indicates the possibility of interesting trade-offs determined by RNA polymerase sequestration from rDNA.

**Derivation of Eqs. [S118].** This subsection details the derivation of Eqs. [S118]. Summation of mRNA amounts and volume normalization gives the following expression for the concentration

$$[m] = \left( \omega_U + \sum_{i \neq U,r} \omega_i \right) \gamma_{tx}^m \, f_{bn} \, \phi_N \ . \tag{S119}$$

By analyzing the ratio $\frac{[m]}{[m]^{WT}}$, and assuming that elongation and degradation rates remain unchanged upon the perturbation, we get

$$\frac{[m]}{[m]^{WT}} = \frac{\omega_m}{\omega_m^{WT}} \frac{f_{bn}}{f_{bn}^{WT}} \frac{\phi_N}{\phi_N^{WT}} \ , \tag{S120}$$

where we have used the normalization condition $\omega_U + \sum_{i \neq U,r} \omega_i + \omega_r = 1$ and $\sum_{i \neq U,r} \omega_i^{WT} + \omega_r^{WT} = 1$. This ratio decreases with increasing $\omega_r$, indicating that the presence of unneeded genes affects RNA polymerase allocation.

To get Eq. [S118] we need to express the RNA polymerase allocation parameters in terms of protein fractions. By using the normalization condition linking $\omega_i$ to $\omega_U$ and $\phi_U$, assuming that the unneeded proteins subtract RNA polymerase from genes and rDNA equally and that the Q sector remains constant, we obtain the following expression in terms of $\phi_U$

$$\frac{[m]}{[m]^{WT}} = \left( 1 + \frac{\omega_U}{1 - \phi_Q} \frac{\omega_r^{WT}}{1 - \omega_r^{WT}} \right) \frac{f_{bn}}{f_{bn}^{WT}} \frac{\phi_N}{\phi_N^{WT}} \ . \tag{S121}$$

Expressing $\omega_U$ in terms of $\phi_U$ gives Eq. [S118].

### S14. Data Analysis: Estimating the proteome fraction of unneeded proteins from Kafri *et al.* 2016

In their study, Kafri and coworkers (17) generate strains with unneeded proteins by integrating $g_U$ gene copies of mCherry/GFP constructs into the genome of *S. cerevisiae*. As a proxy of the levels of gene expression, they measure the total fluorescence level per cell, which we label as $F_U$. Our model quantifies the growth cost in terms of both $g_U$ and the unnecessary protein fraction $\phi_U$, *i.e.*, the number of unneeded proteins divided by the total number of proteins, which is the sole relevant quantity in the regime of translation limitation (2). This section describes how we have used the fluorescence level $F_U$ from their data to infer the protein fraction $\phi_U$.

If the number of unneeded proteins per cell is $P_U$, the total fluorescence is $F_U = f_U P_U$, where $f_U$ is the amount of fluorescence per protein. Let $P$ be the total number of proteins per cell. Then, the protein fraction is $\phi_U = \frac{P_U}{P}$. Finally, let us remember that $[P]$ is the total protein concentration, i.e., $[P] = \frac{P}{V}$. These considerations lead to the following equation linking fluorescence $F_U$ to the protein fraction $\phi_U$,

$$\phi_U = \frac{1}{f_U[P]} \frac{F_U}{V} \ . \tag{S122}$$

Kafri and coworkers measure both $F_U$ and $V$. The factor $\frac{f_U}{[P]}$ is the calibration parameter $c$ that links the fluorescence concentration to the protein fraction. In order to estimate this factor, we make the following assumptions:

1. we assume that $f_U$ does not change across conditions, *i.e.*, that the fluorescence is calibrated so that fluorescence per protein does not vary;

2. we assume that $[P]$ does not change across conditions; that is, the overall protein concentration does not change with the gene copy number $g_U$ or with the stability of the unneeded transcript $d_U$. This is a stronger assumption than the previous one but we note that (i) protein concentration is indeed known to change little across a broad spectrum of conditions for different organisms (27, 28) and that (ii) the coupling between protein content and volume may be based on a robust mechanisms (29–31) that is independent of specific conditions.

Next, according to Kafri and coworkers a single gene construct contributes to $\approx 2\%$ of the proteome (see Fig. 1E and 1H in their study). Note that this is an estimate for the strain without destabilized mRNA. Let us indicate with $\phi_U(1)$, $F_U(1)$ and $V(1)$ the value of the proteome fraction, fluorescence level and cell volume with a single unneeded gene integrated into the genome, for the strain without destabilized mRNA in standard conditions (indicated with SC in the study of Kafri et al.). Using this condition, we estimate

$$c = \frac{1}{f_U[P]} = \frac{V(1)}{F_U(1)} \phi_U \approx \frac{V(1)}{F_U(1)} 0.02 \, . \tag{S123}$$

Estimating $V(1)$ is straightforward because the authors provide a plot of the relationship between volume $V$ and copy number $g_U$ (see Fig. 7 in their study and Fig. S3B here). On the contrary, the authors do not provide a direct plot of the relationship between absolute fluorescence and gene copy number. However, they show a plot of the volume $V$ against the fluorescence $F_U$, as reported in Fig. S3C here. Consequently, we extract the curve $F_U$ vs $g_U$ by putting together data of $V$ *vs* $g_U$ and of $V$ *vs* $F_U$. It is not possible to do this directly because these two plots come from different experiments, therefore there is no one-to-one correspondence between $g_U$ and $F_U$. Instead, we associate a fluorescence $F_U$ to the gene copy number $g_U$ as follows: (i) for each $g_U$ in the plot of $V$ *vs* $g_U$, we extract the corresponding volume $V$, (ii) we find the closest volume to this value from the plot of $V$ *vs* $F_U$, and finally (iii) we extract the corresponding $F_U$. Therefore, we are able to link the gene copy number $g_U$ to the fluorescence $F_U$. Fig. S3D shows the results of this procedure.

The values of $V(1)$ and $F(1)$ were estimated from these plots (see again Fig. S3B and S3D). To account for noisy measurements, we estimate $V(1)$ and $F(1)$ by taking the average of all the points corresponding to $0.5 < g_U < 1.5$. With this procedure, we finally estimate $c \approx 0.62\ \mu\text{m}^3 \times 10^4$A.U. by using the value of $V(1)$ and $F_U(1)$ provided in the study. We have used this value throughout our study to obtain the protein fraction from the fluorescence concentration.

Finally, we convert the plot of the relative growth rate against the fluorescence provided in ref. (17) (also shown in panel S3F) into a plot of the relative growth rate against $\phi_U$. To obtain $\phi_U$, we need to know the volume at a given fluorescence, *i.e.*, the plot $V$ vs $F_U$. Unfortunately, this is not directly possible since the authors perform different experiments for the different curves. To solve this issue, we infer a curve $V(F_U)$ by fitting phenomenologically the plot $V$ vs $F_U$ for the strains without destabilized mRNA. Fig. S3G shows a linear fit that allows us to obtain a functional relationship $V(F_U)$, which can be used to extract a volume from any fluorescence $F_U$. We therefore estimate $\phi_U$ as $c\frac{F_U}{V(F_U)}$ as shown in Fig. S3H. To make sure that our procedure is robust, we also used fits with functional relationship other than linear. Fig. S3M and S3N shows the results using a generalized logistic function and a Fermi function respectively. The resulting curves are very similar, showing that our procedure does not depend on the assumption of a particular functional relationship $V(F_U)$.

Finally, we also note that for the so-called DAmP strain (with destabilized mRNA), no volume measurement are provided, but only fluorescence levels. Throughout this work, we assume that the functional relationship $V(F_U)$ applies also for the strain with destabilized mRNA. While future experiments should test this assumption, we believe that this is reasonable if it is mainly the protein change that causes the volume change. Therefore, we are also able to predict the volume of the strain with destabilized mRNA by using only its measured fluorescence levels and use the volume to convert the fluorescence to protein fraction.

### S15. Data Analysis: Estimating the change in transcript degradation rate from Kafri *et al.* 2016

Kafri and coworkers (17) are able to modulate the mRNA stability of their unneeded mCherry gene construct through a method called DAmP (decreased abundance by mRNA perturbation). DAmP constructs have the same useless gene under the same promoter, but they also integrate an antibiotic resistance cassette after the gene sequence. Therefore, the genes lacks a terminator in the mCherry gene, which reduces mRNA stability. To quantify the reduction of mRNA stability, the authors do not directly measure the degradation rate of the transcript in the two constructs. They provide an indirect measurement through lower level of protein and mRNA expression. In particular they state that "the DAmP strain produced ∼10-fold lower mCherry fluorescence (Fig. S4A) and ∼30-fold less mRNA as measured by qPCR (not shown)", which they take as evidence of reduced stability.

For our model, it is important to know the exact ratio of the two transcript degradation rates to make predictions. To this purpose, the estimates above can be used as a guide, but they are not sufficient. Indeed, they are only order-of-magnitude estimates and they are only indirectly linked to transcript degradation rates. In the following, we outline the procedure we used to estimate the WT-strain-to-DAmP-strain ratio of the transcript degradation rate based on our model and their data.

**Relevant model equations for estimating the ratio of the transcript degradation rate.** Our model provides a general expression of the unneeded protein fraction ($\phi_U$) in terms of the fraction of RNA polymerase transcribing the unneeded genes ($\omega_U$) and the transcript degradation rate ($d_U$) given in Eq. [S61]. In this equation, $d$ is the degradation of all of the other transcripts (consequently, an average degradation rate). Importantly, $\omega_U$ should not depend on the value of the transcript degradation rate $d_U$. This is because the binding affinity of RNA polymerases to the unneeded genes should be independent of the mRNA stability. The RNA polymerase allocation $\omega_U$ depends nearly exclusively on the gene copy number $g_U$ and the promoter strength associated to such genes (see previous sections).

Let us first assume that the degradation rate the degradation rate of the unneeded gene is equal to the average transcript degradation, $d_U = d$. In that case, following Eq. [S61] we get $\omega_U = \phi_U$. Instead, the DAmP strain has a $d_U > d$, implying that its proteome allocation $\phi_U^{DAmP} \neq \phi_U$. After using Eq. [S61] to compute $\phi_U^{DAmP}$, we obtain

$$\frac{\phi_U^{DAmP}}{\phi_U} = \frac{d}{d_U}\frac{1}{1-\omega_U\left(1-\frac{d}{d_U}\right)}\,, \qquad \text{[S124]}$$

where we used the fact that $\omega_U^{DAmP} = \omega_U$ as it is independent of transcript degradation by hypothesis.

To gain some intuition, we provide a rough estimate of $\frac{d}{d_U}$. The ratio $\frac{\phi_U^{DAmP}}{\phi_U^{WT}}$ is roughly 0.1 according to the authors in (17). $\omega_U$ is difficult to measure directly, but it seems reasonable to assume that it should be $\omega_U \approx 0.1 - 0.2$ at most, as RNA polymerases still have to transcribe all the other transcripts as well. Consequently, since $\frac{d}{d_U} < 1$, the factor $\frac{1}{1-\omega_U\left(1-\frac{d}{d_U}\right)}$ is close to 1 up to 10-20%.

Therefore, we can estimate

$$\frac{\phi_U^{DAmP}}{\phi_U^{WT}} \approx \frac{d}{d_U}\,. \qquad \text{[S125]}$$

This estimate tells us that the transcript degradation rate ratio can be approximated by the ratio of the protein fraction. Thus, we can use this input to fix the ratio $d/d_U$ and predict $\lambda/\lambda^{WT}$, as done in Fig. 2 of the main text.

Next, we define a more formal and data-driven procedure to estimate the transcription degradation rate ratio.

**Systematic procedure to estimate the ratio of the transcript degradation rate in the DAmP strain.** To establish a more formal procedure to derive this parameter, we first note that Eq. [S125] above would be true for any gene copy number $g_U$, or rather, for any $g_U$ such that $\omega_U$ is sufficiently small. These considerations also imply that for this range of gene copy numbers, the ratio $\frac{\phi_U^{DAmP}}{\phi_U^{WT}}$ is constant with $g_U$. Additionally, according to Eq. [S124], the ratio $\frac{\phi_U^{DAmP}}{\phi_U^{WT}}$ can be constant with the gene copy number only if it does not depend on $\omega_U$. While technically never true, this condition is approximately true only if $\omega_U$ is small. Therefore, if in the data $\frac{\phi_U^{DAmP}}{\phi_U^{WT}}$ is constant with $g_U$, the system can only be in the regime of the model where Eq. [S125] holds. Consequently, the constant value taken by $\frac{\phi_U^{DAmP}}{\phi_U^{WT}}$ is the ratio of the transcript degradation rates.

Fig. S8 shows how the ratio $\frac{\phi_U^{DAmP}}{\phi_U^{WT}}$ is indeed roughly constant with the gene copy number. Mathematically, this trend is a consequence of the empirical linear dependency of the relative growth rate with both the gene copy number and the protein fraction. To prove this, let us write explicitly the linear trends of the relative growth rate,

$$\frac{\lambda}{\lambda^{WT}} = 1 - \phi_U \, s_\phi \qquad \text{[S126]}$$

and

$$\frac{\lambda}{\lambda^{WT}} = 1 - g_U \, s_g \ , \qquad \text{[S127]}$$

where $s_\phi$ and $s_g$ are the slope of the two trends. By equating the right-hand side of the two equations, it is immediate to obtain $\phi_U = \frac{s_g}{s_\phi} g_U$. Therefore, the ratio of the protein fraction in two different conditions would depend only on the slopes, as long as the trend remains linear, not on the specific value of $g_U$. Fig. S8B shows the relationship between the protein fraction and the gene copy number obtained from the relationship $\phi_U = \frac{s_g}{s_\phi} g_U$, which we use to obtain,

$$\frac{\phi_U^{DAmP}}{\phi_U^{WT}} = \frac{s_g^{DAmP}}{s_\phi^{DAmP}} \frac{s_\phi^{WT}}{s_g^{WT}} \approx \frac{d}{d_U} \ . \qquad \text{[S128]}$$

Fig. S8C reports a plot of this ratio against $g_U$. The value of the ratio corresponds to our estimate of the transcript degradation rate ratio $\frac{d}{d_U}$. Numerically, we find $\frac{d}{d_U} \approx 0.08$, confirming the previous conclusions simple considerations. To conclude, two independent procedures lead to essentially the same estimate of the transcript degradation rates of the DAmP strain from experimental data.

### S16. Data Analysis: Parameters for modelling the growth response to overexpression of unneeded proteins in *S. cerevisiae*

This section describes the procedures to fix the model parameters in order to produce genuine model predictions for the measurements of Kafri and coworkers. Our model of growth response to overexpression of unneded proteins depends on three parameters: (i) the inverse effective promoter strength $\Pi$ of the unneeded construct, (ii) the fraction $\phi_Q$ of the housekeeping protein sector $Q$, which is kept constant across perturbations and (iii) the global promoter activity $Z_g$, which is related to the fraction of bound RNA polymerases. Throughout this work, we make the assumption that $Z_g \approx 0$. This is equivalent to stating that almost all RNA polymerases are bound to genes and transcribing. There is evidence that this is the case for *Escherichia coli* (13, 14), but our main conclusions do not dependent on this choice, and could be generalised.

We hence need to obtain the parameters $\Pi$ and $\phi_Q$ only. For all model variants in any regime, we always extracted these parameters using the two curves of the relative growth rate against the gene copy number and the protein function in ref. (17). Moreover, we always used the two curves corresponding to the stable construct to fit the parameters, generating genuine predictions (without any adjustable parameters) for the unstable mRNA strain (the DAmP strain). We fixed the parameters in different ways depending on the specific model variant and regimes, as described below.

**A. Parameter estimate in the translation limitation regime.** In the translation limiting, TL-LIM, regime, we have

$$\frac{\lambda}{\lambda^{WT}} = 1 - \frac{\phi_U}{1 - \phi_Q} \ . \qquad \text{[S129]}$$

Therefore, we used a linear fit of the plot $\frac{\lambda}{\lambda^{WT}}$ *vs* $\phi_U$. The slope of this plot is equal to $1/(1 - \phi_Q)$, fixing our estimate of $\phi_Q$.

Next, we extract the parameter $\Pi$ from the theoretical curve $\frac{\lambda}{\lambda^{WT}}(g_U)$ having already fixed $\phi_Q$. The exact expression for $\frac{\lambda}{\lambda^{WT}}(g_U)$ is lengthy, but it can be obtained from the previous sections. We performed a fit by minimizing the sum of relative residuals between $\frac{\lambda}{\lambda^{WT}}^{\text{data}}(g_U)$ and $\frac{\lambda}{\lambda^{WT}}^{\text{model}}(g_U)$ (see Fig. S2).

**B. Parameter estimate in the CF-LIM regime.** In the complex-formation limiting, CF-LIM, regime, the predictions depend on the specific value of the parameter $K_m$. In the limit $K_m \to 0$, complex formation matters less and less for determining the growth rate, and cell growth increasingly approximates the translation limiting regime. In the limit $K_m \to \infty$, complex formation strongly affects the growth rate. We have considered these two limiting cases. In the following, we always take the limit $K_m \to \infty$ when we speak of complex-formation limiting regime, where the relative growth rate takes the form given by Eq. [S68]. This assumption may be relaxed without much effort, but $K_m$ would need to be specified.

In order to fix the parameters in the CF-LIM regime, it is important to distinguish between the case of constant and variable RNA polymerase fraction. If the RNA polymerase fraction is constant, the fitting procedure is identical to the TL-LIM regime as the equations do not change as long as the unneded transcript degradation rate is equal to the average transcript degradation rate, i.e., $\frac{d}{d_U} = 1$. Since we fix our parameters from this reference condition, we obtain the same values as for the TL-LIM regime (note however that the model has different predictions when $d_U \neq d$).

If the RNA polymerase fraction is variable, the fitting procedure changes slightly. The following equation now holds

$$\frac{\lambda}{\lambda^{WT}} = \left(1 - \frac{\phi_U}{1-\phi_Q}\right)\left(1 - \frac{\phi_U}{1-\phi_Q}\right), \qquad \text{[S130]}$$

due to the fact that both ribosomes and RNA polymerases are changing. We fit this expression to the empirical plot $\frac{\lambda}{\lambda^{WT}}$ vs $\phi_U$ to obtain the best estimate for $\phi_Q$. Next, we extract $\Pi$ from the theoretical curve $\frac{\lambda}{\lambda^{WT}}(g_U)$ having already fixed $\phi_Q$. Once again, we perform a fit by minimizing the sum of relative residuals between $\frac{\lambda}{\lambda^{WT}}(g_U)^{\text{data}}$ and $\frac{\lambda}{\lambda^{WT}}(g_U)^{\text{model}}$ (see Fig. S2).

Following the procedure above we obtain both model parameters, which we then use to make predictions for the DAmP strain.

### S17. Data Analysis: Parameters for modelling the mRNA growth laws from Balakrishnan *et al.* 2022

This section describes the procedure to fix the parameters in order to formulate predictions for the data of Balakrishnan and coworkers (10). As outlined in the main text and in Section S11 of this SI Appendix, our model provides the following expressions in terms of RNA polymerase-protein fraction and mRNA, under the assumption of growth-optimized conditions

$$\frac{\phi_N^*}{\phi_{max}} = \frac{\epsilon^{-1}}{\phi^{max}}\frac{\bar{\nu}}{1+\bar{\nu}}\left[\sqrt{1+\left(\frac{\epsilon^{-1}}{\phi^{max}}\frac{\bar{\nu}}{1+\bar{\nu}}\right)^{-1}} - 1\right], \qquad \phi_{max} := 1 - \phi_Q\,, \qquad \text{[S131]}$$

and

$$[m]^* = K_m\,\epsilon\,\phi_N^*\,. \qquad \text{[S132]}$$

As noted before, these expressions are independent of whether the supply-demand trade-off parameter $\epsilon$ depends on the effective nutrient quality $\bar{\nu}$, therefore they can be used in different scenarios for this quantity. We explored the two cases where $\epsilon$ is a simple constant and when $\epsilon$ is a function of the nutrient conditions, motivated by the findings of ref. (10).

In addition, our model also provides an expression of the optimized growth rate in terms of $\phi_N^*$ only, which we rewrite here for completeness,

$$\frac{\lambda^*}{\gamma} = \phi_{max}\frac{\bar{\nu}}{1+\bar{\nu}}\frac{\frac{\phi_N^*}{\phi_{max}}\left(1-\frac{\phi_N}{\phi_{max}}\right)}{\frac{\bar{\nu}}{1+\bar{\nu}}\frac{\epsilon^{-1}}{\phi_{max}} + \frac{\phi_N^*}{\phi_{max}}} \qquad \text{[S133]}$$

Using these expressions we obtain the curves (i) $\phi_N^*$ vs $\frac{\lambda^*}{\gamma}$ and (ii) $[m]^*$ vs $\frac{\lambda^*}{\gamma}$. Balakrishan *et al.* (10) provide in their data these curves across different nutrient conditions without rescaling the growth rate by $\gamma$ (the inverse of the time needed to translate all the ribosomal protein making a ribosomes, see section S2). We rescale the growth rate, taking into account the further complication that the parameter $\gamma$ is known to depend on the nutrient conditions for *Escherichia coli*. We use data from Dai *et al.* (26) for $\gamma$ as a function of the growth rate using, and obtain $\frac{\lambda}{\gamma}$ for each nutrient condition.

Subsequently, our parameter-estimation procedure consists of identifying the parameters that fit best all the provided empirical curves. To outline the details of the fitting procedure, we need to consider the possible difference in the behavior of the supply-demand mRNA trade-off $\epsilon$. Next, we consider the two cases discussed in the main text and describe our fitting procedure for each of them.

**Parameter estimation for the CF-LIM model with constant mRNA supply-demand trade-off parameter $\epsilon$.** When we consider that $\epsilon$ is a constant, independent of growth, we use the following steps to estimate it from data, and fit $K_m$:

1. We extract $\phi_Q$ from ref. (2) and set $\phi_Q = 0.55$;
2. We set a series of nutrient qualities $\bar{\nu}$ to obtain the observed range of growth rates using $\frac{\lambda^*(\bar{\nu})}{\gamma}$ from Section S11;
3. We fit the parameter $\epsilon$ by minimizing the sum of squared residuals between the predicted curve $\phi_N^*$ vs $\lambda^*$ and the observed curve in the data. We obtain $\epsilon \simeq 2000$;
4. Given the parameter $\epsilon$ obtained in the previous step, we fit the parameter $K_m$ by minimizing the sum of squared residuals between the predicted curve $[m]^*$ vs $\lambda^*$ and the observed curve in the data. We obtain $K_m = 0.16\ 10^3/\mu m^3$.

**Parameter estimation for the CF-LIM model with growth-rate dependent mRNA supply-demand trade-off $\epsilon$.** The scenario $\epsilon = \epsilon_{max}\frac{n\bar{u}}{1+\bar{\nu}}$ is motivated by data in ref. (10), who find increased sequestration of RNA polymerase by Rsd in conditions with decreasing nutrient quality. This condition, translated into our model, generates a growth-dependent supply-demand mRNA trade-off $\epsilon$ (see Fig. S13). To fix the parameters of our model, in this case we follow the following steps:

1. We extract $\phi_Q$ from ref. (2) and set $\phi_Q = 0.55$;
2. We set a series of nutrient qualities $\bar{\nu}$ to obtain the observed range of growth rates using $\frac{\lambda^*(\bar{\nu})}{\gamma}$ from Section S11;
3. Because of the particular form of $\epsilon$, the expression for the proteome fraction of RNA polymerase proteins simplifies dramatically, and in particular it is independent of nutrient conditions, $\frac{\phi_N^*}{\phi_{max}} = \frac{\epsilon_{max}^{-1}}{\phi^{max}}\left[\sqrt{1+\left(\frac{\epsilon_{max}^{-1}}{\phi^{max}}\right)^{-1}} - 1\right]$. Consequently, RNA polymerase fraction in this scenario does not depend on the effective nutrient quality $\bar{\nu}$, and therefore cannot depend on the growth rate; this behavior is consistent with the data of Balakrishnan et al. (10); to obtain $\epsilon_{max}$, we invert the equation in terms of $\phi_N^*$ and we use the empirical value of the observed RNA polymerase fraction, giving $\epsilon_{max} = 11184$;
4. To obtain $K_m$, we invert Eq. [S132] to get $K_m = \frac{[m^*]}{\epsilon\phi_N^*}$; we use $\epsilon$ with the functional form and parameters outlined above. We use $\phi_N^*$ extracted above (which is the empirical one by construction); we use the empirical mRNA concentrations for $[m]^*$ to obtain a curve $K_m$ vs $\lambda^*$, which is roughly a constant with some experimental fluctuations; finally, we take the average value of $K_m$ across the growth rates and we obtain $K_m = 0.16 \times 10^3/\mu m^3$.

### S18. Data Analysis: Comparison of the predicted response to transcription-targeting drugs to experimental data

This section specifies the procedure to obtain the model-data comparison shown in Fig. 4 of the main text. As outlined in the previous sections, we find the proteome composition that optimizes the growth rate from the following three equations,

$$\lambda = \gamma\phi_R\frac{\phi_N}{\epsilon^{-1}+\phi_N} \qquad \text{[S134]}$$

$$\gamma\phi_R\frac{\phi_N}{\epsilon^{-1}+\phi_N} = \nu\phi_C \qquad \text{[S135]}$$

$$\phi_R + \phi_N + \phi_C = 1 - \phi_Q\,. \qquad \text{[S136]}$$

We denote the growth-rate optimized proteome fractions as $\phi_R^*$, $\phi_N^*$, $\phi_C^*$, and $\lambda^*$. This solution depends on the parameter $\epsilon$, which we assume in our description to be the effective target of drugs that affect transcription. We define $\bar{\lambda} = \frac{\lambda}{\gamma}$, $\bar{\nu} = \frac{\nu}{\gamma}$, and $\phi^{max} = 1 - \phi_Q$, which are the rescaled growth rate, rescaled nutrient quality, and the maximum attainable protein fraction, respectively. As previously shown, the solution of the optimization problem is

$$\phi_R^* = \frac{\phi_{max}-\phi_N^*}{1+\frac{1}{\bar{\nu}}\frac{\phi_N^*}{\epsilon^{-1}+\phi_N}}\,, \qquad \phi_C^* = 1-\phi_Q-\phi_R^*-\phi_N^*\,, \qquad \bar{\lambda}^* = \frac{\bar{\nu}}{1+\bar{\nu}}\frac{\phi_N^*\left(\phi_{max}-\phi_N^*\right)}{\frac{\bar{\nu}}{1+\bar{\nu}}\,\epsilon^{-1}+\phi_N^*}\,. \qquad \text{[S137]}$$

To plot the solutions as a function of $\epsilon$, we must specify the values of $\bar{\nu}$, $\phi^{max}$, and $K_m$, which determines the total mRNA concentration $[m^*] = K_m\ \epsilon\ \phi_N^*$. Fig. 4 of the main text shows the solutions for three values of $\bar{\nu}$ (0.3, 1, and 2) corresponding to poor, intermediate, and rich nutrient conditions, respectively. We set $\phi^{max} = 0.45$ based on ref. (2) and $K_m = 0.16/\mu\text{m}^3$ using the value extracted from our analysis of ref. (10), as shown in Fig. 3 of the main text. We also set $\gamma = 8\ h^{-1}$, which is approximately the maximum translation elongation rate observed in ref. (26), to obtain the rescaled growth rate $\bar{\lambda}$ used in the equation, starting from the growth rate observed in ref. (2).

### S19. Table summarising the symbols used in the main text

**Table S1. Summary of the symbols used in the main text.**

| Definition and Symbol | Typical values *E. coli* (if present) | Typical values *S. cerevisiae* (if present) |
|---|---|---|
| growth rate $\lambda$ | 0.1-2 $\mathrm{h}^{-1}$ (26) | 0.1-0.5 $\mathrm{h}^{-1}$ (19) |
| total protein concentration $[P]$ | 2-3 $\mathrm{mM}$ (24) | 1-2 $\mathrm{mM}$ (24) |
| total mRNA concentration $[m]$ | 0.1-5 $\mu\mathrm{M}$ (10) | 0.1-1 $\mu\mathrm{M}$ (24) |
| translation flux per mRNA $J^{TL}$ | 100-1000 $\mathrm{h}^{-1}$ [a] | 100-1000 $\mathrm{h}^{-1}$ [b] |
| translation elongation rate $k_{tl}$ | 8-20 $\mathrm{aa/s}$ (26) | 10.5 $\mathrm{aa/s}$ (19) |
| transcription elongation rate $k_{tx}$ | 10-100 $\mathrm{nt/s}$ (24) | 40 $\mathrm{nt/s}$ (32) |
| total number of amino acids (aa) in a ribosome $L_R$ | $\sim$ 7300 (24) | $\sim$ 12500 (24) |
| total number of aa in an RNA polymerase $L_N$ [c] | $\sim$ 3600 (33) | $\sim$ 5000 (Pol II) (34) |
| inverse time to elongate a ribosome (translation) $\gamma = \frac{k_{tl}}{L_R}$ | 4-10 $\mathrm{h}^{-1}$ | 3 $\mathrm{h}^{-1}$ |
| inverse time to elongate an RNA polymerase $\gamma_{tx} = \frac{k_{tx}}{L_N}$ | 3-30 $\mathrm{h}^{-1}$ | 12 $\mathrm{h}^{-1}$ |
| average transcript degradation rate $d$ | 0.1-1 $\mathrm{min}^{-1}$ (24) | 0.02-0.2 $\mathrm{min}^{-1}$ (24) |
| fraction of DNA-bound RNA polymerases $f_{bn}$ | 0.9-1 (13) | - |
| effective mRNA concentration $K_m$ | 0.05-0.15 $\mu\mathrm{M}$ (see MM) | 0.25-0.25 $\mu\mathrm{M}$ (see MM) |
| mRNA demand-supply trade-off $\epsilon = \frac{\gamma_{tx} f_{bn}}{d} \frac{[P]}{K_m}$ | $10^2$-$10^5$ | $10^3\ \mathrm{f_{bn}}$-$10^7\ \mathrm{f_{bn}}$ |
| growth-independent protein fraction $\phi_Q$ | 0.55 (2) | - |
| ribosomal protein fraction $\phi_R$ | 0.05-0.2 (26) | 0.1-1 (19) |
| RNA polymerase protein fraction $\phi_N$ | 0.01 (10) | 0.01 (19) |
| catabolic protein fraction $\phi_C = 1 - \phi_Q - \phi_R - \phi_N$ | 0.2-0.4 | - |
| induced unneeded protein fraction $\phi_U$ | 0.1-0.4 (2) | 0.1-0.15 (17) |
| integrated unneeded gene copy number $g_U$ | - | 1-20 (17) |

[a] from previous estimates for $\lambda$, $[m]$, $[P]$ and using the formula $J^{TL} \approx \lambda \frac{[P]}{[m]}$ obtained by summing all equations [S2] and assuming exponential growth

[b] see previous footnote

[c] the total number of aa in an RNA polymerase was computed by taking the ratio of the RNA polymerase weight given by the references to the weight of a single amino acid. We have taken the latter weight to be 110 Da (24).

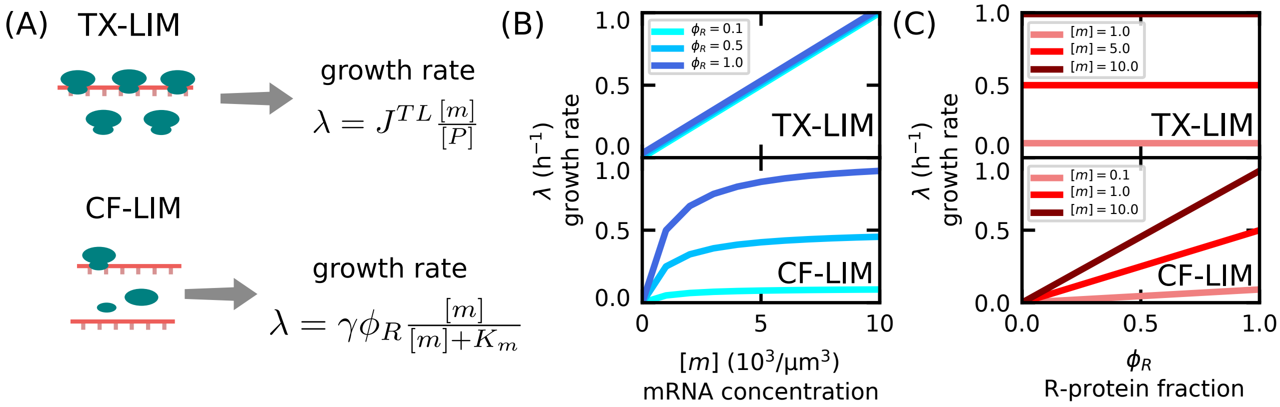


**Fig. S1.** The figure compares the growth laws that relate total mRNA, growth rate, and ribosome allocation in two different regimes: the transcription-limited (TX-LIM) and complex-formation limited (CF-LIM) regimes. The cartoon in panel A illustrates the differences between the two regimes: in TX-LIM ribosomes cannot bind to transcripts due to their saturation, while in CF-LIM increasing both mRNA and ribosome levels increases the formation of translationally active complexes. The two regimes predict different expressions for the growth rate, as shown on the right. Panels B anc C fix the parameters of the model to $J^{TL} = 1\mathrm{h}^{-1}$, $\gamma = 1\mathrm{h}^{-1}$, $[P] = 10 \cdot 10^3/\mu\mathrm{m}^3$ and $K_m = 1 \cdot 10^3/\mu\mathrm{m}^3$ to explore the main growth laws. (B) Growth rate dependence on total mRNA concentration at fixed R-protein fraction. Lighter to darker blue shades indicate increasing R-protein fractions. (C) Growth rate dependence on R-protein fraction at fixed total mRNA concentration (in units of $10^3/\mu\mathrm{m}^3$). Lighter to darker red shades indicate increasing total mRNA concentration. The two model regimes lead to qualitatively different predictions. See also Fig. 3 of the main text for the CF-LIM predictions with growth rate optimization and under increasing mRNA supply demand trade-off.

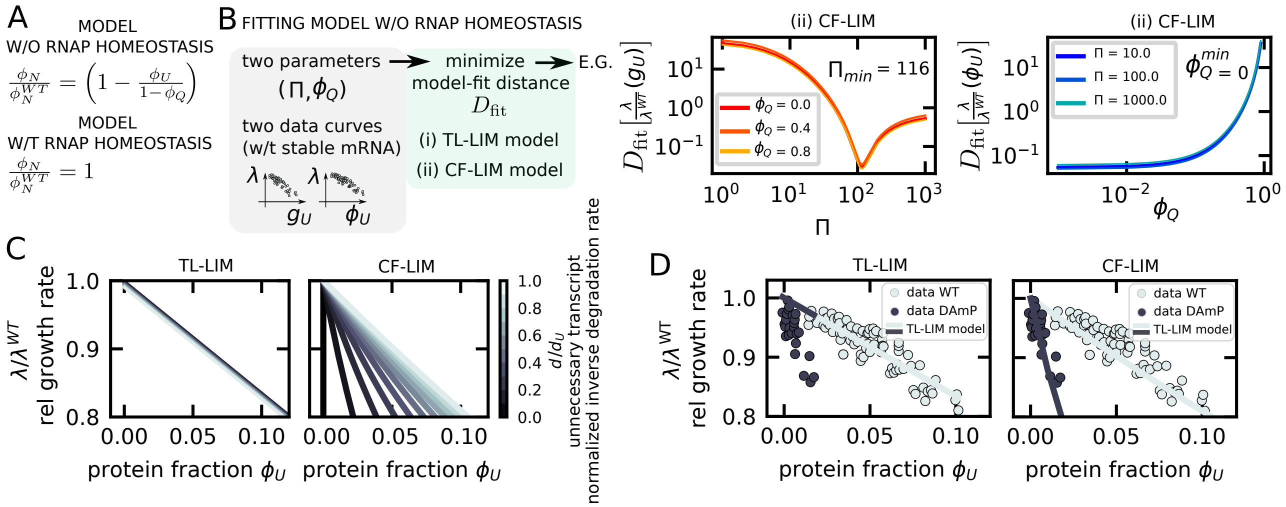


**Fig. S2.** Prediction of the growth cost of unneeded protein overexpression in a model without RNA polymerase homeostasis. (A) In the model variant without RNA polymerase homeostasis, the fraction of RNA polymerase proteins decreases with unneeded protein overexpression, which in turn decreases the mRNA concentration. Conversely, in the model variant presented in the main text with RNA polymerase homeostasis, the fraction of RNA polymerase proteins stays constant. (B) Parameter-estimation procedure of the model without RNA polymerase homeostasis. The model depends on two parameters, the inverse effective promoter strength $\Pi$ of the unneeded construct and the size $\phi_Q$ of the protein class $Q$ that is kept constant across growth perturbations. Experimental data from ref. (17) provide two curves of the relative growth rate, against the gene copy number and against unnecessary protein fraction. We find the pair of parameters that minimizes the residual of both plots (using a modified residual sum of squares). Note that we use only the curves of the stable construct to infer the parameters, in order to generate a true prediction for the DAmP construct. We perform the fit for both the translation-limited regime (TL-LIM) and the complex-formation-limited regime (CF-LIM). The right sub-panels illustrate the fitting procedure for the case of the CF-LIM model. In particular, note how each parameter only affects one curve and therefore can be extracted with a single-parameter fit. (C) The model predicts a drop in growth rate (quantified by the relative growth rate $\lambda/\lambda^{WT}$) as a function of protein fraction $\phi_U$ of the unnecessary proteins. The model also predicts how such trend changes as the degradation rate of the unnecessary transcript $d_U$ varies (through the ratio $d/d_U$, where $d$ is the average degradation rate of the other transcripts). The left sub-panel shows the TL-LIM prediction. The right sub-panel shows the CF-LIM prediction. (D) Data from ref. (17) falsify a scenario of translation-limited growth and validate the CF-LIM regime in *S. cerevisiae*. The left sub-panel shows the comparison between the data (circles) and a model of growth under pure translation limitation (solid lines). Light-grey circles represent data corresponding to stable transcripts ($d/d_U \approxeq 1$), and lines of the same color are model fits. Dark-grey circles represent data corresponding to unstable trascripts ($d/d_U \approxeq 0.08$), and dark-grey lines are model predictions. The right sub-panel shows that the prediction from the CF-LIM regime reproduces the data. All model curves in the panels are obtained with parameters $\Pi = 116$, $\phi_Q = 0$, $Z_g = Z_g^{WT} = 0$ (with $\Pi$ and $\phi_Q$ obtained from the fit in panel B, while $Z_g$ and $Z_g^{WT}$ are assumed to be zero).

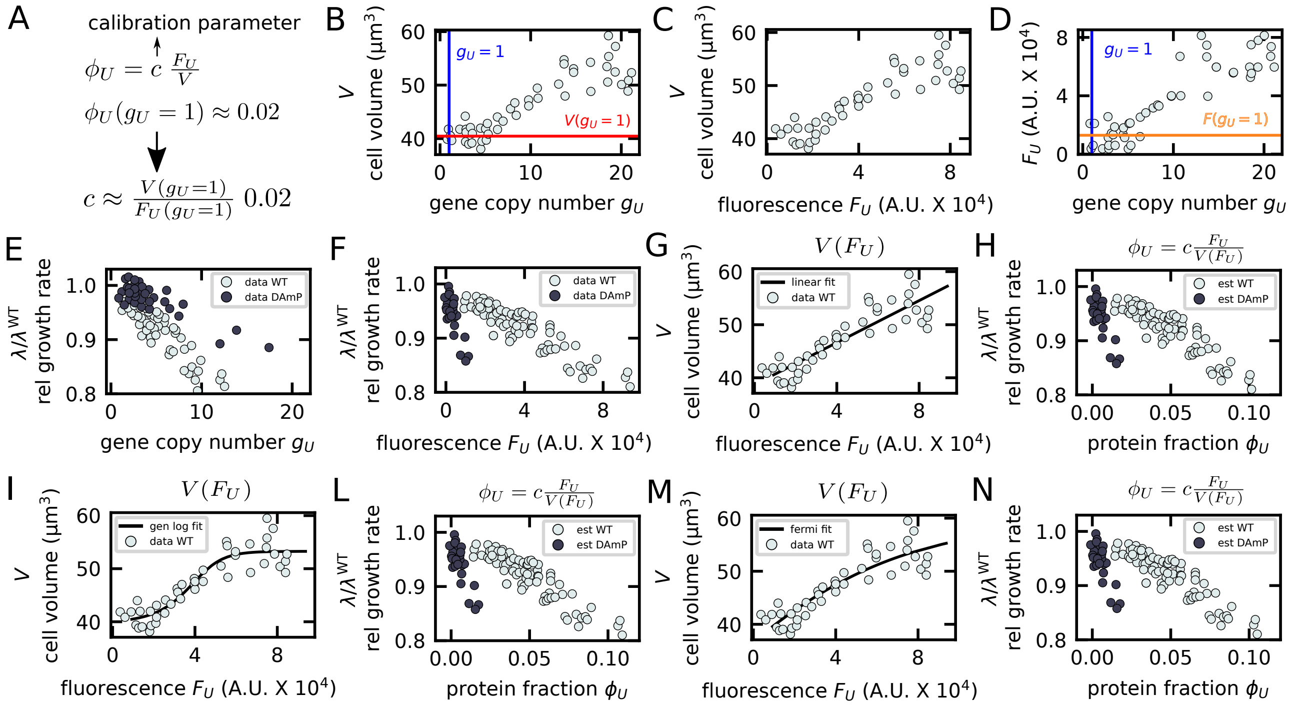


**Fig. S3.** Procedure to extract the dependence of the relative growth rate on the protein fraction of the unneeded proteins using *S. cerevisiae* data from ref. (17) (all circles in the plots refer to data from this study). (A) Relationship between unnecessary protein fraction $\phi_U$, fluorescence $F_U$ and volume $V$. In presence of a single unneeded construct ($g_U = 1$) ref. (17) find that the unneeded proteins occupy roughly $0.02\%$ of the proteome. By extracting $F_U(g_U = 1)$ and $V(g_U = 1)$ we obtain the parameter linking $\phi_U$ to $F_U$ and $V$. (B) Plot of measured cell volume against gene copy number of the unneeded stable construct. $V(g_U = 1)$ is obtained by averaging all data with $0.5 < g_U < 1.5$. Note that $g_U$ is not necessarily an integer in these data due to noisy estimation. (C) Plot of measured cell volume against gene copy number of the unneeded stable construct against the fluorescence of the unneeded proteins. (D) Plot of fluorescence against gene copy number of the unneeded stable construct obtained by associating each $g_U$ to a fluorescence value $F_U$ using panel (B) and (C). $F(g_U = 1)$ is obtained by averaging all data with $0.5 < g_U < 1.5$. (E) Plot of the relative growth rate against the gene copy number for the stable construct (data WT, light grey circles) and unstable construct (DAmP, dark grey circles). (F) Plot of the relative growth rate against the fluorescence for the stable construct (data WT) and unstable construct (DAmP). (G) Linear fit of the relationship between $V$ and $F_U$ for the unneeded stable construct. (H) Plot of the relative growth rate against the protein for the stable construct (data WT) and unstable construct (DAmP). The plot is obtained by using the relationship described in panel A in conjuction with the functional relationship $V(F_U)$ obtained though panel G (this is necessary because panels F and C do not come from the same experiment). Panels ILMN are identical to panels G and M, but using a different functional relationship between the volume and the fluorescence, respectively a generalized logistic and a Fermi function. Since the plots in panel (H), (L) and (N) are extremely similar, our procedure to extract the curves of the relative growth rate against the protein fraction does not depend on the precise form of the functional relationship.

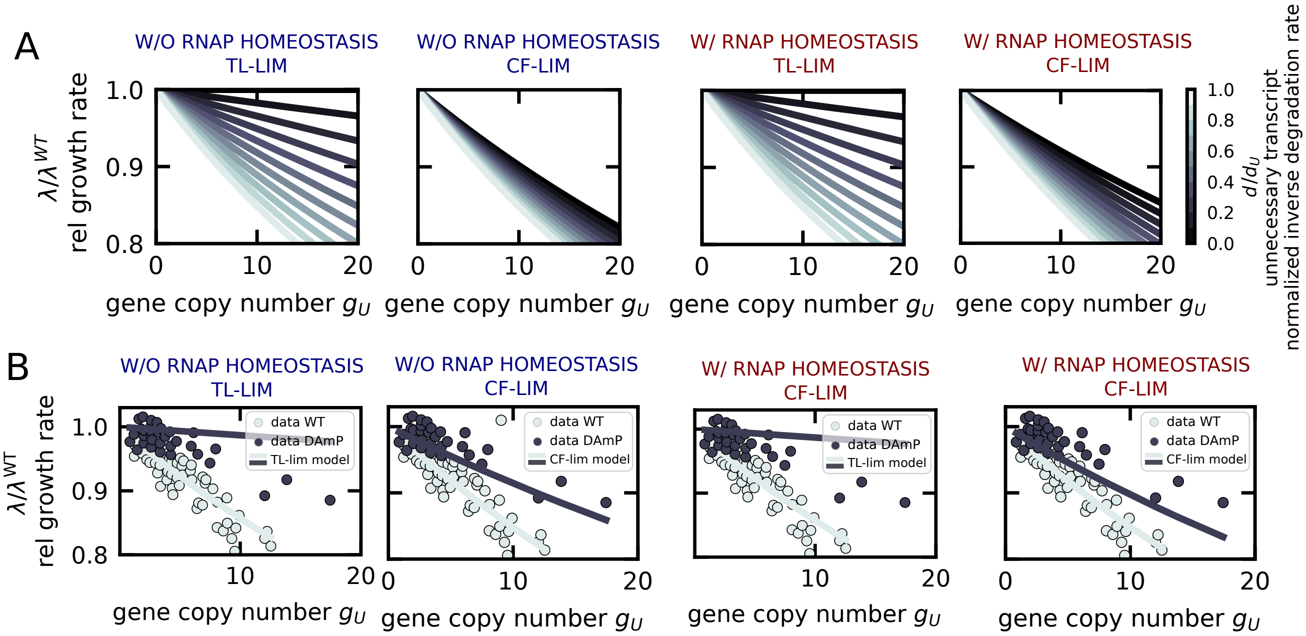


**Fig. S4.** Relative growth rate of unneeded protein overexpression as a function of gene copy number for the CF-LIM regime considered in this work, compared to the "usual" translation-limited TL-LIM regime. For these plots, the differences between the two scenarios are only quantitative. (A) Theoretical predictions for the relative growth rate against the gene copy number for different model variants and regimes. (B) Comparison with data from ref. (17) for different model variants and regimes. The best-fit scenario is CF-LIM without RNA polymerase homeostasis. The model parameters are fixed following the procedures described in this SI Appendix. All model curves in the panels without RNA polymerase homeostasis are obtained with parameters $\Pi = 116$, $\phi_Q = 0$, $Z_g = Z_g^{WT} = 0$. All model curves in the panels with RNA polymerase homeostasis are obtained with parameters $\Pi = 55$, $\phi_Q = 0.2$, $Z_g = Z_g^{WT} = 0$.

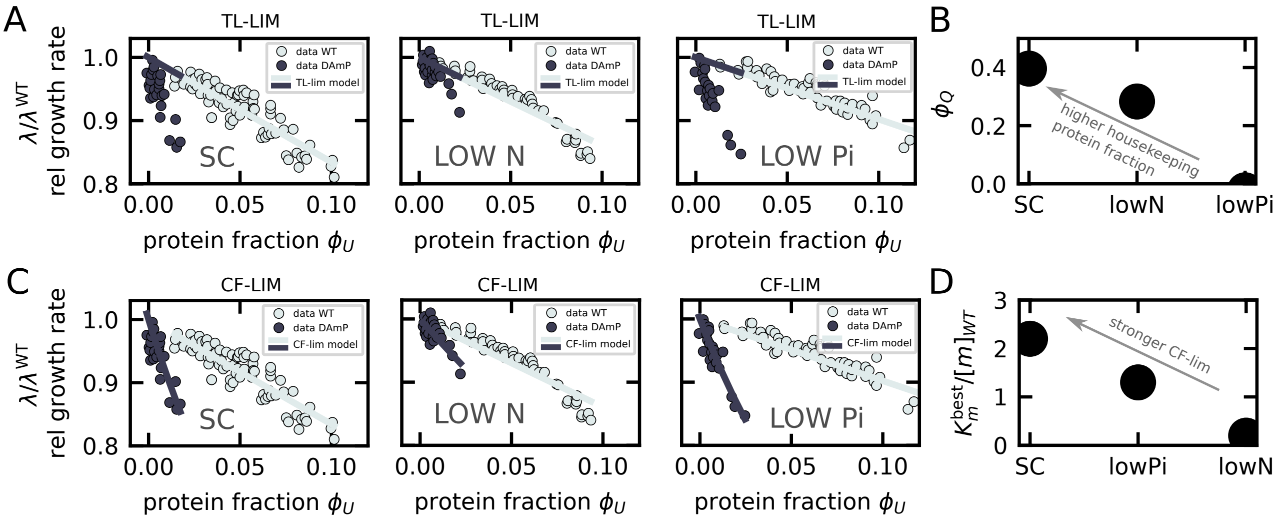


**Fig. S5.** Comparison of experimental data (yeast) and model predictions across the different conditions in ref. (17). The labels SC, Low N and Low Pi refer respectively to standard (Synthetic Complete), low nitrogen and low phosphate condition. The SC condition is also shown in Fig. 2 of the main text. (A) TL-LIM model predictions (solid lines) *vs* data (circles) across the different conditions in ref. (17). Model parameters were extracted from a fit of the WT data (light-grey circles), as described in sec. S16A, and the black solid lines are predictions for the DAmP data (black circles). (B) Parameter $\phi_Q$, the fraction of housekeeping proteins, across different conditions, from model fits assuming a TL-LIM model. The higher this parameter, the steeper the slope of the curve in panel A. (C) CF-LIM model predictions (solid lines) *vs* data (circles) across the different conditions in ref. (17). The CF-LIM model has fixed RNA polymerase fraction (RNAP homeostasis), as in Fig. 2 of the main text. The fitting is the one described in sec. S16B. All the parameters except $K_m$ are extracted from the WT curve. Note that the WT curve (assuming a model with RNAP homeostasis, $f_{bn} = 1$ and stable mRNA) does not depend on the parameter $K_m$, because total mRNA concentration is unchanged and consequently the term $\frac{[m]}{[m]+K_m}$ in the expression of the growth rate also does not change. Instead, we set $K_m$ by fitting the DAmP curve. (D) Fitted parameter $K_m^{\mathrm{best}}/[m]_{\mathrm{WT}}$ across different conditions, assuming a CF-LIM model. Higher values of this ratio imply stronger limitation for mRNA-ribosome complex formation. $[m]_{\mathrm{WT}}$ is the total mRNA concentration in the condition *without* overexpression burden (that is, $\phi_U = 0$). We note that the 'Low N' condition appears to be the one with the weakest limitation for complex formation, which is consistent with our microscopic interpretation of $K_m$ - this parameter is proportional to the translation elongation rate, which is affected negatively by low nitrogen. More surprisingly, the 'SC' condition lays deeper in the CF-LIM regime than the low phosphate condition 'low Pi' according to this fit, despite the fact the low phosphate should affect directly the production of mRNA, decreasing $[m]_{\mathrm{WT}}$ and increasing the ratio $K_m^{\mathrm{best}}/[m]_{\mathrm{WT}}$. One possibility is that the CF-lim regime is an incomplete model of the low phosphate condition, as low phosphate may affect mRNA levels so greatly that the system transits to a full transcriptional limitation, TX-LIM, regime (Fig. 1C of the main text). In that case, $K_m^{\mathrm{best}}/[m]_{\mathrm{WT}}$ is not a meaningful parameter to characterize the the overexpression cost. Low phosphate may also affect rRNA production, which may become the limiting process. Throughout the figure, we have assumed $f_{bn} = f_{bn}^{WT} = 1$, or equivalently $Z_g = Z_g^{WT} = 1$.

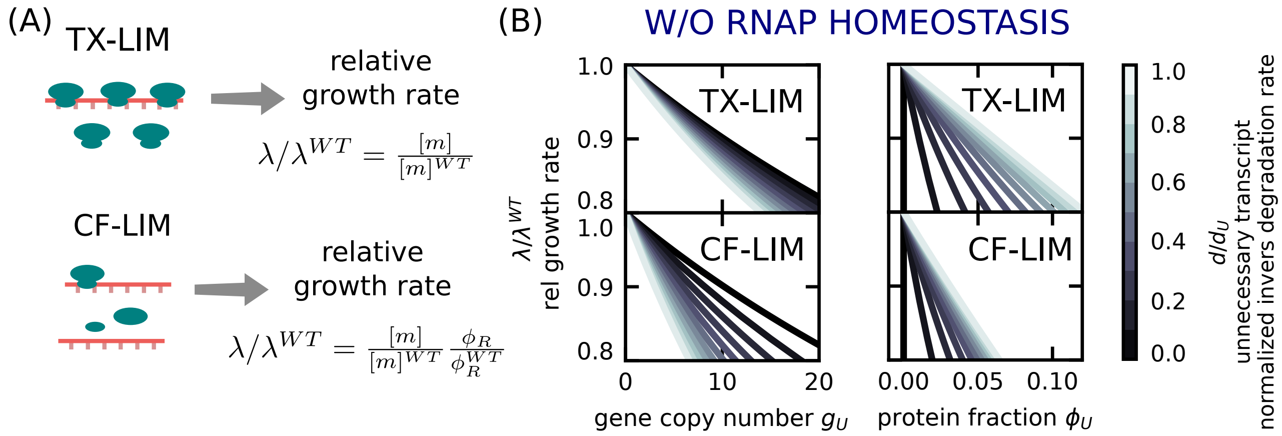


**Fig. S6.** Comparison of the cost of overexpression in the transcription-limited (TX-LIM) vs complex-formation limited regime (CF-LIM) under variable RNA polymerase protein levels (constant $\phi_N$). (A) Top-left: cartoon of the transcription-limited (TX-LIM) where ribosomes cannot bind to transcripts due to spatial saturation. Top-right: cost of protein overexpression as quantified by the relative growth rate $\lambda/\lambda^{WT}$. Bottom-left: cartoon of the transcription-limited (CF-LIM) where increasing both mRNA and ribosome levels increases binding rates. Bottom-right: cost of protein overexpression as quantified by the relative growth rate $\lambda/\lambda^{WT}$. (B) Cost of protein overexpression (quantified by the relative growth rate $\lambda/\lambda^{WT}$) as a function of gene copy number $\phi_U$ and protein fraction $\phi_U$. Lighter to darker shades of blue indicate the inverse degradation rate $\frac{d}{d_U}$ of the unnecessary transcript (or equivalently its stability life time) normalized to the average degradation rate of the other transcripts. Parameters for the plots in the TX-LIM regimes are $\Pi = 56$, $\phi_Q = 0.39$, $Z_g = Z_g^{WT} = 0$. Parameters for the plots in the CF-LIM regime are $\Pi = 116$, $\phi_Q = 0.2$, $Z_g = Z_g^{WT} = 0$ (from the fit in Fig. S2).

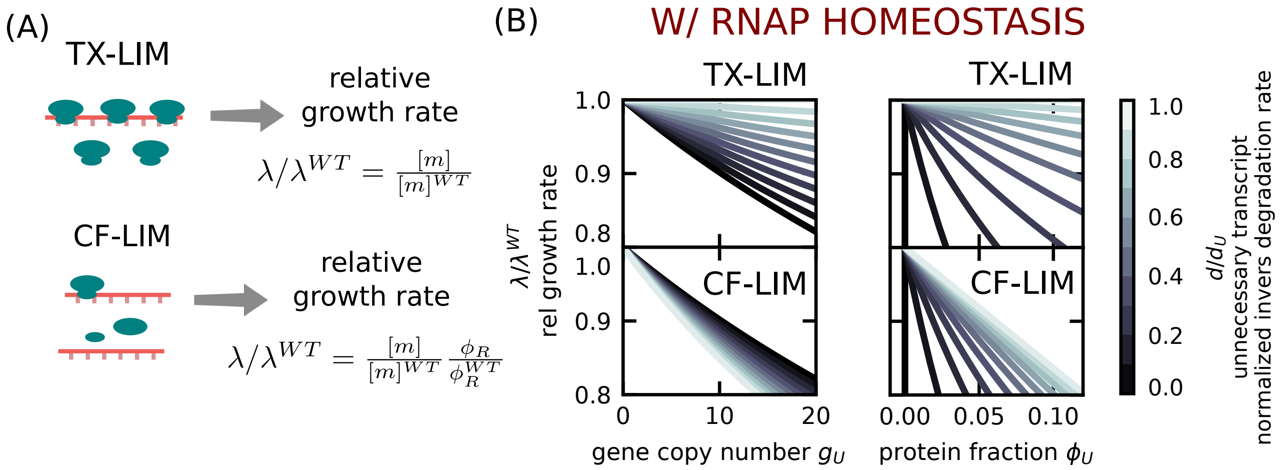


**Fig. S7.** Comparison of the cost of overexpression in the transcription-limited (TX-LIM) and the complex-formation limited regime (CF-LIM) under constant RNA polymerase protein levels (variable $\phi_N$). (A) Cartoon illustrating the differences between the transcription-limited (TX-LIM) regime, where ribosomes cannot bind to transcripts due to spatial saturation (top) and the CF-LIM regime, where increasing both mRNA and ribosome levels increases binding rates. The right side of the panel compares mathematically the cost of protein overexpression as quantified by the relative growth rate $\lambda/\lambda^{WT}$ in the two regimes. (B) Plots of the mathmeatical predictions for the cost of protein overexpression (quantified by the relative growth rate $\lambda/\lambda^{WT}$) as a function of gene copy number $\phi_U$ and protein fraction $\phi_U$. Lighter to darker shades of blue indicate the inverse degradation rate $\frac{d}{d_U}$ of the unnecessary transcript normalized to the average degradation rate of the other transcripts. The differences between the CF-LIM and TX-LIM predictions are quantitative only, but the TX-LIM scenario struggles to describe the unperturbed situation (see Fig. S1). The parameters for these plots (for both the TX-LIM and the CF-LIM regime) are $\Pi = 56$, $\phi_Q = 0.2$, $Z_g = Z_g^{WT} = 0$.

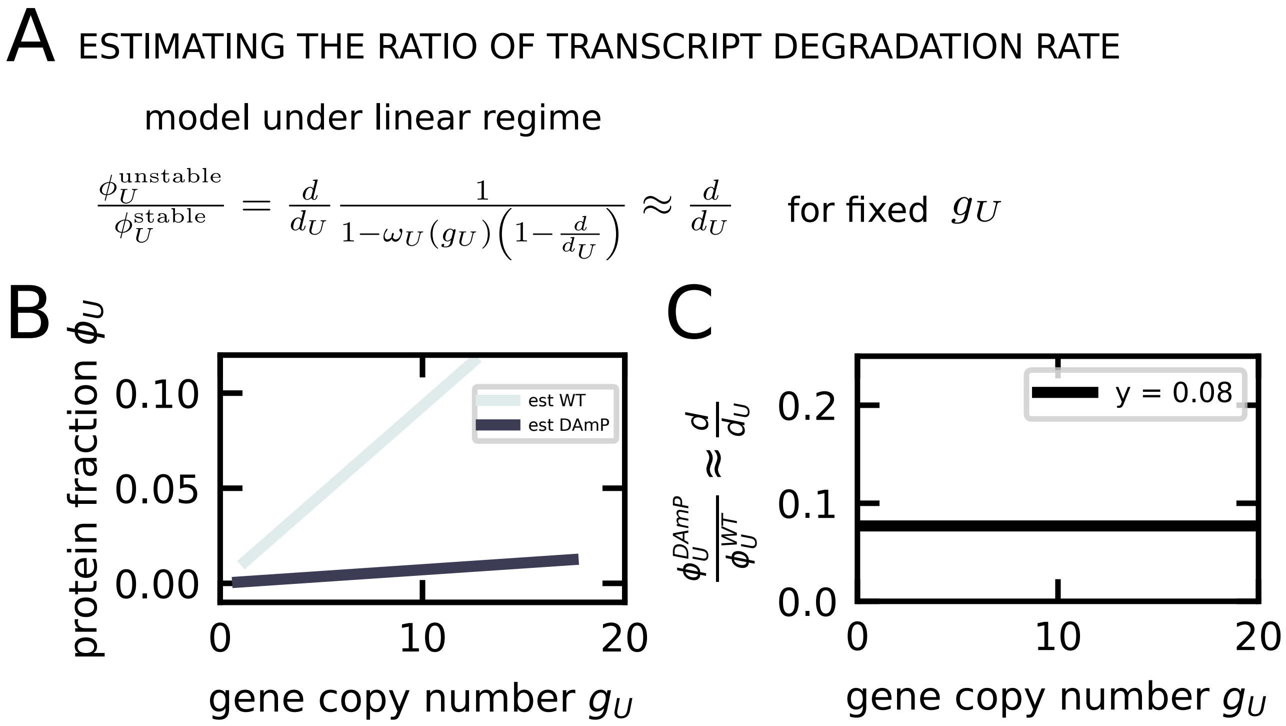


**Fig. S8.** Procedure to estimate the ratio of the transcript degradation rate from the two strains of *S. cerevisiae* obtained in ref. (17). (A) Illustration of the steps involved into estimating the degradation rate ratio. (B) The relationship between the protein fraction and the gene copy number for the wild-type and DAmP strain as estimated by the procedure outlined in the SI Appendix. (C) Final estimate of the ratio of the transcript degradation rates between the two strains, as obtained by dividing the two curves from panel (B). The parameters for plotting panel B are $\Pi = 56$, $\phi_Q = 0.2$, $Z_g = Z_g^{WT} = 0$. The data in panel (C) do not depend on these parameters (as they cancel out by taking the ratio).

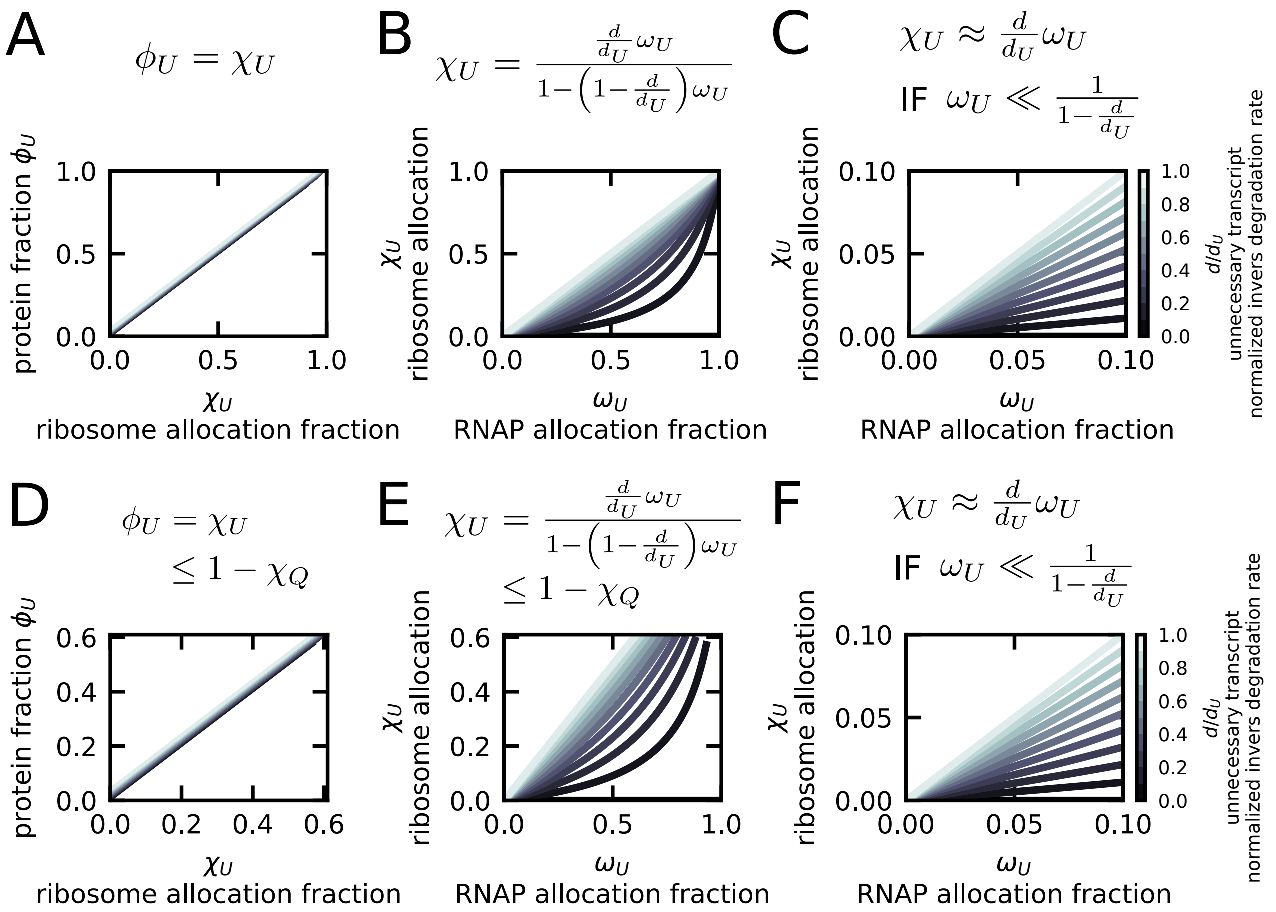


**Fig. S9.** The model predicts relationships between the protein fraction of the unneeded gene, the fraction of ribosomes translating the unneeded transcript (ribosome allocation fraction) and the fraction of RNA polymerases transcribing the unneeded genes (RNA polymerase allocation fraction). Panels (A), (B) and (C) show such relationships when the unneeded protein fraction can take any value between 0 and 1. (A) The protein fraction $\phi_U$ is always equal to the ribosome allocation fraction $\chi_U$ regardless of the transcript degradation rate (all curves collapse onto the same master curve). (B) The ribosome allocation fraction increases with the RNA polymerase allocation fraction with a rate that depends on the transcript degradation rate. Darker grey curves correspond to faster degradation rate, lighter grey curves to slower degradation rates (faster/slower compared to the average transcript degradation of the other transcript $d$). (C) Magnification of panel (B) in the region corresponding to experimental parameters. The plot shows that the ribosome allocation fraction scales linearly with the RNA polymerase allocation fraction in this regime, with a slope that corresponds to the inverse normalized transcript degradation rate. Panels (E), (D) and (F) show the same plot, but the unneeded protein fraction can only take any value between 0 and $1-\chi_Q$, where $\chi_Q$ is the size of the housekeeping proteome sector, kept constant across all growth perturbations, including forced overexpression of unneeded protein.

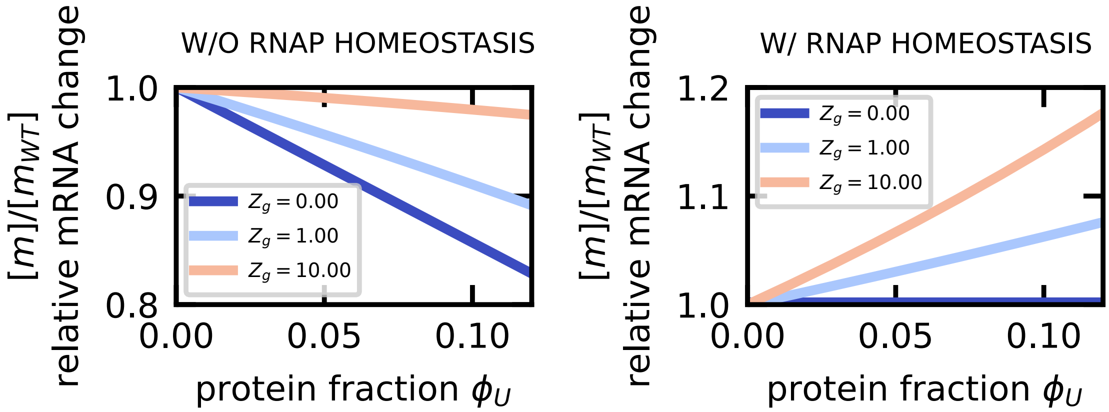


**Fig. S10.** Total mRNA can increase under unneeded protein over-expression due to free RNA polymerase recruitment to the genome. The plots show the relative decrease in the mRNA concentration $[m]/[m]_{WT}$ ($y$ axis) versus the fraction of unneeded protein $\phi_U$ in case RNA polymerase (RNAP) is not maintained homeostatically, or in other words $N$ is not in the Q sector (left), or in case RNA polymerase is maintained homeostatically, i.e. $N$ is in the Q sector (right). Different curves report different values of $Z_g$, which correspond to different values in the fraction of bound RNA polymerases $f_{bn}$ (greater $Z_g$ means lower $f_{bn}$). In the main regime considered in this study, $Z_g \approx 0$ and the fraction of specifically bound RNA polymerases $f_{bn} \approx 1$, and this effect is not observed. However, some studies have argued in favor of $f_{bn} \approx 0.5$, or $Z_g \approx 1$ (15, 16). All plots are obtained with $\phi_Q = 0.3$.

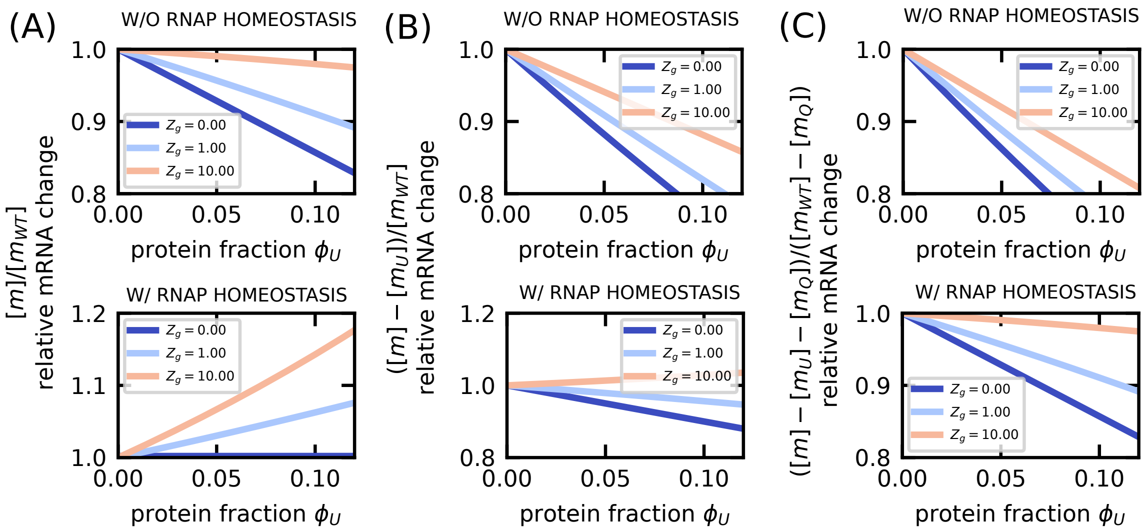


**Fig. S11.** Trend of total mRNA as a function of expression level of a useless protein under different definitions of total mRNA. (A) Relative change of total mRNA, including the mRNA of the unneeded gene, versus the fraction of unneeded protein $\phi_U$. (B) Relative change of total mRNA excluding unneeded mRNA versus the fraction of unneeded protein $\phi_U$. Note that the normalization remains the same because there are no unneeded proteins in the WT (un perturbed) condition. (C) The relative change of total mRNA excluding unneeded mRNA and Q-sector mRNA versus the fraction of unneeded protein $\phi_U$. Different curves report different values of $Z_g$, which correspond to different values in the fraction of bound RNA polymerases $f_{bn}$ (greater $Z_g$ means lower $f_{bn}$). All plots are obtained with $\phi_Q = 0.3$.

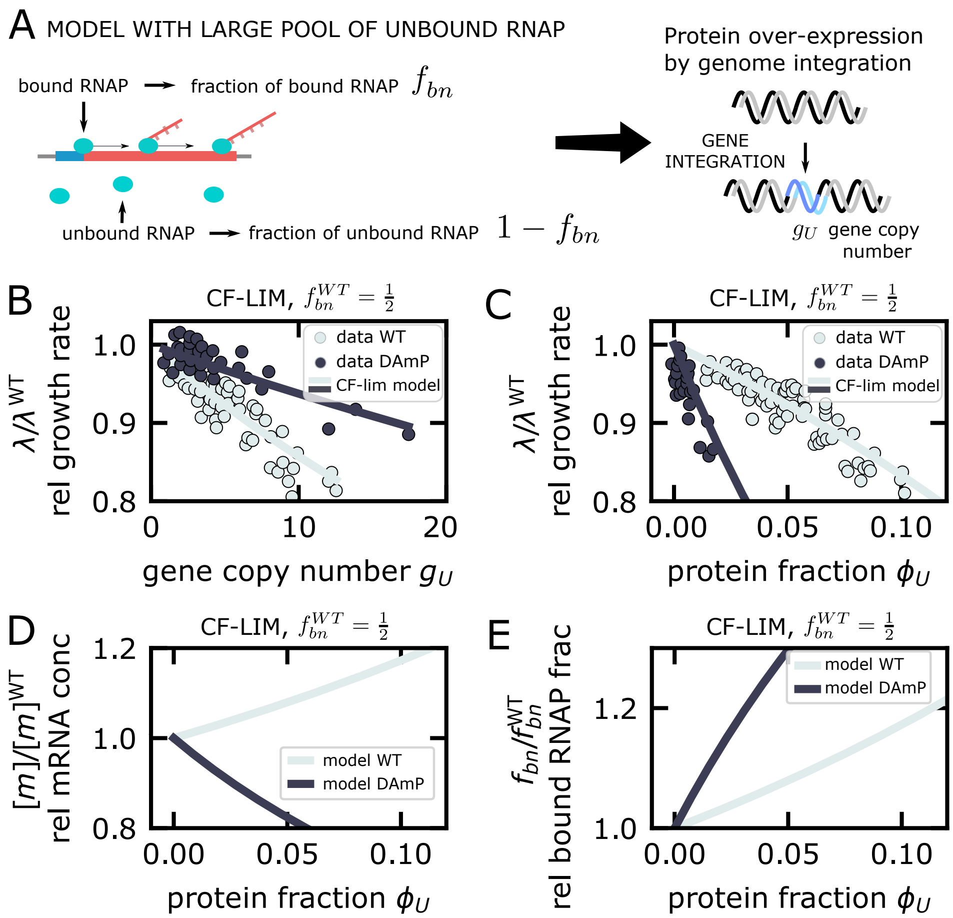


**Fig. S12.** Response to unnecessary protein expression in a model with a large pool of inactive RNA polymerase. (A) Illustration of the question: we ask how the presence of a significant pool of RNA polymerase that is not not actively transcribing affects the response to unnecessary protein expression. For simplicity our model distinguishes only two classes of RNA polymerase, actively transcribing or not (hence either unbound or non-specifically bound to DNA). The model predictions are given by Eq. [S79] and [S80], and show that RNA polymerase recruitment can increase the total mRNA concentration upon the perturbation. (B) Relative growth rate in presence of unneeded protein expression as a function of gene copy number $g_U$. The panel shows the comparison between data from ref. (17) (circles) and a CF-LIM model (solid lines) for stable (light grey symbols and lines) and unstable transcripts (dark grey symbols and lines). (C) Relative growth rate in presence of unneeded protein expression as a function of protein fraction $\phi_U$ in the same conditions. (D) Relative total mRNA concentration as a function of unneeded protein fraction $\phi_U$. The panel shows the prediction of the model for stable (light grey) and unstable transcripts (dark-grey). (E) Relative fraction of actively transcribing RNA polymerase as a function of unneeded protein fraction $\phi_U$. The panel shows the prediction of the model for stable (light grey) and unstable transcripts (dark grey). All model curves are obtained assuming RNA polymerase homeostasis (i.e., the complex belongs to the Q sector and its concentration does not change with growth perturbations). All model curves are obtained with parameters $\Pi = 50, \phi_Q = 0.66$, $Z_g = Z_g^{WT} = \frac{1}{1-\phi_Q}$. The value of $Z_g^{WT}$ are chosen so that in the unperturbed condition $f_b n^{WT} = 1/2$. As above, the other parameters are chosen in order to fit the WT points (light-grey circles), in a way that the DAmP data (dark-grey circles) are a prediction of the model.

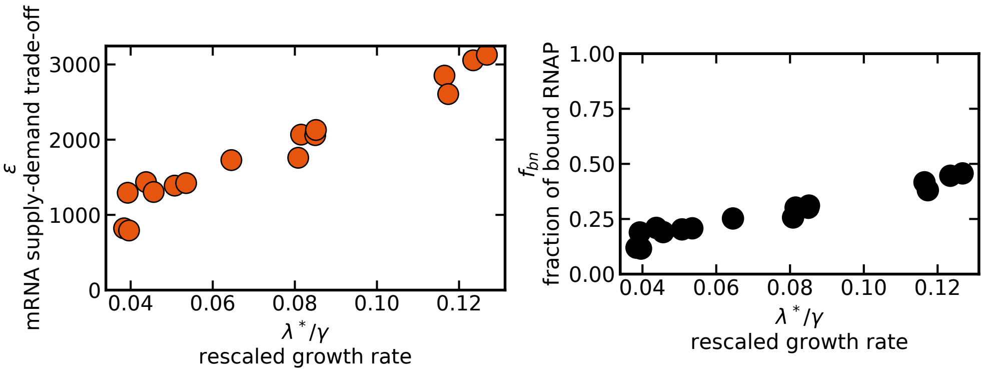


**Fig. S13.** (A) Estimate of the mRNA supply-demand trade-off parameter $\epsilon$ from the data of ref. (10) for *Escherichia coli*. We use $K_m = 0.16\ 10^3/\mu\mathrm{m}^3$ for our estimate (which is in accordance with mRNA concentration ranges measured by Balakrishnan and coworkers (10)), using the procedure outlined in section S17 of this SI Appendix. (B) Estimate of the fraction of bound RNA polymerase. We recall that in our model $f_{bn} = \frac{[m]}{[P]\phi_N}\ \frac{d}{\gamma_{tx}}$ or equivalently $f_{bn} = \epsilon \frac{K_m}{[P]}\ \frac{d}{\gamma_t x}$. We used this expression in conjuction with the mRNA data from ref. (10) to obtain $f_{bn}$. We used protein density $[P] = 3.15\ 10^6\ \mu\mathrm{m}^{-3}$ as indicated in ref. (10). For the average mRNA degradation rate $d$ and transcription elongation rate $\gamma_t x$, we took the geometric mean of the ranges identifies in table S1, that is, $d = \sqrt{3 \cdot 30}\ \mathrm{h}^{-1}$ and $\gamma_{tx} = \sqrt{0.1 \cdot 1}\ \mathrm{min}^{-1}$.

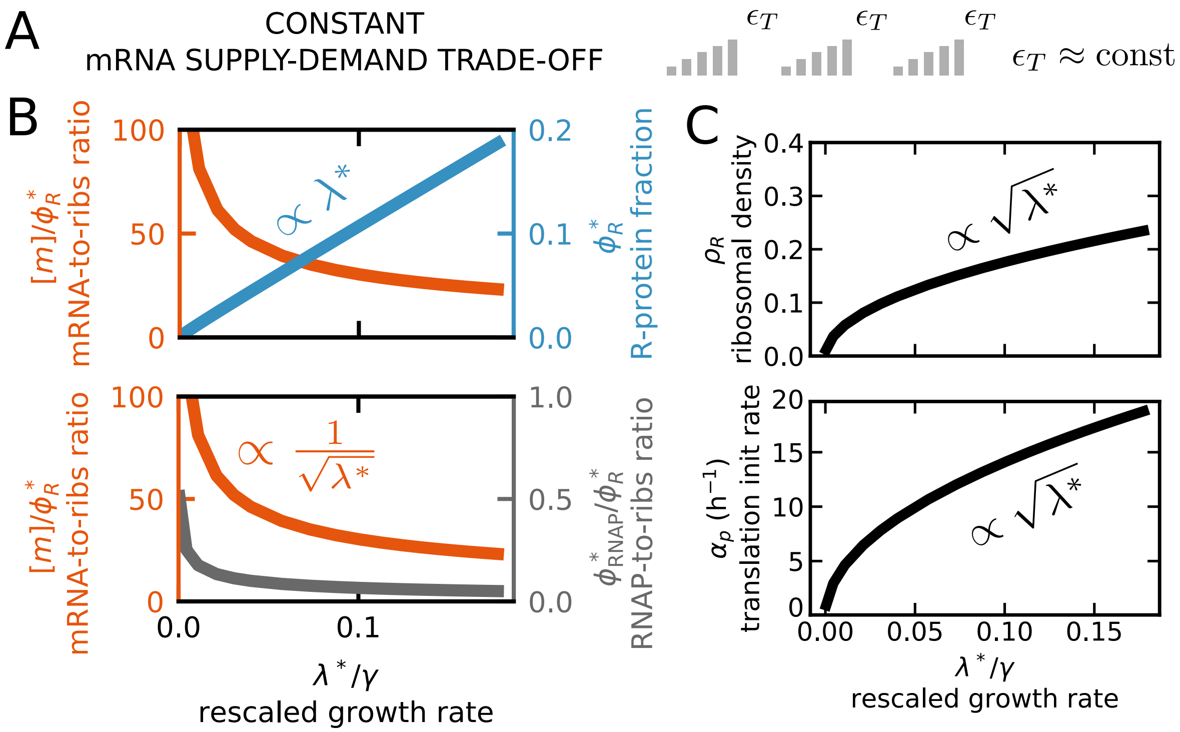


**Fig. S14.** The model predicts square-root growth laws between transcription and translation across growth conditions under constant mRNA supply-demand trade-off and growth-rate optimization. (A) Under costant mRNA supply-demand trade-off, the rate of mRNA production per RNA polymerase does not change across growth conditions. (B) The analytical prediction for the ratio of mRNA concentration to ribosomal protein fraction (orange line) decreases in this regime as the inverse of the square root of the growth rate, due to the square root increase of the RNA polymerase fraction (see main text) and the ribosomal fraction increasing linearly (blue line). The bottom sub-panel shows the analytical predictions for the optimal ratio of RNA polymerase to ribosomes (grey line). (C) The ribosomal density on mRNA increases as the square root of the growth rate (top panel) under this assumption. As the initiation rate and the ribosomal density are proportional to each other, the initiation rate is also predicted to increase with the square root of the growth rate (bottom panel). The plotted density is the ribosome coverage (linear) density on the mRNA, to obtain the linear number density one should divide these values by $\simeq 10$ (roughly the size of a ribosome in codon units). Across all panels, we employ the parameters described in section S17.

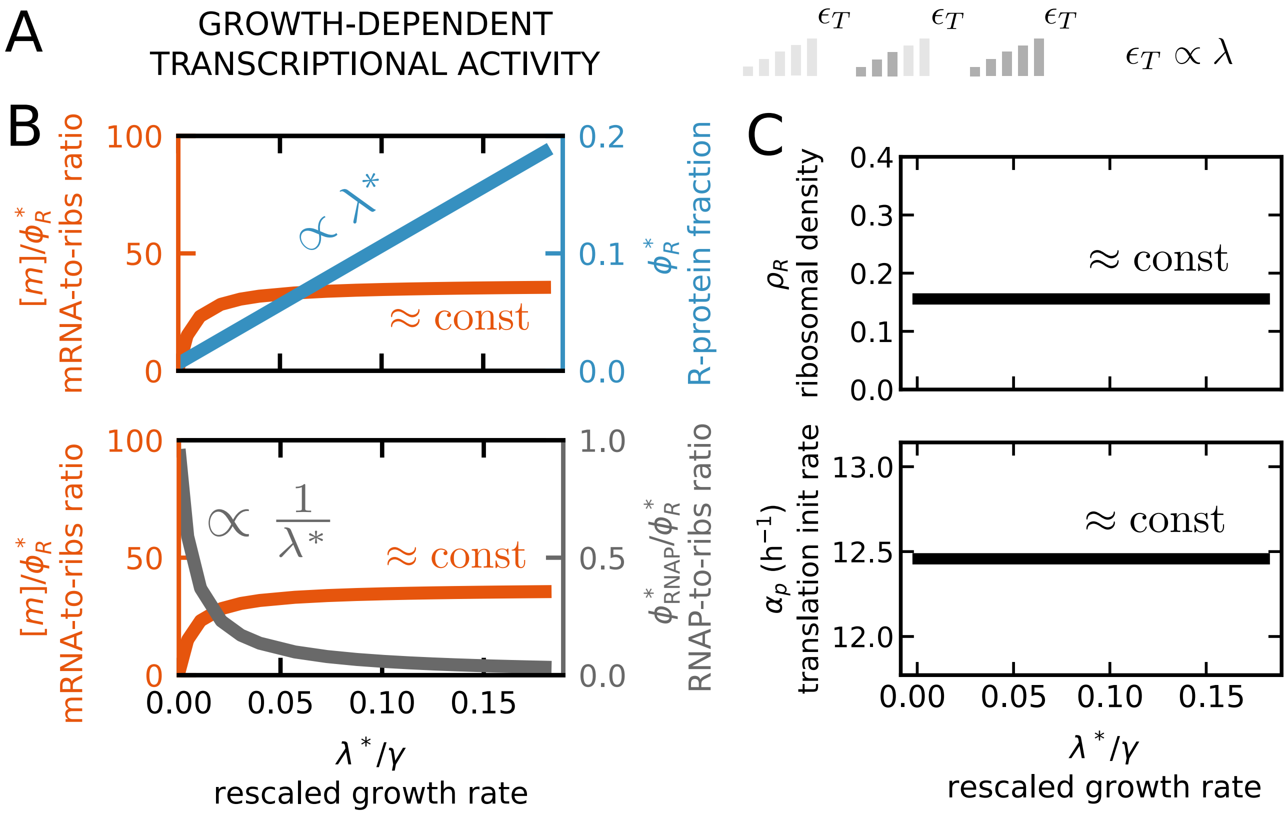


**Fig. S15.** The model predicts linear growth laws between transcription and translation across growth conditions under growth-dependent mRNA supply-demand trade-off and growth-rate optimization. (A) Under growth-dependent mRNA supply-demand trade-off, the rate of mRNA production per RNA polymerase increases linearly with the growth rate. (B) The predicted ratio of (optimal) mRNA concentration to ribosomal protein fraction (orange line) stays nearly constant with the growth rate due to the fact that both the transcriptional activity and ribosomal fraction (blue line) increase linearly with the growth rate. The bottom sub-panel shows the trend for the ratio of RNA polymerase to ribosomes (grey line). (C) The optimal ribosomal density on mRNA is predicted to stay constant with the growth rate under this assumption (top panel), in agreement with the hypothesis that a growth-dependent mRNA supply-demand trade-off parameter implements a constraint optimizing ribosome linear density on transcripts (10). Since the initiation rate and the ribosomal density are proportional to each other, the initiation rate also stays constant with the growth rate in this scenario (bottom panel). The plotted density is the ribosome coverage density on the mRNA, to obtain the number density one should divide these values by $\simeq 10$ (roughly the size of a ribosome in codon units). Across all panels, we employ the parameters described in section S17.

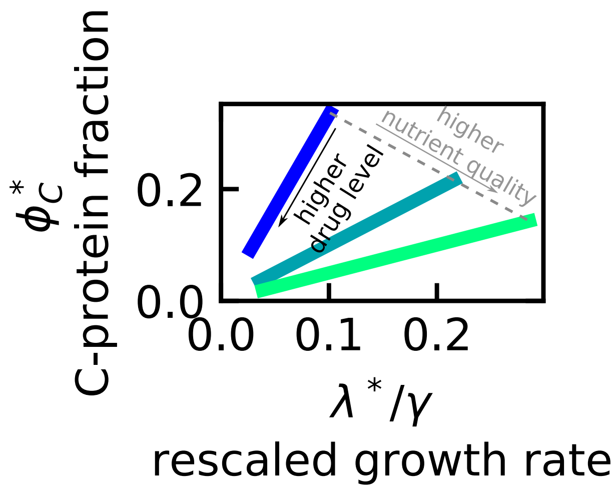


**Fig. S16.** The predicted catabolic fraction $\phi_C$ under growth optimization changes as transcription is inhibited (mRNA supply-demand trade-off parameter $\epsilon$ reduced). The predictions refer to the CF-LIM regime. Different solid lines indicate different nutrient conditions, with darker colors representing poorer media. The dashed line represents the first growth law. We set $\phi^{max} = 0.45$, $K_m = 0.16/\mu\text{m}^3$ and $\gamma = 8\ h^{-1}$, using ref. (2, 10) and our own analysis detailed in section S18.
